# Antimalarial Potential of Synthetic Geraniol and Nerol Analogs

**DOI:** 10.1101/2025.08.01.664114

**Authors:** Maurício Mazzine Filho, Isaac Agnelo da Silva Sobreiro, Gabriel dos Santos e Silva, Carlos Alberto de Araújo Silva, Leonardo Vinícius Nogueira Lima, Arthur Vicentini Furtado, João Vitor da Silveira, Marcus Vinícius Nora de Souza, Matheus Felipe Silva Santos, Gabriela Oliveira Castro, Manoel Aparecido Peres, Marcell Crispim, Alejandro Miguel Katzin, Mauricio Frota Saraiva, Ignasi Bofill Verdaguer

## Abstract

Drug resistance is a major threat to malaria control, and thus new drugs are required to fight against this parasitosis. To accelerate drug development, it is of special interest the exploration of natural compounds or repositioning drugs already employed for other diseases. Considering this, previous studies have found that diverse plant terpenes arrest *Plasmodium* parasites *in vitro* and *in vivo* models. However, most terpenes possess low toxicity for the parasite and/or face pharmacokinetic issues. Here, we report several new acyclic monoterpene analogs which possess great antiplasmodial activity *in vitro* (50% inhibitory concentration at low micromolar scale) against *P. falciparum* parasites. Also, *in vitro* studies using hepatocellular carcinoma cells (HepG2) demonstrated remarkable selectivity for malaria parasites. Furthermore, bioinformatic approaches revealed that these compounds possess acceptable pharmacological properties. All these results suggest that acyclic monoterpene analogues could serve as a promising starting point for the development of synthetic terpenes as antimalarial drugs.

## 1 INTRODUCTION

*Plasmodium falciparum* is the causative agent of human malaria. In 2022, the World Health Organization reported an estimated 249 million cases of malaria and 608,000 malaria-related deaths^[1]^. Resistance to current antimalarial drugs is arguably the most significant challenge to controlling malaria^[1]^. Therefore, the identification and development of novel antimalarial therapies are urgently needed. To accelerate drug development, there is significant interest in exploring natural compounds or their analogues^[2,3]^.

One of the most studied metabolic pathways in the malaria parasite is the isoprenoid biosynthesis via the methylerythritol phosphate (MEP) pathway^[4]^. The MEP pathway leads the biosynthesis of farnesyl pyrophosphate (FPP, 15 carbon), and geranylgeranyl pyrophosphate (GGPP, 20 carbon)^[5,6]^. These metabolites are essential for the farnesylation and geranylgeranylation of proteins, as well as the biosynthesis of ubiquinone or dolichol^7–9]^. Due to their chemical structures being similar to isoprenoid intermediates, our group explored the antiplasmodial activity and metabolic effects of several plant terpenes, including nerolidol, perillyl alcohol, limonene, and linalool^[10–14]^. Most of these terpenes inhibit the *in vitro* growth of *P. falciparum* parasites. *In vivo*, perillyl alcohol and nerolidol administered via the inhalatory route demonstrated some antimalarial activity in animal models^[12,14]^. Nerolidol, limonene, and linalool inhibited dolichol biosynthesis and protein prenylation; nerolidol and linalool also inhibited the biosynthesis of the isoprenic side chain of ubiquinones^[11]^. However, most of the mentioned terpenes only exhibit toxicity for the parasite at millimolar concentrations (e.g., linalool and limonene), which are unlikely to be achieved in a clinical setting. Furthermore, these compounds have not yet been used in clinical settings for any pathology, probably due to their low bioavailability via preferred administration routes, such as oral administration^[12,14]^. In the previous work^[15]^ we described the synthesis and antitrypanosoma activity of GIB24 a geranyl-diamine that showed promising antiparasitic activity. To explore this compound as a potential agent to other parasitic diseases and inspired by the potential of natural terpenes to act in the MEP pathway of *P. falciparum* we investigated the antiplasmodial activity of GIB24 and fourteen others acyclic monoterpenes, analogous to nerol or geraniol. As demonstrated below, these compounds exhibited promising antiplasmodial activity and interesting selectivity to the parasite.

## 2 RESULTS AND DISCUSSION

### 2.1 Chemistry

#### 2.1.1 Preparation of nerol and geraniol analogues 3a-h; 4a-e and 6a-b

The synthesis of the proposed analogs was initiated by treating terpene alcohols, nerol (**1**) or geraniol (**2**), with phosphorus tribromide (PBr_3_) at negative temperatures in anhydrous tetrahydrofuran (THF). The formation of the respective terpenyl bromides **3** and **4** were monitored using thin-layer chromatography (TLC) and infrared (IR) spectroscopy. In the IR spectra of geraniol and nerol bromides, the band of O-H stretching (approximately 3300 cm⁻¹) was not observed, indicating the occurrence of the reaction. In the TLC a spot with high retention factor (rf) was observed. Subsequently, the bromides reacted with excess of various nucleophiles at room temperature to yield the desired compounds **3a-h**, with yields in the range of 20-65% and **4a**-**e** (yields 34-85%). In the IR spectra of geranyl or neryl analogues **3a-d**, **3g**, **4a**-**b**, and **4d** was observed the typical broadband of O-H stretching between 3291-3394 cm⁻¹, indicating the occurrence of the reaction. For the analogues **3e**, **3f**, and **3h** without hydroxyl group, the occurrence of reaction was verified by the presence in the IR spectra of N-H stretching between 3301-3352 cm⁻¹, indicating the occurrence of the reaction.

The neryl aminoalcohols **3a** and **3b** were subjected to mesylation reactions by treatment with methanesulfonyl chloride (MsCl) and triethylamine (Et_3_N) in anhydrous dichloromethane (DCM). The reactions were initiated at low temperature (0°C), and after the addition of the reagents, the system was allowed to warm to room temperature for 12 hours, yielding the mesylated intermediates **5a**-**b**. The formation of compounds **5a**-**b** was monitored using thin-layer chromatography (TLC) and infrared (IR) spectroscopy, and the intermediates were used immediately after their preparation. The final compounds (**6a**-**b**) were obtained by reacting to the mesylated intermediates **5a**-**b** with ethanolamine at room temperature. The compounds **6a** and **6b** were obtained with yields of 45 and 29% respectively. In the IR spectra of neryl analogues **6a** and **6b** was observed the typical broadband of O-H stretching between 3351-3362 cm⁻¹, indicating the occurrence of the reaction. In the ^13^C NMR spectra of these compounds the signal of both, terpene (C=C; 151.5-109.9 ppm) and the aminoalcohol, diamine or thioamine groups (65.2-8.3 ppm) were observed, indicating the final product formation. Moreover, the observed m/z values in mass spectra agree with the calculated values (see supplementary material). The reactions outlined are shown in **Figure 1**.

**Figure 1:**
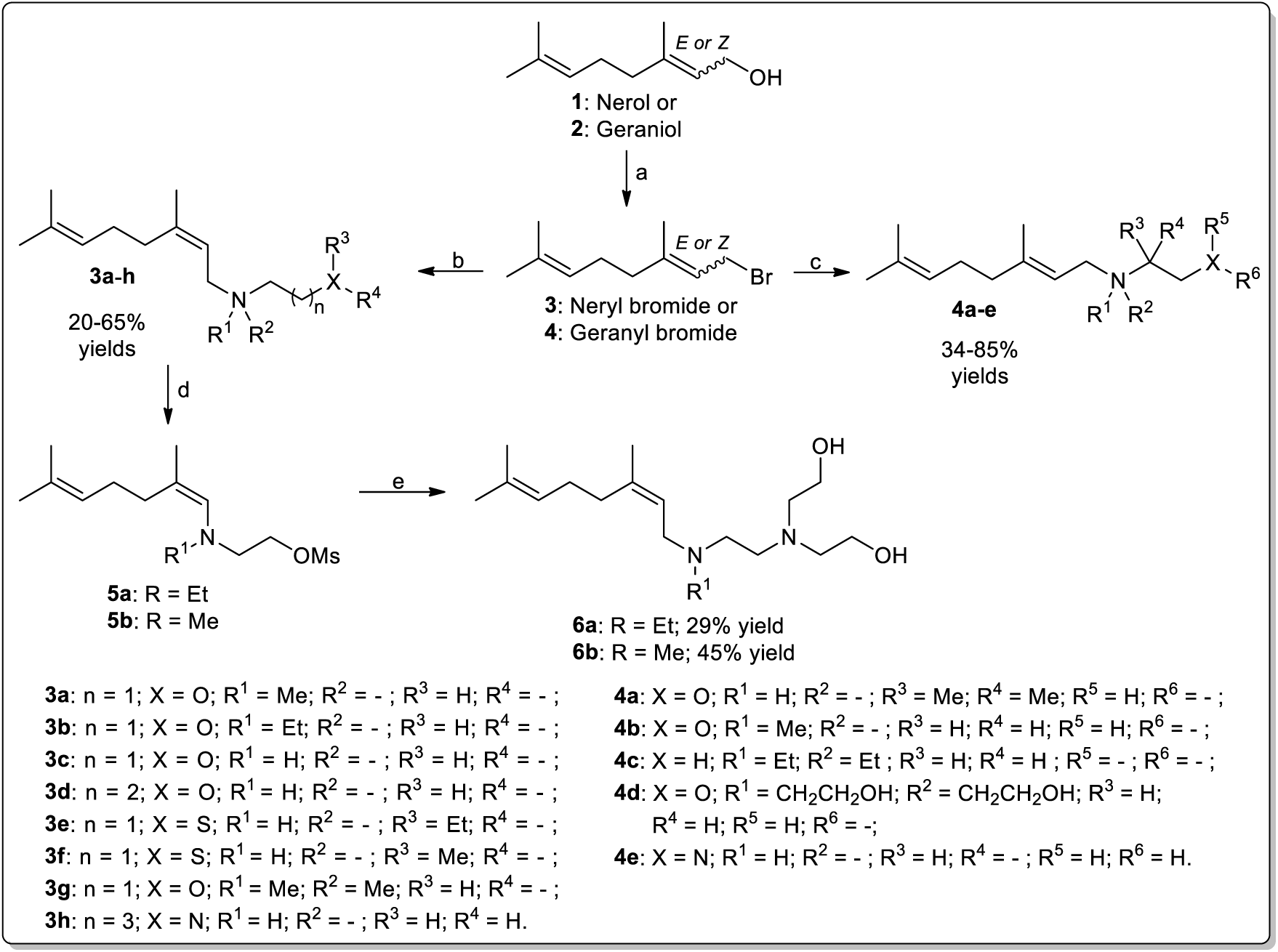
Reagents and conditions: (a) PBr_3_, anhydrous THF, (-18°C for geraniol, 40 min.) or (-35°C for nerol, 90 min); (b and c) methylaminoethanol (**3a** and **4b**), 2-(ethylamino)ethanol (**3b**), ethanolamine (**3c**), propanolamine (**3d**), 2- (ethylthio)ethylamine (**3e**), 2-(methylthio)ethylamine (**3f**), 2-(dimethylamino)ethanol (**3g**), 1,4-diaminobutane (**3h**), 2-amino-2- methyl-1-propanol (**4a**), triethylamine (**4c**), triethanolamine (**4d**), ethane-1,2-diamine (**4e**)^[15]^, dry DCM, r.t, 24h; (d) MsCl, dry DCM, 0 °C – r.t, 12h; (e) diethanolamine, DMF, r.t, 18h.

### 2.2 Biological assays

#### 2.2.1 Effects on *Plasmodium falciparum* growth and HepG2 selectivity

The growth of *P. falciparum* in the presence of various concentrations of acyclic monoterpene analogues was monitored to obtain the dose-response curve and calculate the IC_50_ values (Table 1). The results showed that all acyclic monoterpene analogues followed a sigmoid dose-response curve and exhibited diverse IC_50_ values. The three most potent compounds were **4e (GIB24)** with 8.94 ± 1.31 µM, **3e** with 7.49 ± 1.43 µM, and **3f** with 3.14 ± 1.18 µM. In contrast, the least potent compounds were **4b** with 121.10 ± 20.96 µM, **4a** with 133.23 ± 14.25 µM, and **6b** with 190.63 ± 33.95 µM. Despite the high IC_50_ values for (**4b**, **4a**, and **6b**) all synthesized compounds were more active than their respective natural terpene.

**Table 1.** *In vitro* antimalarial and cytotoxic activities of acyclic monoterpene analogs. The figure shows dose-response curves of *P. falciparum* proliferation and HepG2 cell viability after 48 hours of exposure to acyclic monoterpene analogs. The table presents the IC_50_ values (mean ± SD from three independent experiments) for *P. falciparum* and HepG2 cells. Additionally, the table includes the Selectivity Index (SI) of each compound. * Indicates that the SI values for nerol and geraniol. Values were estimated using the maximum evaluated concentration tested against HepG2: >1000 µM for nerol and >400 µM for geraniol as reported by Polo et al., 2011^[16]^ and Silva et al., 2021^[17]^, respectively.

| Compound | IC<sub>50</sub> (μM) | CC<sub>50</sub> (μM) | Selectivity index |
| --- | --- | --- | --- |
| **3a** | 48.90 ± 7.15 | 604.83 ± 28.88 | 12.37 |
| **3b** | 41.31 ± 3.08 | 430.37 ± 111.06 | 10.42 |
| **3c** | 49.71 ± 2.70 | 368.17 ± 30.28 | 7.41 |
| **3d** | 36.65 ± 5.60 | 458.00 ± 205.80 | 12.50 |
| **3e** | 7.49 ± 1.43 | 86.51 ± 29.13 | 11.55 |
| **3f** | 3.14 ± 1.18 | 101.99 ± 14.62 | 32.49 |
| **3g** | 23.09 ± 3.74 | >1000 | >43.31 |
| **3h** | 10.14 ± 0.99 | 62.58 ± 28.65 | 6.17 |
| **4a** | 133.23 ± 14.25 | 388.53 ± 7.10 | 2.92 |
| **4b** | 121.10 ± 20.96 | >500 | >4.13 |
| **4c** | 14.06 ± 1.05 | 115.43 ± 11.96 | 8.21 |
| **4d** | 70.16 ± 5.78 | >1000 | >14.25 |
| **4e** | 8.94 ± 1.31 | 34.35 ± 2.30 | 3.84 |
| **6a** | 91.00 ± 15.36 | 742.50 ± 230.87 | 8.16 |
| **6b** | 190.63 ± 33.95 | >1000 | >5.25 |
| Geraniol | 190.90 ± 1.038 | >400 | >2.09* |
| Nerol | 212.53 ± 57.558 | >1000 | >4.71* |

**Table 2.** ADME-Tox study of geraniol and nerol analogs. . The figure presents the ADME-Tox parameters of the compounds analyzed in this study. ADME-Tox analysis predicts compound behavior and toxicity through several parameters. TPSA (Topological Polar Surface Area) indicates cell permeability, where values <140 suggest high permeability, while higher values imply reduced absorption. WLOGP assesses lipophilicity, with values between 1 and 3 indicating an optimal balance for solubility and permeability. GI Absorption evaluates gastrointestinal absorption as high, moderate, or low, with high absorption preferred for oral administration. BBB Permeant determines the ability to cross the blood-brain barrier, with“Yes” suggesting potential central nervous system activity. PGP Substrate identifies substrates of P-glycoprotein, where “Yes” may reduce bioavailability due to cellular efflux. CL-Plasma (Plasma Clearance) measures the rate of compound removal from plasma, with lower values indicating prolonged retention. T1/2 (Half-life) reflects the duration of therapeutic action, with higher values suggesting longer action. AMES evaluates mutagenic potential (positive indicating genotoxicity, negative indicating safety), and Carcinogenicity assesses cancer risk, where values between 0-0.3 indicates low probability of being carcinogenic; values between 0.3-0.7 indicates medium probability of being carcinogenic; and values between 0.7-1.0 indicate high probability of being carcinogenic. These parameters guide the selection of safe and effective drug candidates. Computational modeling was performed using the SwissADME and ADMETlab 3.0 platforms^[18]^.

| Compound | TPSA | WLOGP | GI absorption | BBB permeant | Pgp substrate | Caco-2 permeability | cl-plasma | t1/2 | Ames | Carcinogenicity |
| --- | --- | --- | --- | --- | --- | --- | --- | --- | --- | --- |
| **3a** | 23.47 | 2.6 | High | Yes | No | -4.65 | 12.63 | 1.7 | 0.19 | 0.331 |
| **3b** | 23.47 | 2.99 | High | Yes | No | -4.68 | 11.66 | 1.48 | 0.197 | 0.307 |
| 3c | 32.26 | 2.26 | High | Yes | No | -4.87 | 10.52 | 1.37 | 0.144 | 0.097 |
| 3d | 32.26 | 2.65 | High | Yes | No | -4.86 | 12.39 | 1.5 | 0.133 | 0.062 |
| 3e | 37.33 | 4.02 | High | Yes | No | -4.97 | 10.09 | 0.88 | 0.307 | 0.19 |
| 3f | 37.33 | 3.63 | High | Yes | No | -4.77 | 11.46 | 1.26 | 0.374 | 0.154 |
| 3g | 20.23 | -0.25 | Low | No | Yes | -5.06 | 11.86 | 1.52 | 0.101 | 0.318 |
| 3h | 38.05 | 3.01 | High | Yes | No | -5.05 | 9.57 | 1.12 | 0.224 | 0.055 |
| 4a | 20.23 | 3.05 | High | Yes | Yes | -4.65 | 10.5 | 1.48 | 0.322 | 0.52 |
| 4b | 23.47 | 2.68 | High | Yes | No | -5.14 | 9.96 | 0.97 | 0.323 | 0.336 |
| 4c | 0.0 | 1.56 | Low | No | Yes | -5.16 | 12.2 | 1.52 | 0.201 | 0.231 |
| 4d | 60.69 | -1.44 | High | No | Yes | -5.14 | 5.87 | 1.08 | 0.023 | 0.11 |
| 4e | 38.05 | 2.23 | High | Yes | No | -5.28 | 10.18 | 1.27 | 0.305 | 0.167 |
| 6a | 46.94 | 2.25 | High | Yes | No | -4.83 | 10.33 | 1.21 | 0.292 | 0.131 |
| 6b | 46.94 | 1.9 | High | Yes | No | -5.24 | 9.76 | 1.26 | 0.296 | 0.091 |

Subsequently, the CC_50_ values of these compounds were determined in HepG2 cells. The most cytotoxic compounds were **3e** with 86.51 ± 29.13µM, **3h** with 62.58 ± 28.65 µM and **4e** with 34.35 ± 2.30, while the least cytotoxic were **3d** with 458.00 ± 205.80 µM, **3a** with 604.83 ± 28.88 and **6a** with 742.50 ± 230.87 µM. Notably, in this assay, a maximum concentration of 1 mM was tested, and some compounds did not result in a complete loss of viability. For compounds **4b**, **3g**, **6b**, and **4d**, it was not possible to calculate the CC_50_ value, with GraphPad Prism analysis estimating it to be >500 for the first and >1000 for the other three, respectively.

Nonetheless, the Selectivity Index (SI) was calculated either as a defined ratio or as a minimum value when CC_50_ values were not obtained. The most selective compounds for the malaria parasite were **4d** with >14.25 SI value, **3f** with 32.49 SI value and **3g** with >43.31 SI value. Conversely, the least selective were **4b** with >4.13 SI value, **4e** with 3.84 and **4a** with 2.92 SI value. As observed, most compounds exhibited an SI > 10, clearly indicating their promising potential as antimalarial agents.

These results show the promising potential of the synthesized analogs. All synthesized compounds exhibited antiplasmodial activity and were more active than their respective natural terpene nerol or geraniol.

The (*E*/*Z*) isomerism of the double bond play an important role in the antiplasmodial activity of the monoterpene analogs. The nerol analog (*Z*-isomer, compound **3a**, IC₅₀ = 48.90 ± 7.15 µM) displayed greater antimalarial activity than the geraniol analog (*E*-isomer, compound **4b**, IC₅₀ = 121.10 ± 20.96 µM).

The carbon elongation chain between the nitrogen (N) and oxygen (O) atoms also improve the antiplasmodial activity. The compound **3d** (IC₅₀ = 36.65 ± 5.60 µM) with three carbon atoms spacer displayed greater antiplasmodial activity than the respective analog **3c** (IC₅₀ = 49.71 ± 2.70 µM) with two carbon atoms spacer. For the aminoalcohol **3c** with the neryl side chain, the insertion of alkyl substituent group on the nitrogen (N) atom, only promote sligth variation on the IC_50_ values. These can be observed when the aminoalcohols **3a**, *N*-methyl substituted (IC₅₀ = 48.90 ± 7.15 µM) and **3b**, *N*-ethyl substituted (IC₅₀ = 41.31 ± 3.08 µM) were compared to **3c** (IC₅₀ = 49.71 ± 2.70 µM). However, the insertion of an additional methyl group, as in compound **3g** (IC₅₀ = 23.09 ± 3.74 µM), provides a significant increase in antiplasmodial activity. This enhanced activity may be associated with the permanent positive formal charge formed on the nitrogen atom. Supporting this hypothesis, other compounds with positive formal charge, such as **4c** and **4e** (IC₅₀ = 14.06 ± 1.05 and 8.94 ± 1.31 µM, respectively) also exhibited higher activity.

The replacement of a hydroxyl group (OH) by a bulky group as (S-Me or S-Et) also improves the antiplasmodial activity. These can be observed when the sulfur compounds **3e** (IC₅₀ = 7.49 ± 1.43 µM) and **3f** (IC₅₀ = 3.14 ± 1.18 µM) were compared to **3c** (IC₅₀ = 49.71 ± 2.70 µM). The selectivity index (SI) is also improved when ethyl group in the sulfur atom (S-Et) was changed by methyl group (S-Me).

In general, compounds with a hydroxyl group (terpenyl aminoalcohols) showed low antiplasmodial activity (**3a**-**d**, **4a**-**b**, **6a**- **b**). These results suggest that the electronic and steric effects introduced by the exchange of some groups may contribute to improved molecular interactions with the biological target. These findings highlight the importance of atom selection and substitution patterns in modulating the biological efficacy of the synthesized compounds. **Figure 2** summarizes some key observations regarding the structure-activity relationship (SAR) of the synthesized derivatives.

**Figure 2.**
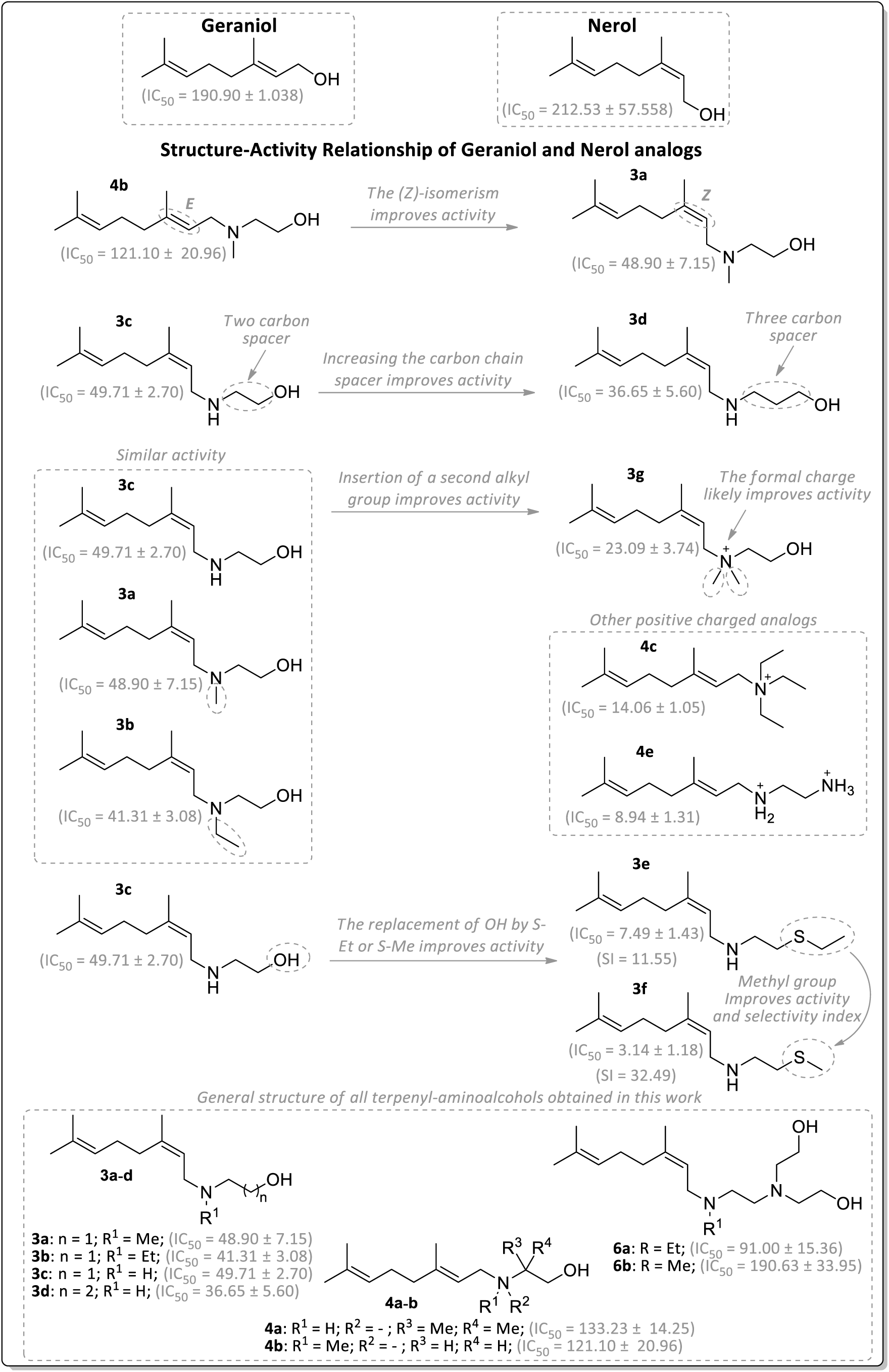
Structure-activity relationship for antiplasmodial activity of geraniol and nerol monoterpene analogs.

### 2.3 ADME-Tox study of acyclic monoterpene analogs

An ADME-Tox study evaluates the behavior and toxicity of compounds in the body, a crucial step in drug development^[18]^. It includes absorption (how the compound is absorbed and its bioavailability), distribution, metabolism (enzymatic transformations that may generate active or toxic metabolites), and excretion (elimination of the compound). Additionally, it analyzes toxicity to identify potential adverse effects, such as genotoxicity or organ toxicity. As previously discussed, one of the challenges in using terpenes as drugs lies in their unfavorable pharmacokinetic characteristics. Therefore, we deemed it essential to conduct an ADME-Tox study to evaluate the pharmacological viability of the new compounds under study, emphasizing predictions of absorption, distribution, and toxicity.

Predictions were performed for TPSA (Topological Polar Surface Area), which measures the polar surface area of a molecule, indicating its ability to form hydrogen bonds. Low TPSA values generally suggest higher cell permeability, while high values indicate lower absorption; WLOGP, a partition coefficient that assesses the lipophilicity of the compound, essential for predicting its solubility in lipids and water, influencing permeability and distribution; The GI Absorption parameter, which indicates the efficiency with which the compound is absorbed in the gastrointestinal tract, categorized as high, moderate, or low; BBB Permeant, a parameter that evaluates whether the compound can cross the blood-brain barrier, a crucial factor for drugs targeting the central nervous system where malaria parasite can be found; the PGP Substrate status, a parameter that checks if the compound is a substrate of P-glycoprotein, an efflux protein that may limit intestinal absorption (Caco-2 permeability) or brain penetration; CL- Plasma (Plasma Clearance) refers to the rate at which the compound is removed from plasma, influencing the duration of its therapeutic action; and the T1/2 (Half-life) time, which represents the time required for the compound’s plasma concentration to be reduced by half, a key parameter for determining dosing frequency. In terms of toxicity, it was decided to evaluate the AMES test, to evaluate the compound’s toxic potential, and Carcinogenicity potential.

The ADME-Tox analysis results indicate that the analyzed molecules have low TPSA values, ranging from 0.00 to 60.69, suggesting good cell permeability and absorption potential for most compounds. WLOGP values for the majority are >1, indicating a favorable range of lipophilicity essential for crossing biological membranes and thus accessing parasites. However, two exceptions, **3g** and **4d**, have negative WLOGP values, which may limit their distribution, as further discussed below. Gastrointestinal absorption also varies, with most compounds exhibiting high absorption, except for **3g** and **4c**, which show low absorption. Regarding blood-brain barrier permeability, most compounds are permeable, with the exception of **3g** and **4c**.

Four compounds, **3g**, **4a**, **4c**, and **4d**, are substrates of P-glycoprotein, which may limit their bioavailability, as this protein is associated with the cellular efflux of drugs. Intestinal permeability values (Caco-2 permeability) range from-4.65 to-5.16, indicating moderate to low permeability. Plasma clearance rates (CL-Plasma) are between 5.87 and 12.63, suggesting that most compounds have moderate elimination rates, while half-life (T1/2) ranges from 0.88 to 1.70, indicating relatively rapid elimination. In mutagenicity tests (AMES), values range from 0.023 to 0.322, with most compounds classified as low mutagenic risk. Regarding carcinogenicity, values range from 0.062 to 0.520, indicating low risk for most compounds, though compounds like **4a** show a higher risk.

Taking in conclusion, most acyclic monoterpenes have predictions to be drug-like compounds. The exceptions are **3g**, **4a**, **4c**, and **4d**, which may present limitations in terms of toxicity and bioavailability. Additionally, the AMDE-Tox study shows that compound **3f** exhibits good permeability (TPSA = 37.33), good solubility levels (WLOGP = 3.63), and high gastrointestinal absorption. It also has the ability to cross the blood-brain barrier and is not a substrate of P-glycoprotein. However, it presents low intestinal permeability (Caco-2 permeability =-4.77). Furthermore, it shows good plasma retention (cl-plasma = 11.46) and a moderate half-life (T1/2 = 1.26). The data also indicate low genotoxicity levels (Ames = 0.374) and low carcinogenic potential (Carcinogenicity = 0.154). For these reasons, compound **3f** appears to be the best antimalarial candidate presented in this study.

### 2.4 Discussion

Several natural terpenes have been shown to inhibit the *in vitro* growth of *Plasmodium falciparum* parasites, highlighting their potential as antimalarial agents. However, a significant limitation of these compounds is their relatively high effective concentrations, often in the millimolar range, which are unlikely to be achievable in clinical settings. Additionally, these terpenes typically suffer from poor bioavailability via preferred administration routes, such as oral administration, further limiting their therapeutic potential^[12–14]^. For example, geraniol is classified as *Generally Recognized As Safe* (GRAS) by the U.S. Food and Drug Administration due to its low toxicity, with a lethal dose 50 (LD_50_) in rats estimated at approximately 3600 mg/kg (oral)^[19]^. In fact, mice treated with 120 mg/kg of geraniol for several weeks showed no toxicity signs but an improvement of anti-oxidative defenses^[20]^. Regarding its bioavailability, studies in rats have demonstrated that geraniol has an absolute oral absorption rate of 92% in the gut and is capable of crossing the blood-brain barrier, reaching the cerebrospinal fluid^[20]^. However, despite its high absorption, geraniol exhibits a short half-life in the bloodstream, calculated to be 12.5 ± 1.5 minutes in rats^[20,21]^. In terms of antimalarial activity, both geraniol and nerol have shown limited efficacy against *Plasmodium* parasites, with IC₅₀ values exceeding 100 µM, indicating weak or negligible antiparasitic effects. This suggests that, while these monoterpenes possess favorable toxicologic properties, their potential as standalone antimalarial agents is limited, and structural modifications may be necessary to enhance their potency and bioavailability.

In this study, we aimed to address these challenges by exploring the antiplasmodial potential of synthetic acyclic monoterpene analogues. These compounds were designed to retain structural similarities to natural terpenes while incorporating modifications to improve their pharmacological properties, including better predicted ADME and toxicity profiles, as well as higher selectivity indices (SI) when compared to HepG2 cells. Discussions on the structure-activity relationship were conducted, emphasizing the effects of various structural modifications introduced in the synthesized analogs. These modifications played a crucial role in modulating the biological activity of the compounds.

The two more active compounds, **3e** (IC₅₀ = 7.49 ± 1.43 µM) and **3f** (IC₅₀ = 3.14 ± 1.18 µM) were also the most lipophilic **3e** (WLOGP = 4.02) and **3f** (WLOGP = 3.63) of all evaluated monoterpene analogs. These results suggest the importance of lipophilicity in the drug design of new lead compounds against *P. falciparum*. Bioinformatic predictions seem to indicate that the new analogues may possess favorable toxicological characteristics, as well as drug-compatible pharmacokinetics and pharmacodynamics.

Our findings demonstrate that Acyclic monoterpene analogues exhibit potent antiplasmodial activity *in vitro*, with significant selectivity against *P. falciparum* parasites in comparison to HepG2 human hepatocellular carcinoma cells. Future studies should aim to elucidate these alternative pathways to provide a comprehensive understanding of the mode of action of these compounds and further optimize their therapeutic potential.

## 3 CONCLUSION

In this work we report the synthesis, antiplasmodial activity and structure activity relationship of fifteen geraniol and nerol monoterpene analogs. These compounds exhibit potent antiplasmodial activity *in vitro* against *P. falciparum* with significant selectivity index. The more potent compound **3f** shows (IC_50_ = 3.14 ± 1.18 µM) and (SI = 32.49), at least 35 times more potent than the respectively monoterpene nerol. The compounds showed improved pharmacological characteristics. Considering these findings, the results presented here pave the way for the development of terpene-like antimalarial drugs with improved efficacy and selectivity.

## 4 EXPERIMENTAL

### 4.1 Biologic assays

#### 4.1.1 Reagents and stock solutions

Albumax I, RPMI-1640 and SYBR Green I nucleic acid gel stain were purchased from Thermo Fisher Scientific (Waltham, Massachusetts, EUA). For *in vitro* use, sterile stock solutions of acyclic monoterpene analogs were prepared at 100 mM in ethanol. All other reagents for biologic assays were purchased from Sigma® (St. Louis, Missouri USA).

#### 4.1.2 *P. falciparum in vitro* culture and synchronization

The *P. falciparum* 3D7 strain parasites were cultured *in vitro* following the Trager and Jensen culture method employing RPMI-1640 medium completed with 0.5% Albumax I into 75 cm^2^ or 25 cm^2^ cell culture flasks at 37 °C^[22,23]^. The culture medium pH was adjusted to 7.4 and was introduced a gas mixture of 5% CO_2_, 5% O_2_ and 90% N_2_ purchased from Air Products Brasil LTDA^®^ (São Paulo, SP, Brazil). Parasites synchronization at ring stages was performed with 5% (w/v) D-sorbitol solution as described previously^[24]^. Parasite development was monitored by Giemsa-stained smears microscopy. PCR for mycoplasma and optic microscopy were used to avoid culture contamination^[25]^.

#### 4.1.3 Monitoring *Plasmodium falciparum* growth

Culture (100 μl) was incubated in a 96-well cell plate in dark at room temperature after adding 100 μl of SYBR green I 2/10,000 (vol/vol) in lysis buffer (20 mM Tris [pH 7.5], 5 mM EDTA, 0.008% saponin [wt/vol], 0.08% Triton X-100 [vol/vol])^[26]^. Fluorescence was measured using a POLARstar Omega fluorometer (BMG Labtech, Ortenberg, Germany) with the excitation and emission bands centered at 485 and 530 nm, respectively. The fluorescence values of uninfected erythrocytes were subtracted from the values obtained for infected cells. Results were analyzed by GraphPad Prism® software to determine percentage of viability respect controls. The dose-response curve was estimated together with the concentration of drug required to cause a 50% reduction in parasite proliferation (IC_50_ value). Results were analyzed by GraphPad Prism software by adjusting data to a dose–response curve to determine the IC_50_ value.

#### 4.1.4 HepG2 cell culture

The HepG2 cells were grown routinely in 75 or 25 cm^2^ flasks in RPMI medium supplemented with 10% FBS and 10 mg/L gentamicin sulfate. The cultures were maintained in a humidified incubator with 5% CO_2_ at 37 °C. Cells were manipulated following the passage and trypsinization procedures described elsewhere^[27,28]^. PCR for mycoplasma and optic microscopy were used to avoid culture contamination^[29]^.

#### 4.1.5 Cell viability assays

For cell-viability experiments, confluent cultures were washed in Phosphate Buffered Saline (PBS), trypsinized, centrifuged at 300 g, and suspended in culture media. The cells were cultured in 96-well plates at a density of 10.000 cells/well. The next day, the cells were subjected to different concentrations of drugs. Ethanol controls were performed to ensure no effects related to solvents, and its concentration was always ≥1%. After 48 hours, it was added to cells 10 µL PBS containing 10 mg/L MTT and incubated at 37°C. After 4 hours, 100 µL isopropanol 0.04 M HCl was added to each well. The next day, the absorbance at 595 nm wavelength corrected to 690 nm was monitored in a POLARstar Omega fluorometer® (BMG Labtech®, Ortenberg, Germany), and the results were analyzed by GraphPad Prism® software to determine percentage of viability respect controls. In some cases, the dose-response curve was estimated together with the concentration of drug required to cause a 50% reduction in cell viability (CC_50_ value). Results were analyzed by GraphPad Prism software by adjusting data to a dose–response curve to determine the CC_50_ value.

#### 4.1.6 Selectivity Index

The selectivity index (SI) is a ratio that measures the therapeutic window between antiplasmodial activity and cytotoxicity. SI was calculated by dividing the IC_50_ value in HepG2 cells by the IC_50_ value in *Plasmodium* parasites. An SI value >10 was considered promising^[30]^.

#### 4.1.7 Bioinformatics

Computational modeling to estimate the bioavailability, aqueous solubility, blood brain barrier potential, human intestinal absorption, mutagenicity, and toxicity for the compounds was performed using the SwissAdme and ADMETlab 3.0 plataforms^[18]^ (Last accessed in January 2025).

### 4.2 Chemistry

#### 4.2.1 Chemicals and general methods

All reagents and solvents were reagent grade and were used without prior purification. All reactions were monitored by thin-layer chromatography (TLC, Sigma-Aldrich^®^ 60). The terpenyl bromides (**3** and **4**) and the mesylated intermediates (**5a**-**b**) were purified only by liquid-liquid extraction, and the compounds (**3a**-**h**, **4a**-**e** and **6a**-**b**) were purified by liquid-liquid extraction followed by flash chromatography on silica gel Sigma-Aldrich^®^ 60 (230-400 mesh) using CH_2_Cl_2_:CH_3_OH or CH_2_Cl_2_:CH_3_OH:NH_4_OH as eluent. The IR spectra were acquired on a PerkinElmer Spectrum 100 FTIR spectrophotometer with an Attenuated Total Reflectance (ATR) attachment, and only significant bands were recorded. ^1^H NMR (300 MHz) and ^13^C NMR (75 MHz) spectra of the final compounds (**3a**-**h**, **4a**-**e** and **6a**-**b**) were recorded on solutions in CDCl_3_ on a Bruker Avance ACX300 (300MHz). The chemical shifts (δ) were reported in parts per million (ppm) with reference to CDCl_3_ (^13^C, 77.00) or residual CHCl_3_ (^1^H, 7.26). The NMR coupling constants (*J*) were recorded in Hertz. Splitting pattern abbreviations are as follows: s = singlet, d = doublet, t = triplet, td = distorted triplet, and m = multiplet. ESI-HRMS mass spectra were carried out on a Bruker MicroTOF spectrometer. The melting points were determined on an Allerbest apparatus.

#### 4.2.2 General procedure for synthesis of neryl and geranyl bromides 3 and 4

Nerol (**1**) and geraniol (**2**) ((*Z*)-3,7-dimethylocta-2,6-dien-1-ol and (*E*)-3,7-dimethylocta-2,6-dien-1-ol) were used as starting material to prepare the intermediate bromides. The terpenes (10 mmol) were solubilized in THF (10 mL), and a solution of PBr _3_ (3.3 mmol) in THF (5 mL) was dropwise added. The reaction mixture was stirred for 40 minutes at-18 °C for geraniol and 90 minutes at-35 °C for nerol. After, the solution was concentrated in vacuo, and the residual oil was solubilized in diethyl ether/hexane (20 mL; 1:1 v/v) and washed sequentially with NaHCO_3_ (2 x 10 mL; 5% m/v) and distilled water (2 x 10 mL). The organic phase was dried over anhydrous Na_2_SO_4_, and the solvent was removed under reduced pressure to furnish the terpenyl bromides **3** and **4**.

#### 4.2.3 General procedure for synthesis of the monoterpene derivatives 3a-h and 4a-e

The terpenyl bromides (0,5 mmol) were solubilized in dry dichloromethane (5 mL) and dropwise added to the respective excess of different amines (solubilized in 8 mL of dichlorometane) at room temperature: 2-(Methylamino)ethanol (5 mmol for **3a** and **4b**), 2-(Ethylamino)ethanol (5 mmol for **3b**), Ethanolamine (10 mmol for **3c**), Propanolamine (10 mmol for **3d**), 2- (Ethylthio)ethylamine (10mmol for **3e**), 2-(Methylthio)ethylamine (10mmol for **3f**), 2-(Dimethylamino)ethanol (1,1 mmol for **3g**), 1,4-diaminobutane (10mmol for **3h**), 2-Amino-2-methyl-1-propanol (10mmol for **4a**), Triethylamine (1,1 mmol for **4c**), and Triethanolamine (1,1 mmol for **4d**). The reactions were maintained under these conditions for 24 hours. After, the solution was washed sequentially with distilled water (2 x 10 mL) to remove excess amine. The organic phase was dried over anhydrous Na_2_SO_4_ and the solvent was removed under reduced pressure. The residual oil was purified by *flash* chromatography on silica gel using CH_2_Cl_2_:CH_3_OH:NH_4_OH as eluent to provide monoterpene analogs (**3a**-**h** and **4a**-**e**). Compound **4e** was transformed in your respective hydrochloride salt according to the methodology previously described by our research group^[15]^.

*(Z)*-2-((3,7-dimethylocta-2,6-dien-1-yl)(methyl)amino)ethanol (**3a**): A yellow oil (31%). FTIR (ATR, cm^-1^): 3352 (υ, O-H); 1668 (υ, C=C); 1135 (υ, C-O). ^1^H NMR (300 MHz, CDCl_3_): δ 5.26 (t, *J* = 6.8 Hz, 1H), 5.07 (s, 1H), 3.67 (t, *J* = 6 Hz, 2H), 3.16 (d, *J* = 6.8 Hz, 2H), 2.64 (t, *J* = 6 Hz, 2H), 2.34 (s, 3H), 2.06 (m, 4H), 1.75 (s, 3H), 1.67 (s, 3H), 1.59 (s, 3H). ^13^C NMR (75 MHz, CDCl_3_): δ 140.80 (C), 132.1 (C), 123.7 (CH), 120.0 (CH), 58.1 (CH_2_), 58.0 (CH_2_), 54.8 (CH_2_), 41.3 (CH_3_), 32.2 (CH_2_), 26.4 (CH_2_), 25.7 (CH_3_), 23.6 (CH_3_), 17.7 (CH_3_). ESI-HRMS [M+H]^+^ *m/z* calculated for C_13_H_26_NO = 212.2009, found = 212.2001.

*(Z)*-2-((3,7-dimethylocta-2,6-dien-1-yl)(ethyl)amino)ethanol (**3b**): A yellow oil (47%). FTIR (ATR, cm^-^¹): 3391 (υ, O-H); 1667 (υ, C=C); 1071 (υ, C-N); 1043 (υ, C-O). ^1^H NMR (300 MHz, CDCl_3_): δ 5.25 (t, *J* = 6.9 Hz, 1H), 5.07 (m, 1H), 3.63 (t, *J* = 6.0 Hz, 2H), 3.24 (d, 6.9 Hz, 2H), 2.71 – 2.64 (m, 4H), 2.06 (m, 4H), 1.74 (s, 3H), 1.67 (s, 3H), 1.60 (s, 3H), 1.11 (t, *J* = 7.2 Hz, 3H). ^13^C NMR (75 MHz, CDCl_3_): δ 140.6 (C), 132.1 (C), 123.7 (CH), 119.9 (CH), 57.9 (CH_2_), 54.6 (CH_2_), 50.4 (CH_2_), 47.5 (CH_2_), 32.2 (CH_2_), 26.4 (CH_2_), 25.7 (CH_3_), 23.6 (CH_3_), 17.7 (CH_3_), 11.4 (CH_3_). ESI-HRMS [M+H]^+^ *m/z* calculated for C_14_H_28_NO = 226.2166, found = 226.2166.

*(Z)*-2-((3,7-dimethylocta-2,6-dien-1-yl)amino)ethanol (**3c**): A yellow oil (26%). FTIR (ATR, cm^-^¹): 3304 (υ, O-H + N-H); 1668 (υ, C=C); 1070 (υ, C-N); 1050 (υ, C-O). ^1^H NMR (300 MHz, CDCl_3_): δ 5.28 (t, *J* = 6.8 Hz, 1H), 5.07 (s, 1H), 3.70 (t, *J* = 6.0 Hz, 2H), 3.31 (d, *J* = 6.8 Hz, 2H), 3.17 (s, 1H, O-H), 2.81 (t, *J* = 6.0 Hz, 2H), 2.06 (m, 4H), 1.73 (s, 3H), 1.67 (s, 3H), 1.59 (s, 3H).

^13^C NMR (75 MHz, CDCl_3_): δ 140.1 (C), 132.2 (C), 123.7 (CH), 121.2 (CH), 60.1 (CH_2_), 50.1 (CH_2_), 46.0 (CH_2_), 32.1 (CH_2_), 26.5 (CH_2_), 25.7 (CH_3_), 23.4 (CH_3_), 17.7 (CH_3_). ESI-HRMS [M+H]^+^ *m/z* calculated for C_12_H_24_NO = 198.1853, found = 198.1857.

(*Z*)-3-((3,7-dimethylocta-2,6-dien-1-yl)amino)propan-1-ol (**3d**): A yellow oil (20%). FTIR (ATR, cm^-^¹): 3291 (υ, O-H + N-H); 1667 (υ, C=C); 1105 (υ, C-N); 1066 (υ, C-O). ^1^H NMR (300 MHz, CDCl_3_): δ 5.25 (t, *J* = 6.9 Hz, 1H), 5.07 (s, 1H), 3.80 (t, *J* = 5.3 Hz, 2H), 3.27 (d, *J* = 6.9 Hz, 2H), 3.08 (s, 1H, O-H) 2.89 (t, *J* = 5.7 Hz, 2H), 2.06 (m, 4H), 1.76-1.72 (m, 5H), 1.67 (s, 3H), 1.59 (s, 3H). ^13^C NMR (75 MHz, CDCl_3_): δ 139.9 (C), 132.1 (C), 123.7 (CH), 121.4 (CH), 63.5 (CH_2_), 48.5 (CH_2_), 46.4 (CH_2_), 32.1 (CH_2_), 30.2 (CH_2_), 26.5 (CH_2_), 25.7 (CH_3_), 23.4 (CH_3_), 17.7 (CH_3_). ESI-HRMS [M+H]^+^ *m/z* calculated for C_13_H_26_NO = 212.2009 found = 212.2007.

*(Z)*-*N*-(2-(ethylthio)ethyl)-3,7-dimethylocta-2,6-dien-1-amine (**3e**): A yellow oil (36%). FTIR (ATR, cm^-^¹): 3302 (υ, N-H); 1668 (υ, C=C); 1109 (υ, C-N). ^1^H NMR (300 MHz, CDCl_3_): δ 5.32 (t, *J* = 6.5 Hz, 1H), 5.07 (s, 1H), 3.38 (d, *J* = 6.5 Hz, 2H), 2.92 – 2.77 (m, 4H), 2.55 (q, *J* = 7.4 Hz, 2H), 2.06 (m, 4H), 1.74 (s, 3H), 1.67 (s, 3H), 1.59 (s, 3H), 1.25 (t, *J* = 7.4 Hz, 3H). ^13^C NMR (75 MHz, CDCl_3_) δ 141.1 (C), 132.2 (C), 123.6 (CH), 120.0 (CH), 47.0 (CH_2_), 45.7 (CH_2_), 32.2 (CH_2_), 30.0 (CH_2_), 26.5 (CH_2_), 25.9 (CH_2_), 25.7 (CH_3_), 23.5 (CH_3_), 17.7 (CH_3_), 14.8 (CH_3_). ESI-HRMS [M+H]^+^ *m/z* calculated for C_14_H_28_NS = 242.1937 found = 242.1931.

*(Z)*-3,7-dimethyl-*N*-(2-(methylthio)ethyl)octa-2,6-dien-1-amine (**3f**): A yellow oil (37%). FTIR (ATR, cm^-^¹): 3301 (υ, N-H); 1668 (υ, C=C); 1108 (υ, C-N). ^1^H NMR (300 MHz, CDCl_3_): δ 5.34 (t, *J* = 6.8 Hz, 1H), 5.07 (s, 1H), 3.42 (d, *J* = 6.8 Hz, 2H), 2.93 (t, *J* = 6.8 Hz, 2H), 2.79 (t, *J* = 6.8 Hz, 2H), 2.11 (s, 3H), 2.07 (m, 4H), 1.75 (s, 3H), 1.68 (s, 3H), 1.59 (s, 3H). ^13^C NMR (75 MHz, CDCl_3_): δ 141.8 (C), 132.3 (C), 123.5 (CH), 119.2 (CH), 46.2 (CH_2_), 45.6 (CH_2_), 32.2 (CH_2_), 32.1 (CH_2_), 26.4 (CH_2_), 25.7 (CH_3_), 23.4 (CH_3_), 17.7 (CH_3_), 15.3 (CH_3_). ESI-HRMS [M+H]^+^ *m/z* calculated for C_13_H_26_NS = 228.1781 found = 228.1774.

(*Z*)-*N*-(2-hydroxyethyl)-*N,N*,3,7-tetramethylocta-2,6-dien-1-aminium bromide (**3g**): A pale-yellow solid (65%). mp (°C): 155-160. FTIR (ATR, cm^-^¹): 3361 (υ, N-H); 1660 (υ, C=C); 1085 (υ, C-O), 1036 (υ, C-N). ^1^H NMR (300 MHz, CDCl_3_) δ 5.39 (t, *J* = 7.9 Hz, 1H), 5.05 (t, *J* = 6.9 Hz, 1H), 4.11-4.08 (m, 4H), 3.68 (td, 2H), 3.26 (s, 6H), 2.19 – 2.08 (m, 4H), 1.86 (s, 3H), 1.65 (s, 3H), 1.57 (s, 3H). ^13^C NMR (75 MHz, CDCl_3_): δ 151.5 (C), 132.9 (C), 122.8 (CH), 111.3 (CH), 65.2 (CH_2_), 63.2 (CH_2_), 56.0 (CH_2_), 50.9 (CH_3_ x2), 32.4, 26.1 (CH_2_), 25.7 (CH_3_), 24.0 (CH_3_), 17.8 (CH_3_). ESI-HRMS [M]^+^ *m/z* calculated for C_14_H_28_NO = 226.2166 found = 226.2169.

*(Z)-N^1^*-(3,7-dimethylocta-2,6-dien-1-yl)butane-1,4-diamine (**3h**): A yellow oil (37%). FTIR (ATR, cm^-^¹): 3352 (υ_as_, N-H); 3285 (υ_s_, N-H); 1668 (υ, C=C); 1587 (d, NH_2_) 1108 (υ, C-N). ^1^H NMR (300 MHz, CDCl_3_): δ 5.26 (t, *J* = 6.7 Hz, 1H), 5.07 (s, 1H), 5.71 (s, 1H, N-H), 3.25 (d, *J* = 6.8 Hz, 2H), 2.75 (t, *J* = 6 Hz, 2H), 2.64 (d, *J* = 6.3 Hz, 2H), 2.04 (m, 4H), 1.71 (s, 3H), 1.66 (s, 3H), 1.57 (s, 7H). ^13^C NMR (75 MHz, CDCl_3_): δ 139.2 (C), 132.0 (C), 123.8 (CH), 122.0 (CH), 48.6 (CH_2_), 46.5 (CH_2_), 41.4 (CH_2_), 32.1 (CH_2_), 30.3 (CH_2_), 26.9 (CH_2_), 26.5 (CH_2_), 25.8 (CH_3_), 23.4 (CH_3_), 17.7 (CH_3_). ESI-HRMS [M+H]^+^ *m/z* calculated for C_14_H_29_N_2_ = 225.2326 found = 225.2334.

*(E*)-2-((3,7-dimethylocta-2,6-dien-1-yl)amino)-2-methylpropan-1-ol (**4a**): A yellow oil (34%). FTIR (ATR, cm^-^¹): 3407 (υ, O-H + N-H); 1667 (υ, C=C); 1164 (υ, C-N); 1052 (υ, C-O). ^1^H NMR (300 MHz, CDCl_3_): δ 5.22 (td, 1H), 5.10 (td, 1H), 3.30 (s, 2H), 3.12 (d, *J* = 6.8 Hz, 2H), 2.17 – 1.97 (m, 4H), 1.67 (s, 3H), 1.63 (s, 3H), 1.59 (s, 3H), 1.08 (s, 6H). ^13^C NMR (75 MHz, CDCl_3_): δ 137.8 (C), 131.6 (C), 124.1 (CH), 122.8 (CH), 68.4 (CH_2_), 53.7 (C), 39.6 (CH_2_), 39.5 (CH_2_), 26.5 (CH_2_), 25.7 (CH_3_), 24.0 (2xCH_3_), 17.7 (CH_3_), 16.2 (CH_3_), 17.7 (CH_3_).

*(E*)-((3,7-dimethylocta-2,6-dien-1-yl)(methyl)amino)ethanol (**4b**): A yellow oil (71%). FTIR (ATR, cm^-^¹): 3355 (υ, O-H); 1667 (υ, C=C); 1073 (υ, C-N); 1030 (υ, C-O). ^1^H NMR (300 MHz, CDCl_3_): δ 5.29 (td, 1H), 5.06 (td, 1H), 3.74 (t, *J* = 5.3 Hz, 2H), 3.28 (d, *J* = 7.1 Hz, 2H), 2.89 (s, 1H), 2.74 (t, *J* = 5.3 Hz, 2H), 2.43 (s, 3H), 2.18 – 2.05 (m, 4H), 1.67 (s, 6H), 1.60 (s, 3H). ^13^C NMR (75 MHz, CDCl_3_): δ 142.4 (C), 131.9 (C), 123.7 (CH), 117.7 (CH), 58.1 (CH_2_), 57.7 (CH_2_), 54.8 (CH_2_), 41.1 (CH_3_), 39.8 (CH_2_), 26.3 (CH_2_) 25.7 (CH_3_), 17.7 (CH_3_), 16.5 (CH_3_).

*(E)-N,N,N-*triethyl-3,7-dimethylocta-2,6-dien-1-aminium bromide (**4c**): A brown light solid (54%). mp (°C): 86-88. FTIR (ATR, cm^-^¹): 1663 (υ, C=C); 1011 (υ, C-N). ^1^H NMR (300 MHz, CDCl_3_): δ 5.18 (t, *J* = 7.8 Hz, 1H), 4.99 (s, 1H), 4.04 (d, *J* = 7.8 Hz, 2H), 3.43 (q, *J* = 7.3 Hz, 6H), 2.14-2.13 (m, 4H), 1.82 (s, 3H), 1.66 (s, 3H), 1.59 (s, 3H), 1.40 (t, *J* = 7.3 Hz, 9H). ^13^C NMR (75 MHz, CDCl_3_): δ 151.0 (C), 132.6 (C), 123.1 (CH), 109.9 (CH), 56.1 (CH_2_), 53.0 (2xCH_2_), 40.1 (CH_2_), 26.0 (CH_3_), 25.8 (CH_3_), 17.8 (CH_3_), 17.5 (CH_3_), 8.3 (CH_3_).

*(E)-N,N,N-*tris(2-hydroxyethyl)-3,7-dimethylocta-2,6-dien-1-aminium bromide (**4d**): A white solid (85%). mp (°C): 95-97. FTIR (ATR, cm^-^¹): 3393, 3334 and 3204 (υ, O-H); 1657 (υ, C=C); 1100 (υ, C-N), 1045 (υ, C-O). ^1^H NMR (300 MHz, CDCl_3_): δ 5.38 (td, 1H), 5.02 (t, *J* = 7.0 Hz, 1H), 4.77 (s, 3H), 4.24 (d, *J* = 7.4 Hz, 2H), 3.70 (sl, 6H), 2.14 (sl, 4H), 1.79 (s, 3H), 1.67 (s, 3H), 1.57 (s, 3H). ^13^C NMR (75 MHz, CDCl_3_): δ 150.7 (C), 132.4 (C), 123.3 (CH), 110.9 (CH), 61.4 (3xCH_2_), 59.7 (CH_2_), 55.8 (3xCH_2_), 40.2 (CH_2_), 26.2 (CH_2_), 17.9 (CH_3_), 17.5 (CH_3_), 8.3 (3xCH_3_). ESI-HRMS [M+H]^+^ *m/z* calculated for C_16_H_32_NO_3_ = 286.2377 found = 286.2373.

#### 4.2.4 General procedure for synthesis of mesylated intermediates 5a-b

In a 25 mL flask, 1 mmol of the derivatives **3a** and **3b** was dissolved in 1 mL of dichloromethane (DCM), previously dried over molecular sieves. To this solution, a stoichiometric amount of 1.5 mmol of methanesulfonyl chloride (MsCl) was added per mmol of products **3a** and **3b**. The reaction mixture was then cooled in an ice bath (0°C), and a solution of 2 mmol of triethylamine in 1 mL of DCM was added dropwise over 30 minutes with constant stirring. The temperature was maintained only during the triethylamine addition, after which the system was allowed to gradually warm to room temperature. The reaction mixture was then left under these conditions for 12 hours. The intermediate compounds **5a** and **5b** were used immediately without purification. The formation of mesylated products was confirmed by thin-layer chromatography (TLC) and infrared (IR) spectroscopy.

#### 4.2.5 General procedure for synthesis of the nerol derivatives 6a-b

In a 25 mL flask, 1 mmol of the derivatives **5a** and **5b** were dissolved in 1 mL of dimethylformamide (DMF). To this solution, a stoichiometric amount of 1.5 mmol of diethanolamine (solubilized in 2mL of DMF) was added. The reaction mixture kept stirring at room temperature for 18 hours. After, the solution was washed sequentially with distilled water (2 x 10 mL) to remove excess amine. The organic phase was dried over anhydrous Na_2_SO_4_ and the solvent was removed under reduced pressure. The residual oil was purified by *flash* chromatography on silica gel using CH_2_Cl_2_:CH_3_OH:NH_4_OH as eluent to provide neryl aminoalcohols **6a** and **6b**.

*(Z)*-2,2’-((2-((3,7-dimethylocta-2,6-dien-1-yl)(ethyl)amino)ethyl)azanediyl)diethanol (**6a**): A yellow oil (29%). FTIR (ATR, cm^-1^): 3362 (υ, O-H); 1667 (υ, C=C); 1068 (υ, C-N); 1139 (υ, C-O). ^1^H NMR (300 MHz, CDCl_3_): δ 5.26 (t, *J* = 6.7 Hz, 1H), 5.07 (s, 1H), 3.57 (t, *J* = 6.0 Hz, 2H), 3.16 (d, *J* = 6.7 Hz, 2H), 2.69 – 2.52 (m, 10H), 2.04 (m, 4H), 1.74 (s, 3H), 1.66 (s, 3H), 1.59 (s, 3H), 1.08 (t, *J* = 7.1 Hz, 3H). ^13^C NMR (75 MHz, CDCl_3_) δ 140.2 (C), 132.0 (C), 123.8 (CH), 119.8 (CH), 59.9 (CH_2_), 57.6 (CH_2_), 52.4 (CH_2_), 51.7 (CH_2_), 50.0 (CH_2_), 47.1 (CH_2_), 32.2 (CH_2_), 26.5 (CH_2_), 25.8 (CH_3_), 23.6 (CH_3_), 17.7 (CH_3_), 10.7 (CH_3_). ESI-HRMS [M+H]^+^ *m/z* calculated for C_18_H_37_N_2_O_2_ = 313.2850, found = 313.2853.

*(Z)-*2,2’-((2-((3,7-dimethylocta-2,6-dien-1-yl)(methyl)amino)ethyl)azanediyl)diethanol (**6b**): A yellow oil (45%). FTIR (ATR, cm^-1^): 3351 (υ, O-H); 1667 (υ, C=C); 1135 (υ, C-O). ^1^H NMR (300 MHz, CDCl_3_): δ 5.27 (t, *J* = 7.0 Hz, 1H), 5.06 (s, 1H), 3.59 (t, *J* = 6.0 Hz, 4H), 3.22 (d, *J* = 7.0 Hz, 2H), 2.69 – 2.59 (m, 8H), 2.35 (s, 3H), 2.06 (m, 4H), 1.76 (s, 3H), 1.66 (s, 3H), 1.59 (s, 3H).

^13^C NMR (75 MHz, CDCl_3_) δ 142.0 (C), 132.2 (C), 123.6 (CH), 118.5 (CH), 60.0 (CH_2_ x2), 58.1 (CH_2_ x2), 55.1 (CH_2_), 54.4 (CH_2_), 51.4 (CH_2_), 40.9 (CH_3_), 32.1 (CH_2_), 26.4 (CH_2_), 25.7 (CH_3_), 23.6 (CH_3_), 17.7 (CH_3_). ESI-HRMS [M+H]^+^ *m/z* calculated for C_17_H_35_N_2_O_2_ = 299.2694, found = 299.2695.

## AUTHOR CONTRIBUTIONS

MMF, MC, GOC, MAP, MFSS, GSS, and IBV contributed to conceptualization, formal analysis, investigation, methodology and writing. IASS, CAAS, LVNL, AVF, JVS and MVNS contributed to the synthesis and characterization of described compounds. IBV, MFS and AMK also contributed in project administration, funding acquisition, supervision and writing – review & editing.

## FUNDING

MFSS is fellow of the *Coordenação de Aperfeiçoamento de Pessoal de Nível Superior* (CAPES). IBV and MMF are fellows from the *Fundação de Amparo à Pesquisa do Estado de São Paulo* (FAPESP); MMF FAPESP process number: 2023/12343-1; IBV FAPESP process number: 2019/13419-6). LVNL is fellow of *Conselho Nacional de Desenvolvimento Científico e Tecnológico* (CNPq). GSS and CAAS are fellows from the *Fundação de Amparo à Pesquisa do Estado de Minas Gerais* (FAPEMIG). This work was also supported funding from CAPES, FAPESP (2024/09997-2) (AMK), FAPEMIG (APQ-01455-22) (MFS) and *Conselho Nacional de Desenvolvimento Científico e Tecnológico* (CNPq). The authors acknowledge the LRMN-UNIFAL (Federal University of Alfenas) for nuclear magnetic resonance analysis and the RELAM-UNIFEI (APQ-002290-23)

## ACKNOWLEDGEMENTS

We thank the Blood Center of Sírio Libanês Hospital (São Paulo, Brazil), for the gift of erythrocytes.

## CONFLICT OF INTEREST

The authors declare no conflicts of interest.

## Entry for the Table of Contents

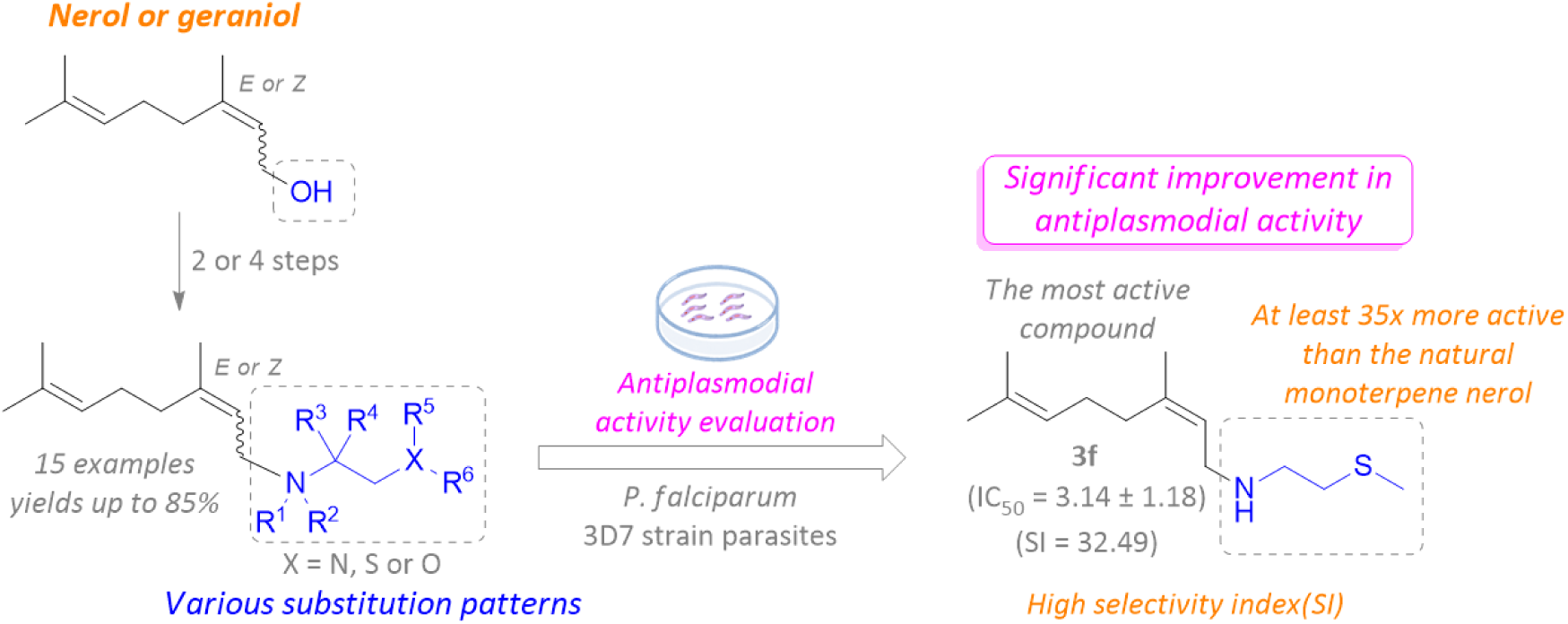

In this work, we synthesized and evaluated the antiplasmodial activity of fifteen analogues of the natural monoterpenes geraniol or nerol. The evaluated compounds exhibited strong inhibitory activity, high selectivity index (SI), and promising pharmacological properties based on bioinformatic analyses. Compound **3f** emerged as the most potent, being at least 35 times more active than nerol, with remarkable pharmacological predictions. The findings presented here highlight the potential of these compounds as starting points for new antimalarial drugs.

## Supplementary material

Compound 3a

**Figure 1.**
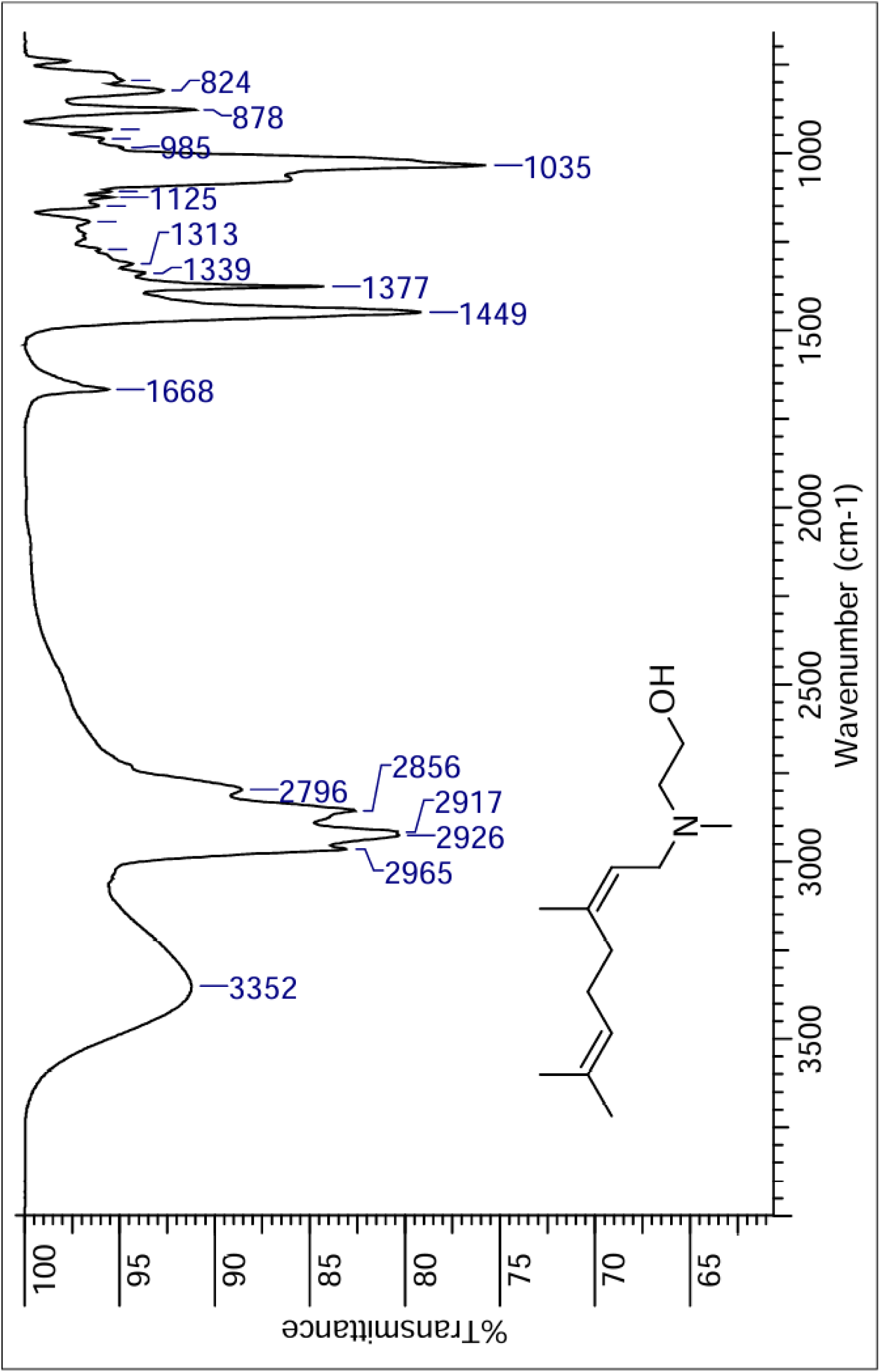
IR spectra of compound **3a**.

**Figure 2.**
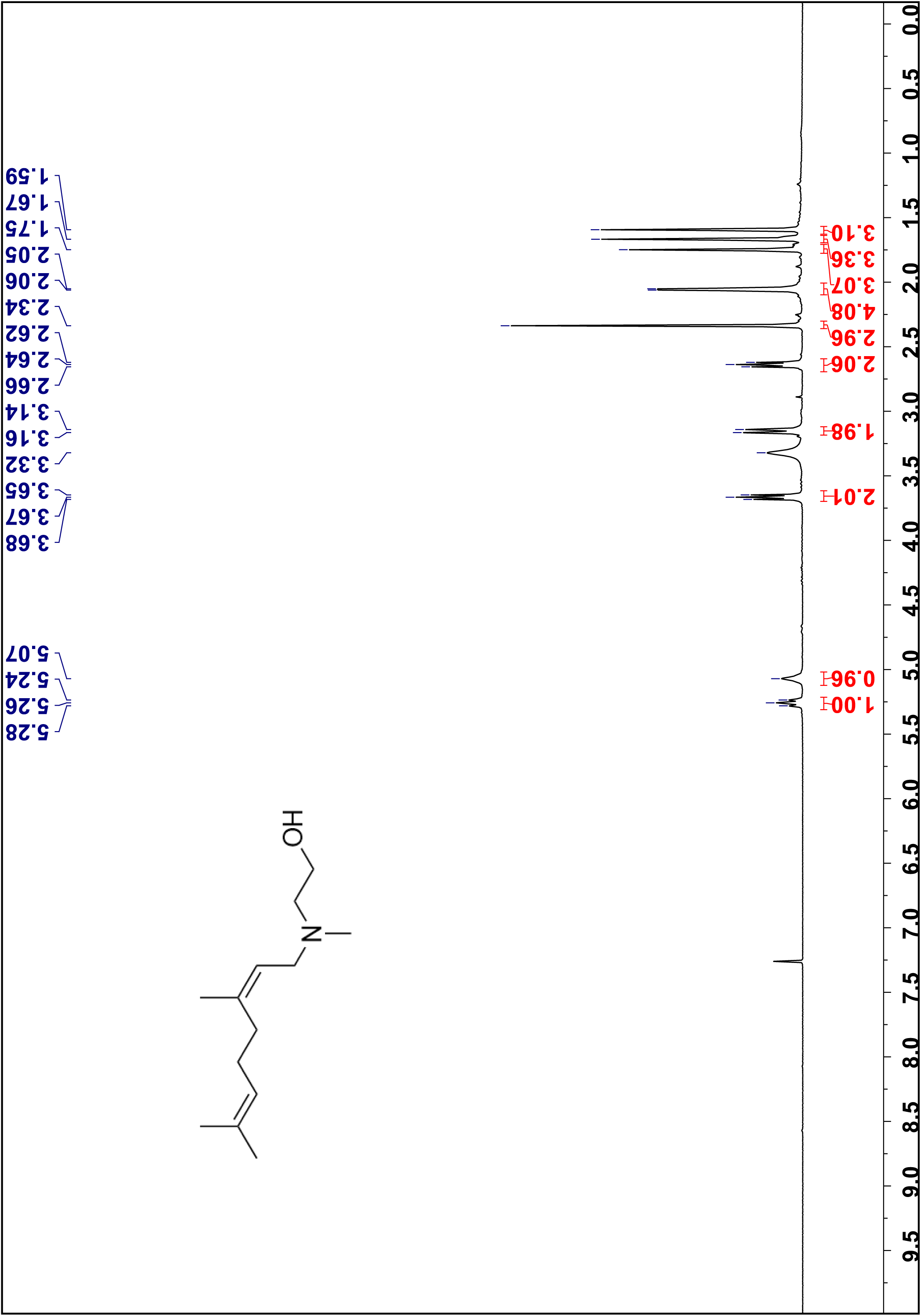
¹H NMR spectra of compound **3a** in CDCl_3_, 300 MHz.

**Figure 3.**
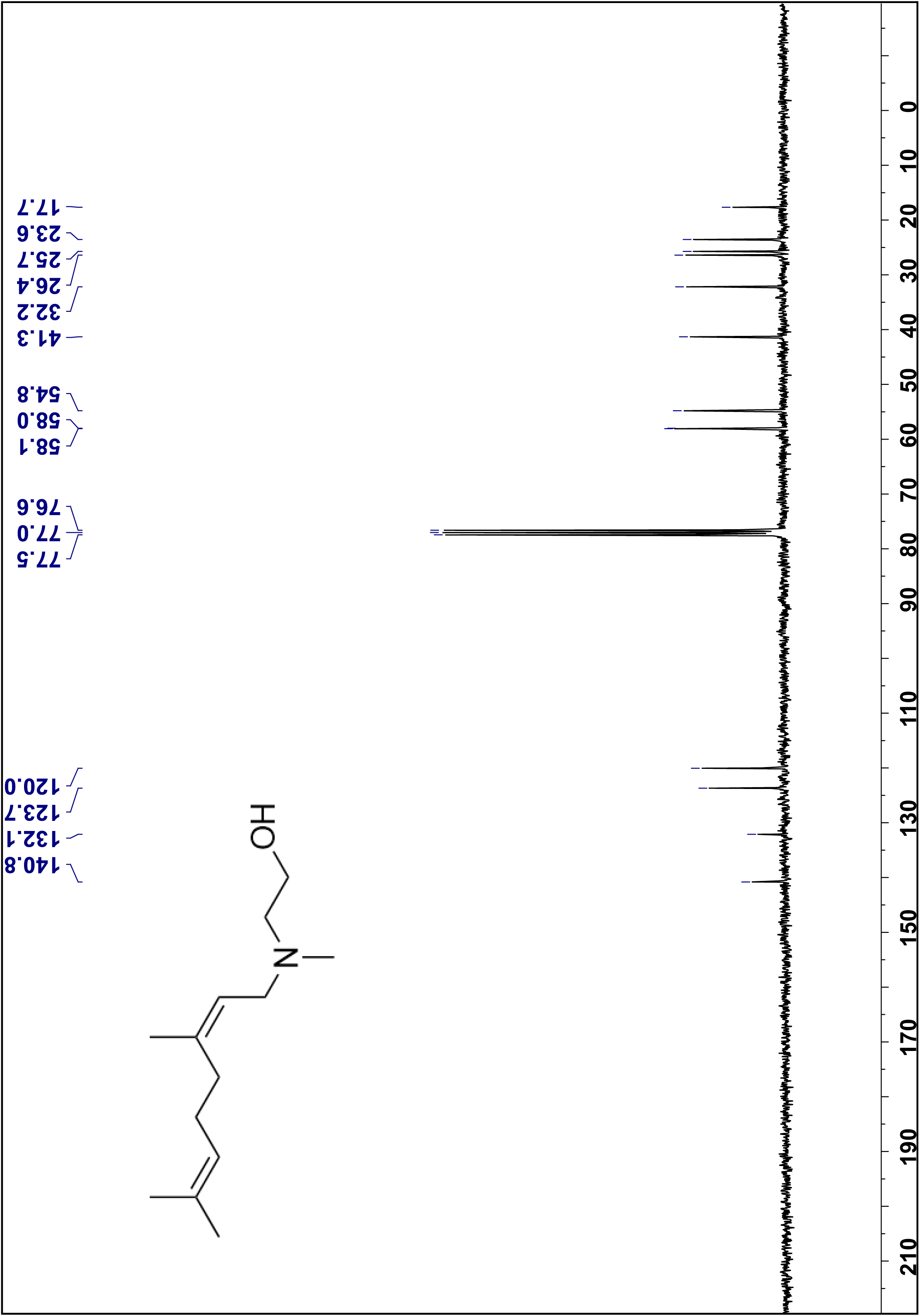
¹³C NMR spectra of compound **3a** in CDCl_3_, 75 MHz.

**Figure 4.**
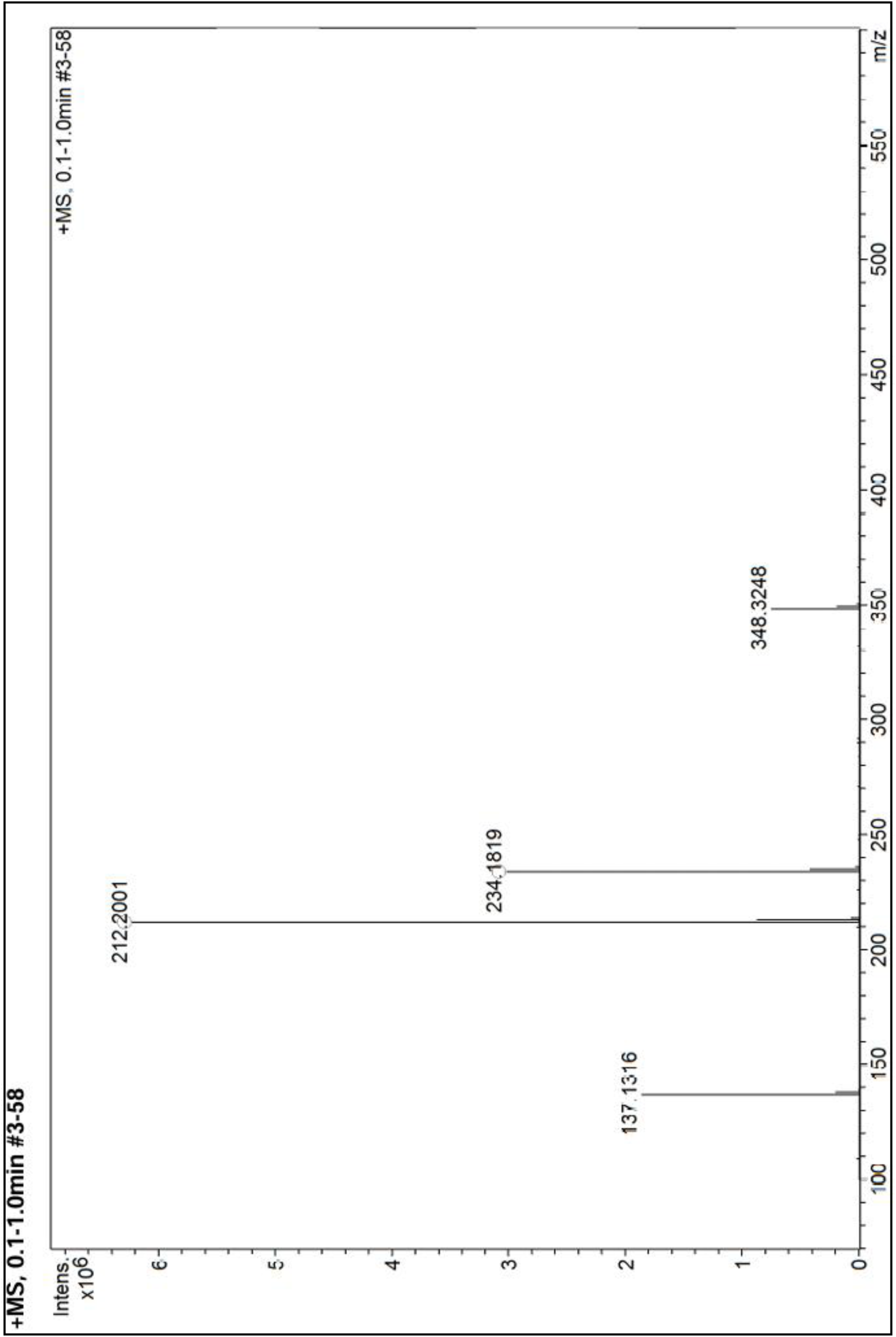
High Resolution Mass spectra of compound **4a**.

Compound 3b

**Figure 5.**
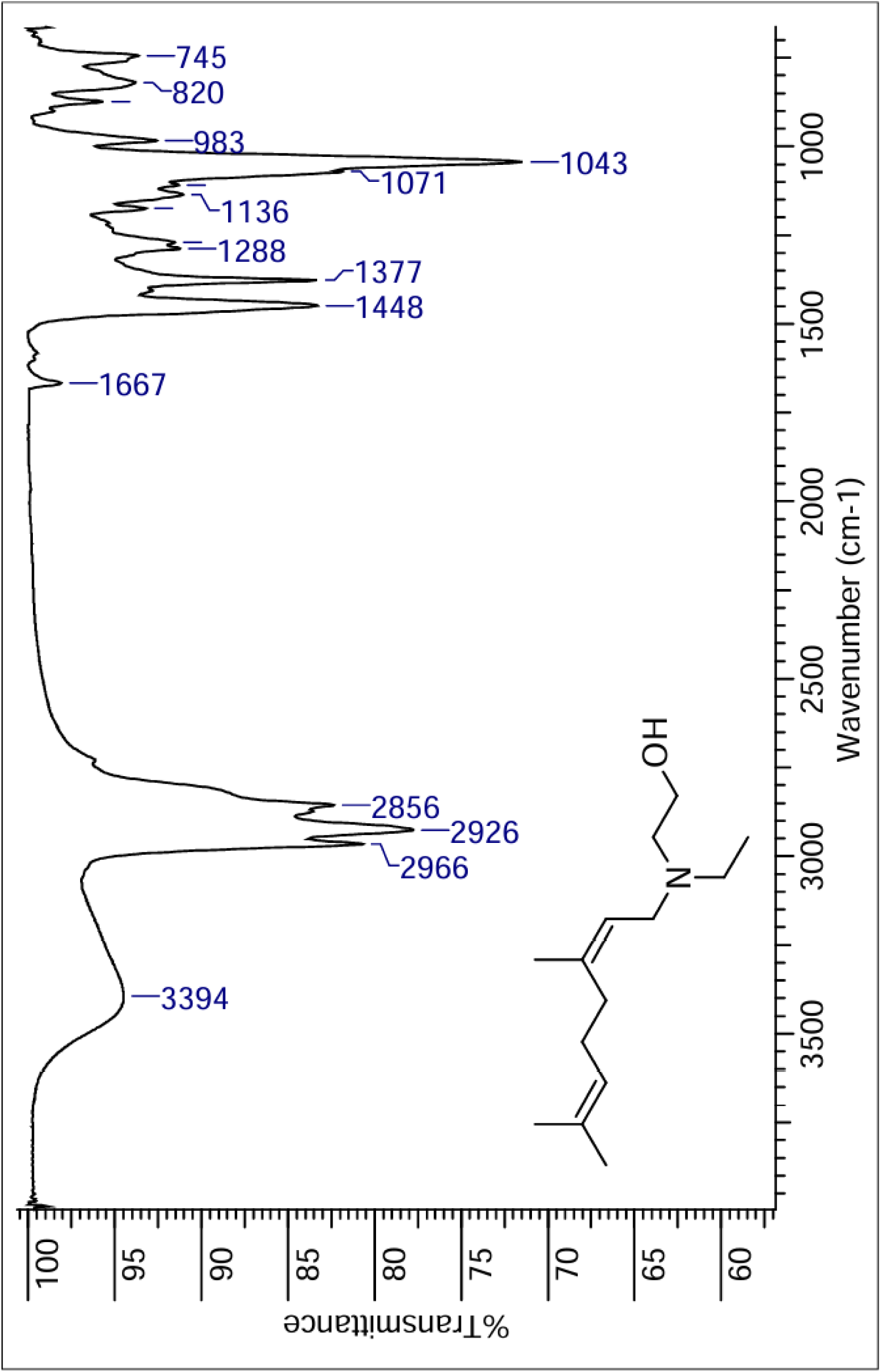
IR spectra of compound **3b**.

**Figure 6.**
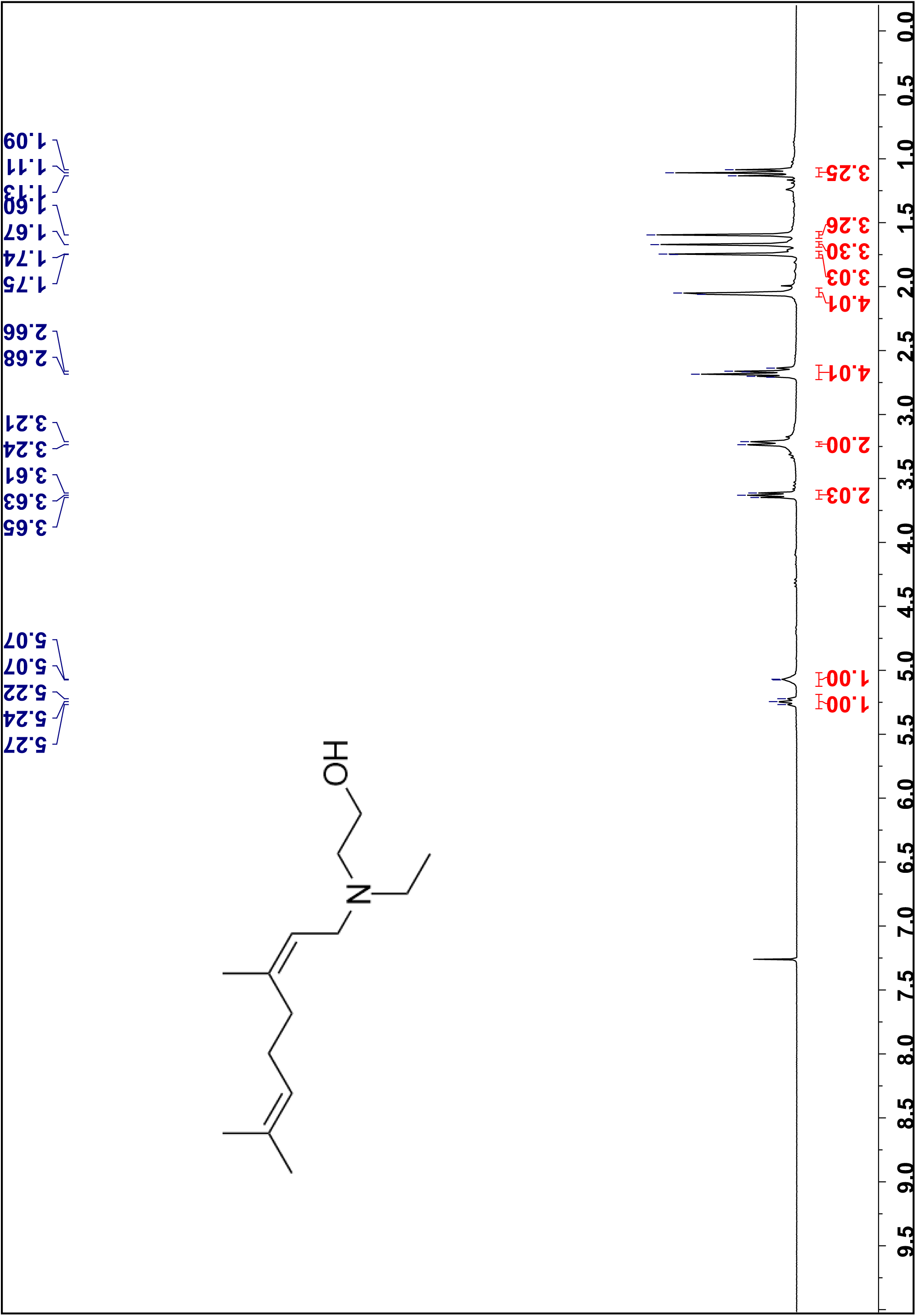
¹H NMR spectra of compound **3b** in CDCl_3_, 300 MHz.

**Figure 7.**
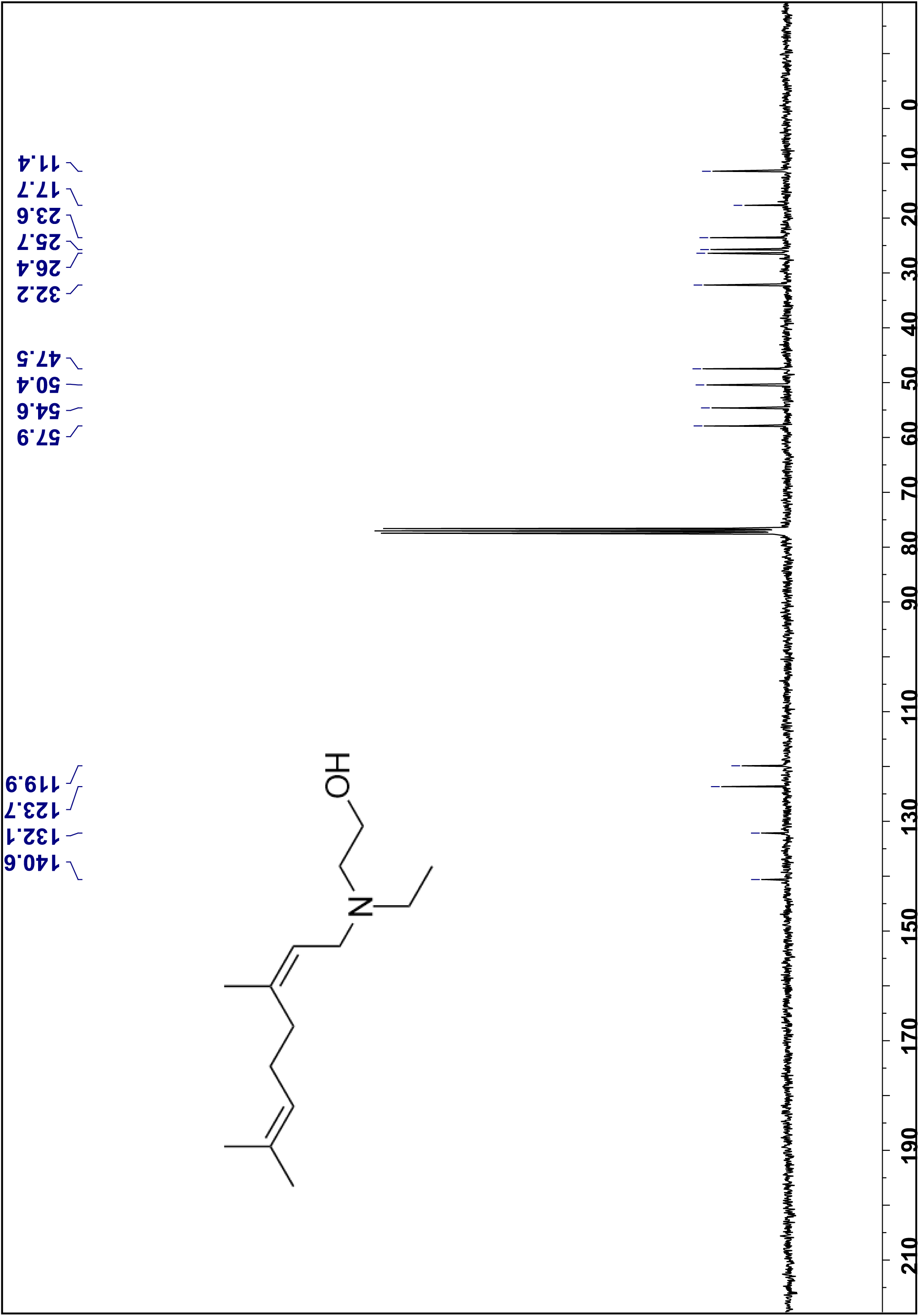
¹³C NMR spectra of compound **3b** in CDCl_3_, 75 MHz.

**Figure 8.**
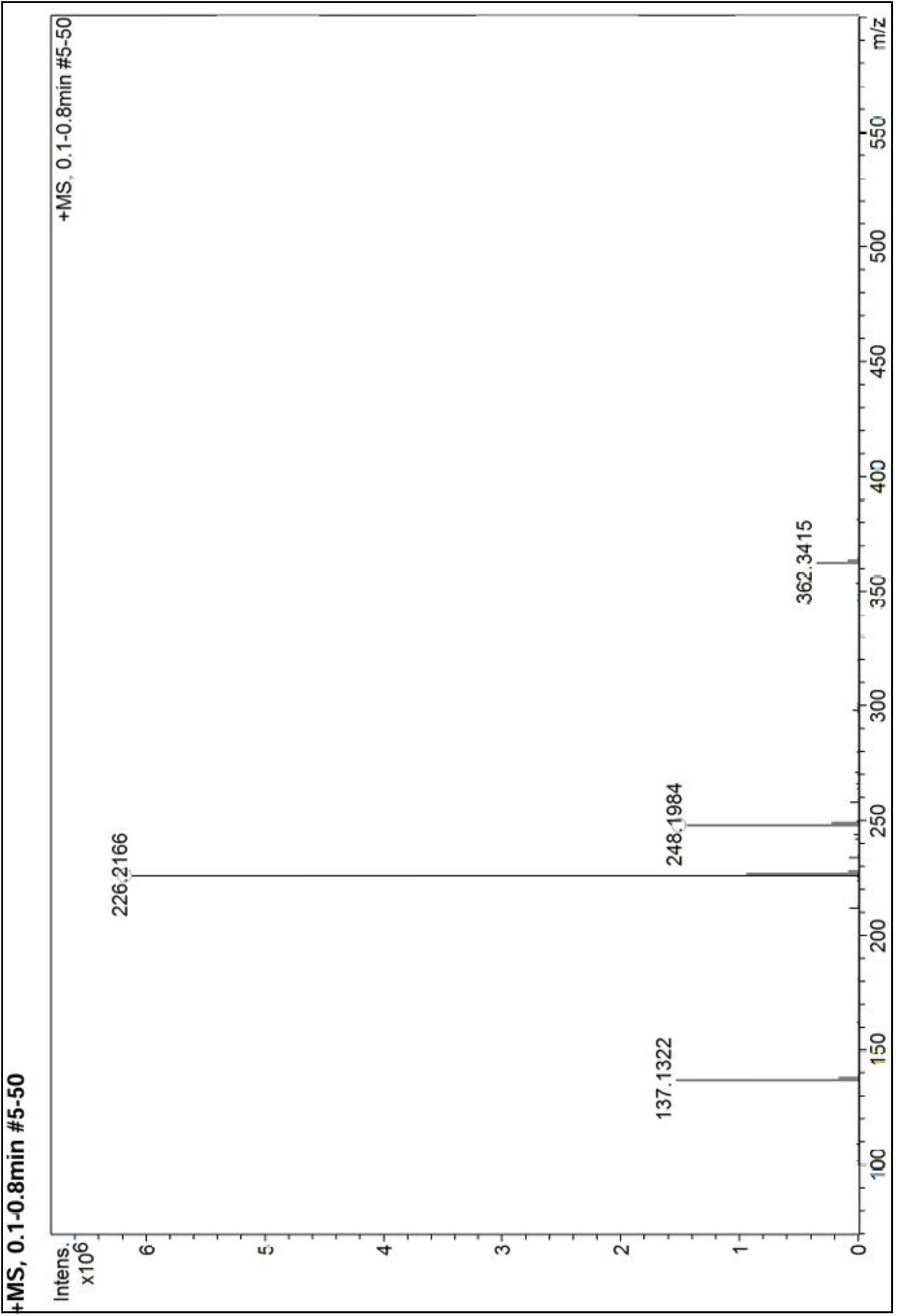
High Resolution Mass spectra of compound **3b**.

Compound 3c

**Figure 9.**
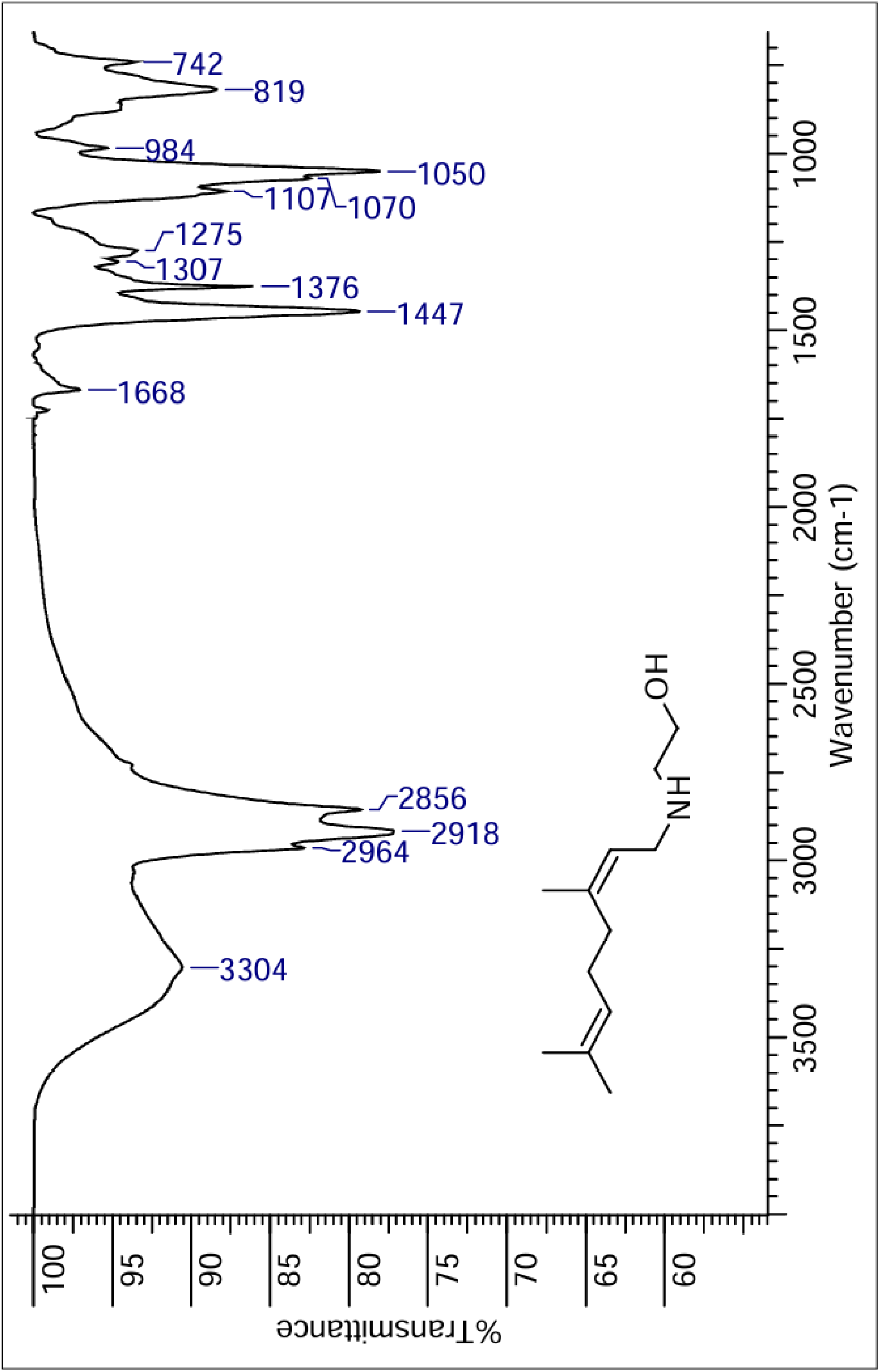
IR spectra of compound **3c**.

**Figure 10.**
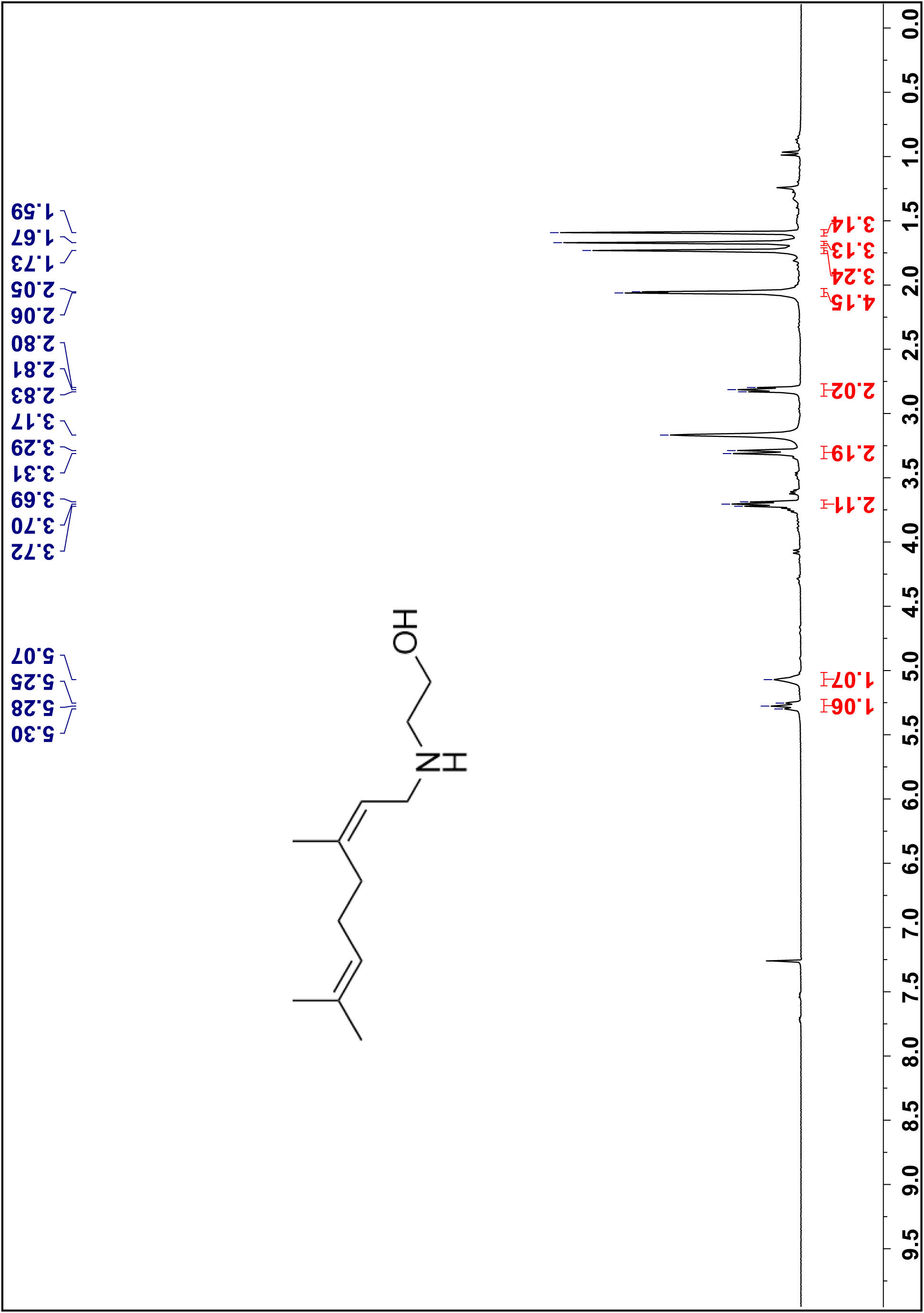
¹H NMR spectra of compound **3c** in CDCl_3_, 300 MHz.

**Figure 11.**
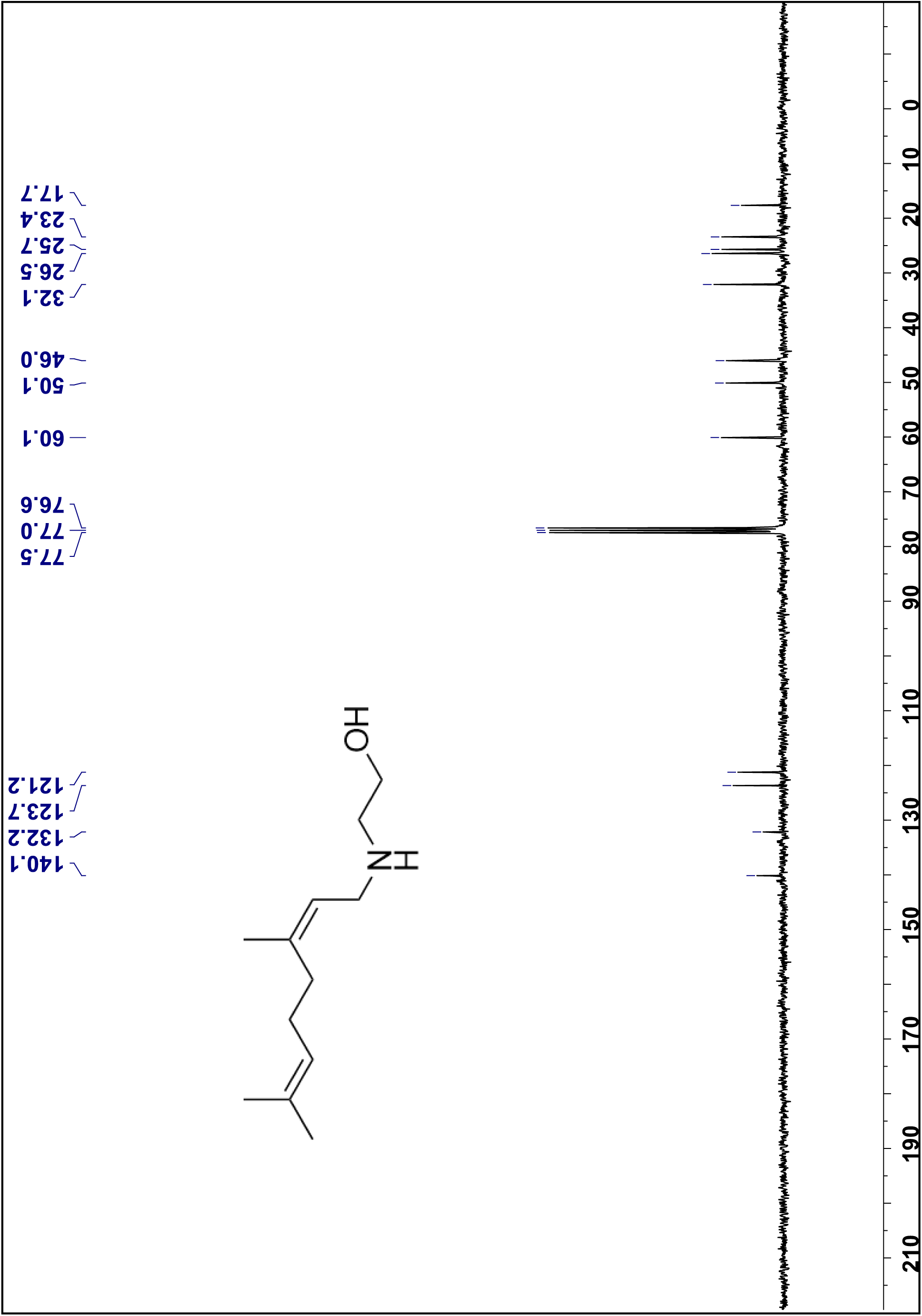
¹³C NMR spectra of compound **3c** in CDCl_3_, 75 MHz.

**Figure 12.**
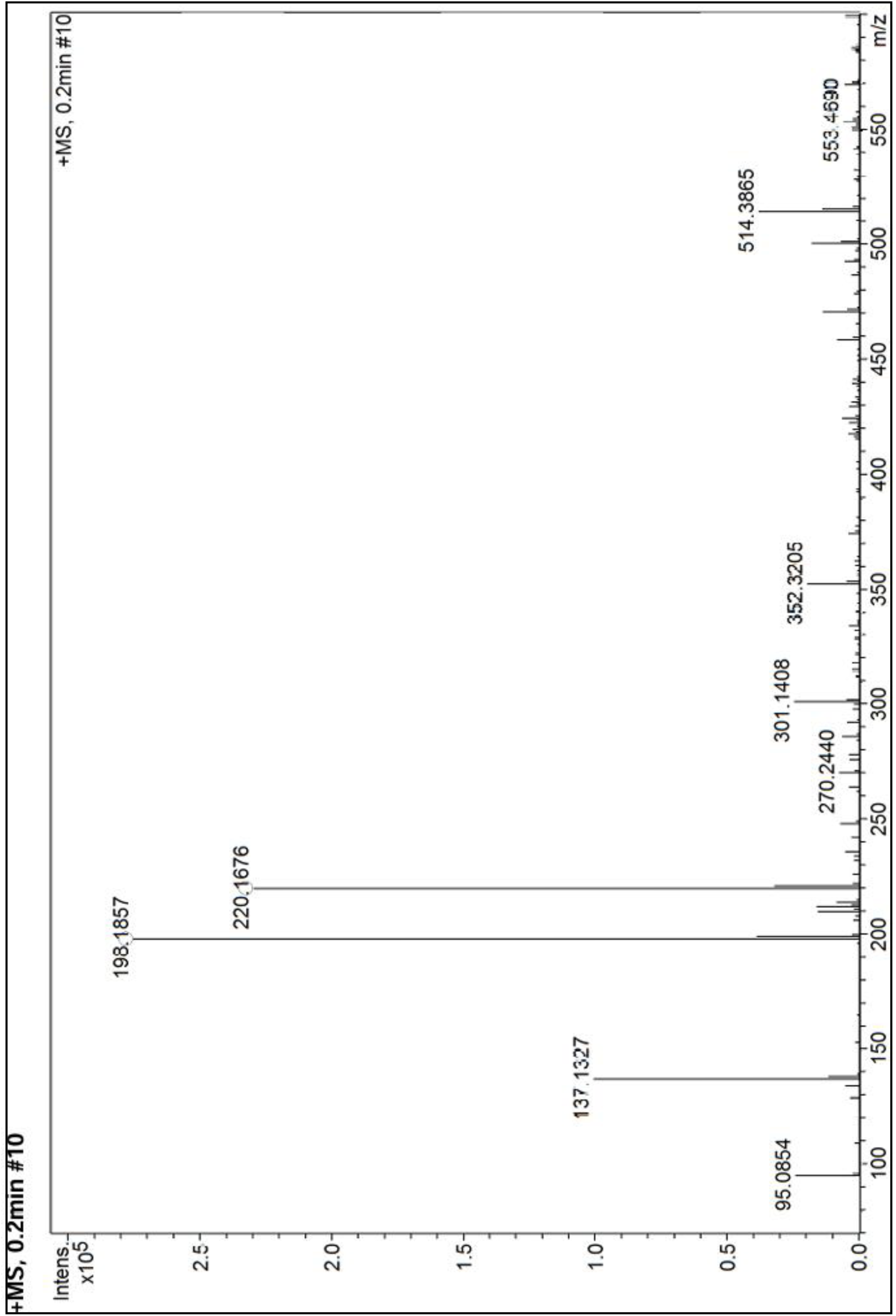
High Resolution Mass spectra of compound **3c**.

Compound 3d

**Figure 13.**
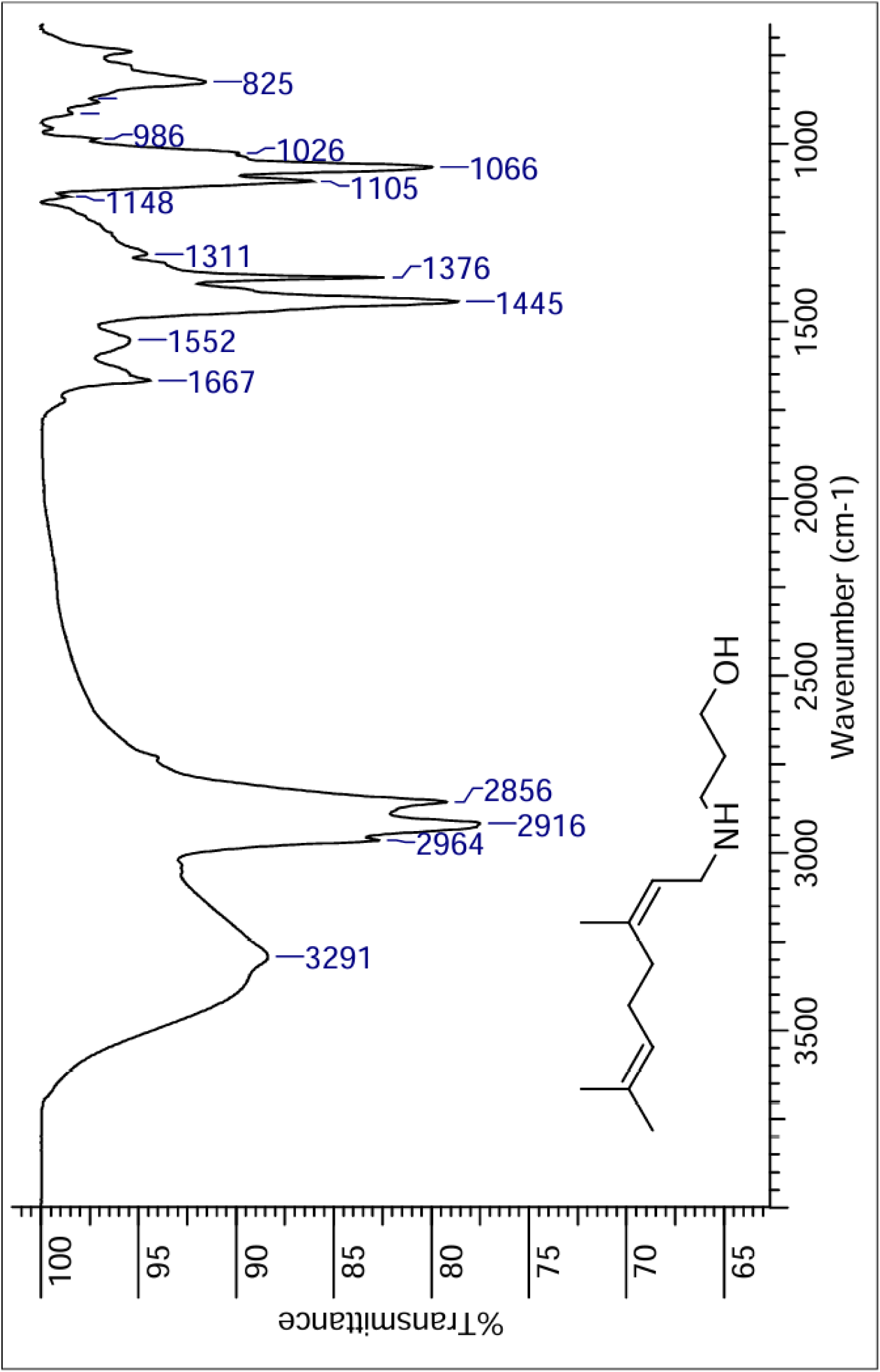
IR spectra of compound **3d**.

**Figure 14.**
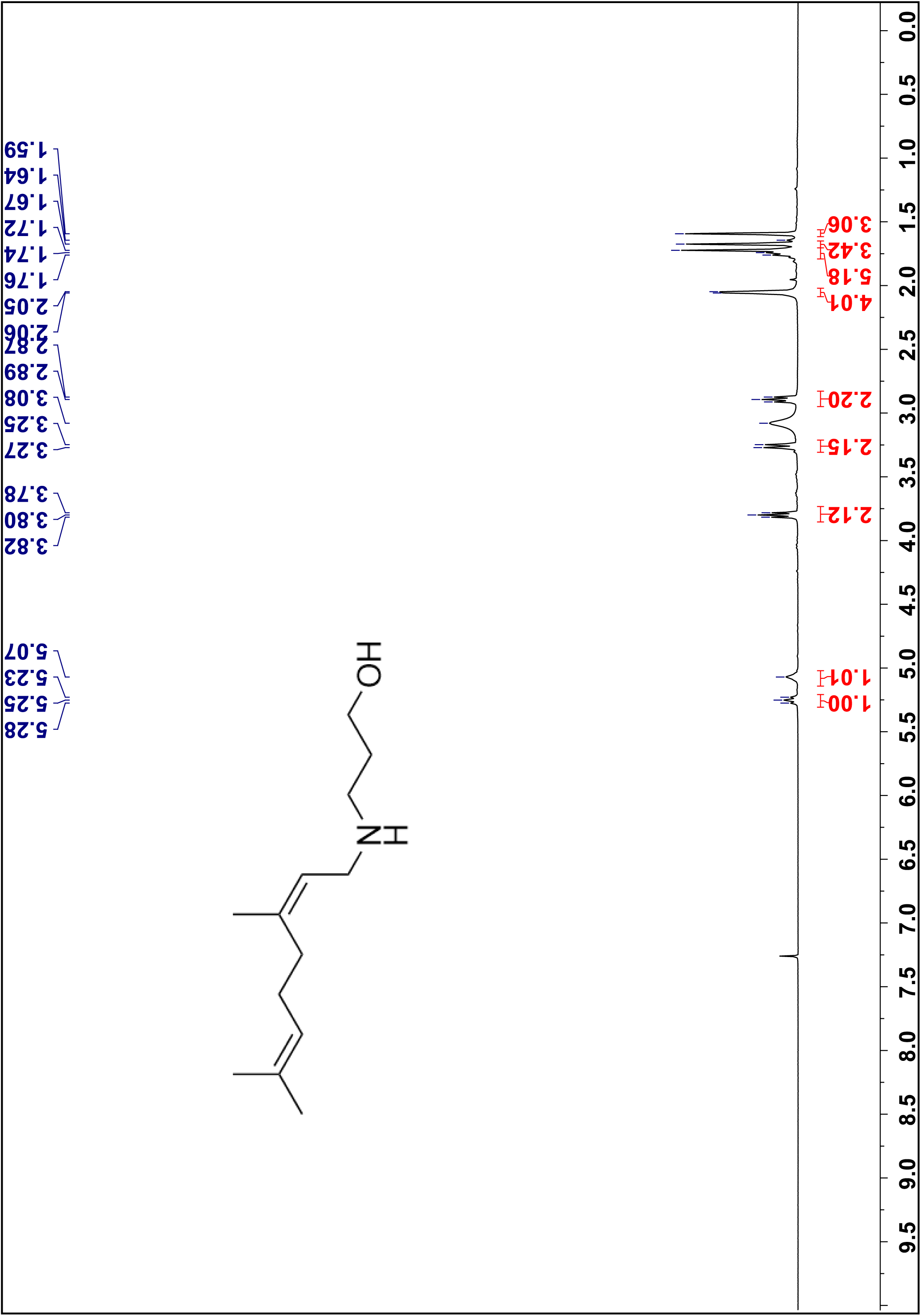
¹H NMR spectra of compound **3d** in CDCl_3_, 300 MHz.

**Figure 15.**
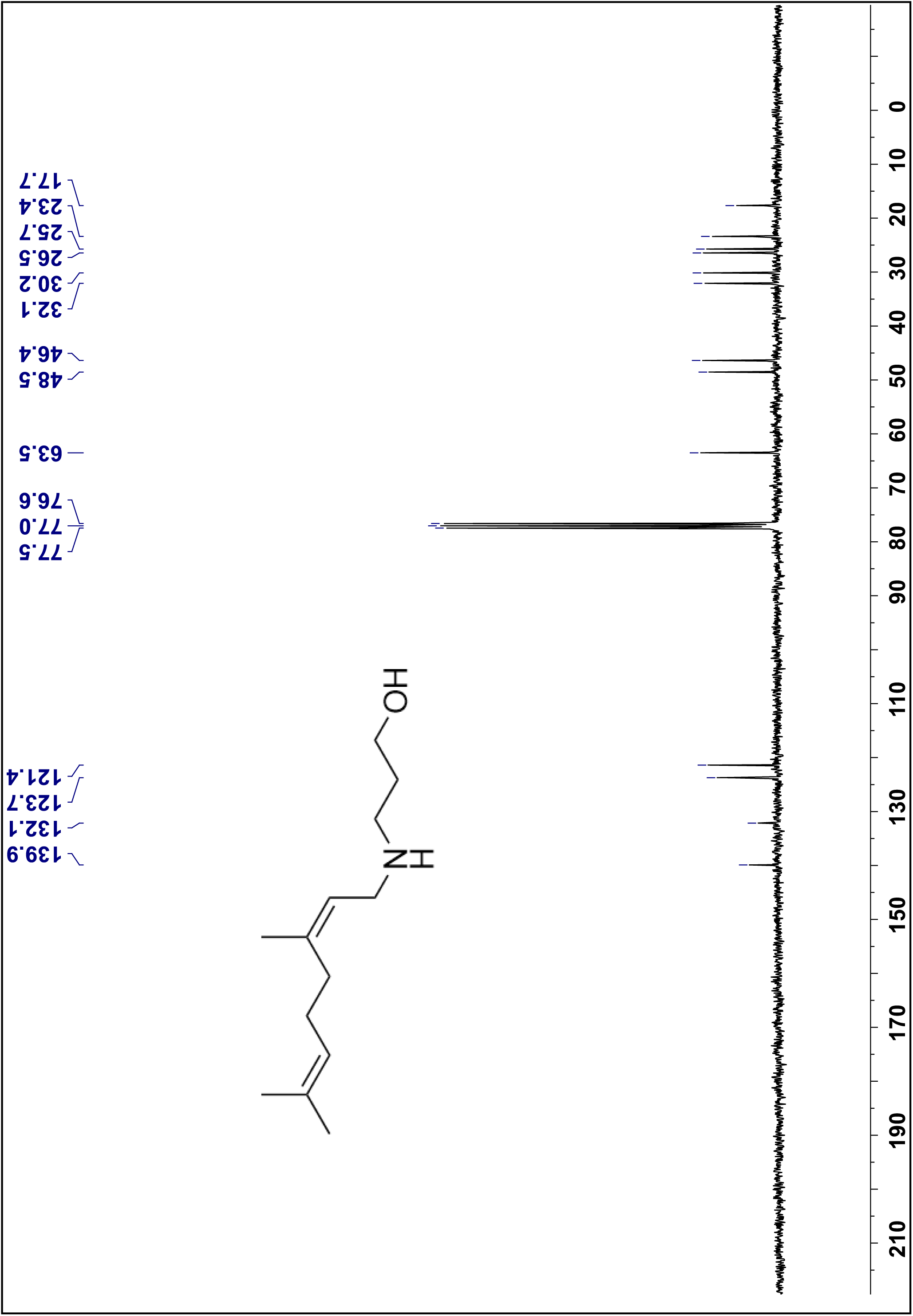
¹³C NMR spectra of compound **3d** in CDCl3, 75 MHz.

**Figure 16.**
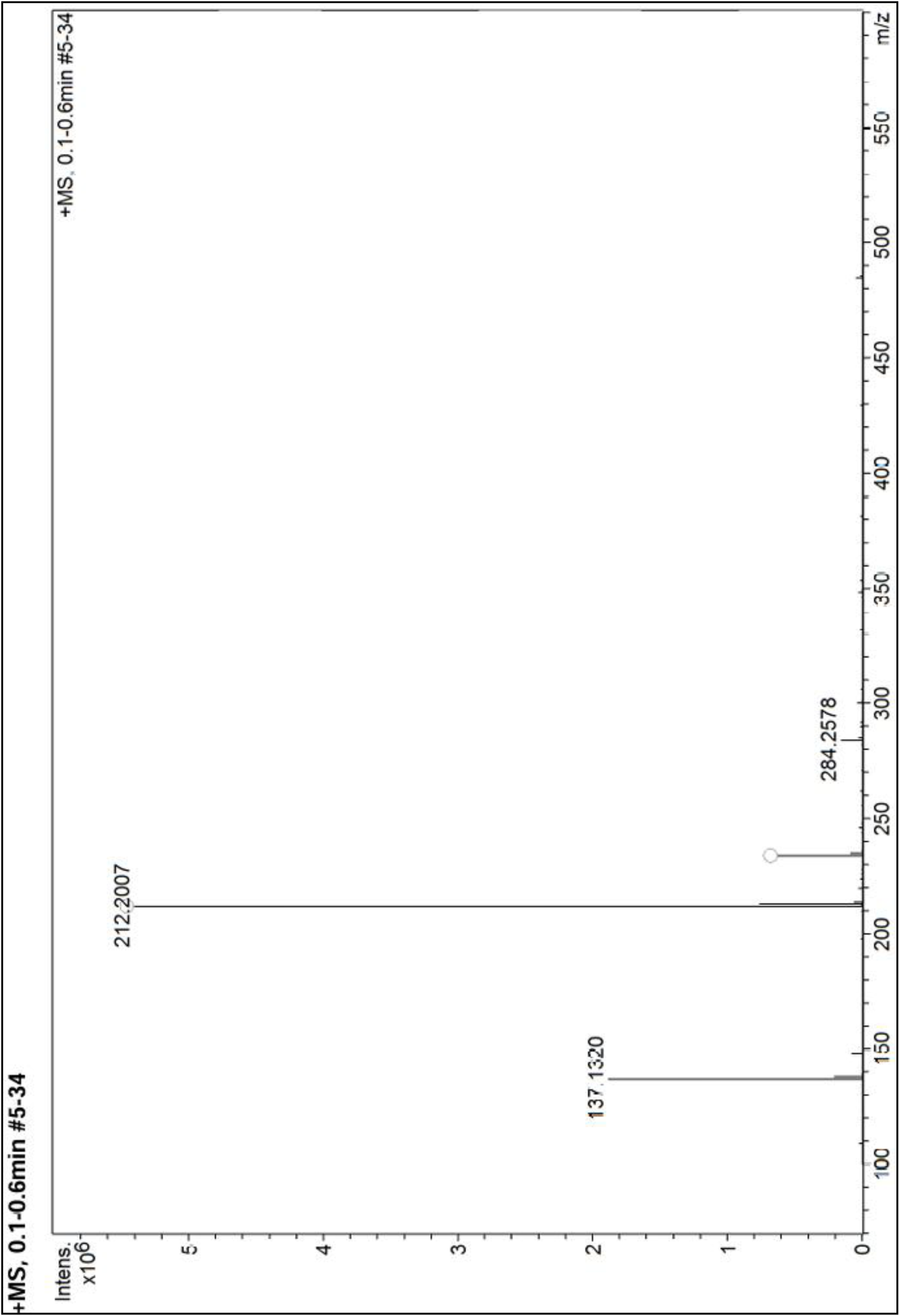
High Resolution Mass spectra of compound **3d**.

Compound 3e

**Figure 17.**
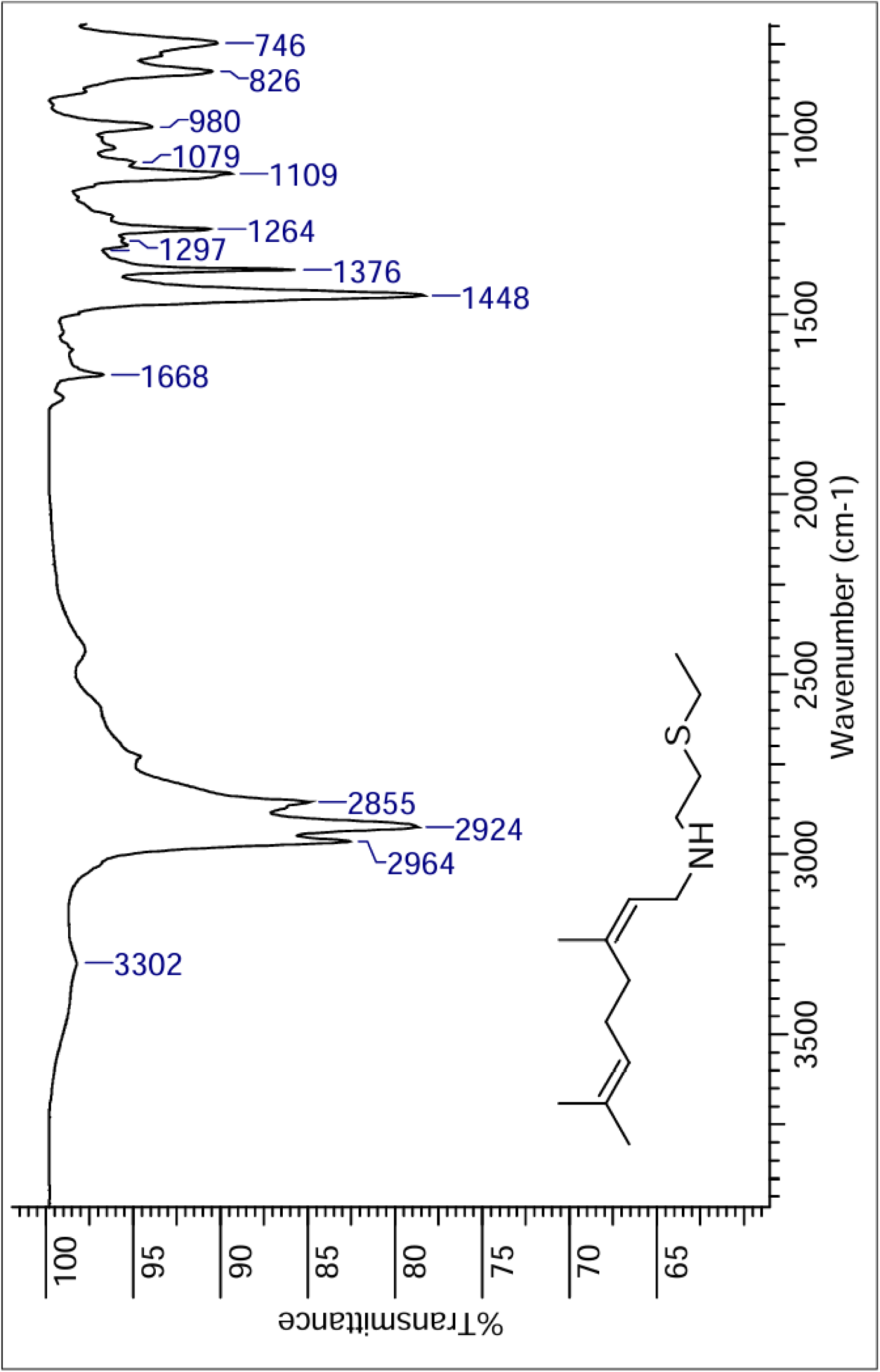
IR spectra of compound **3e**.

**Figure 18.**
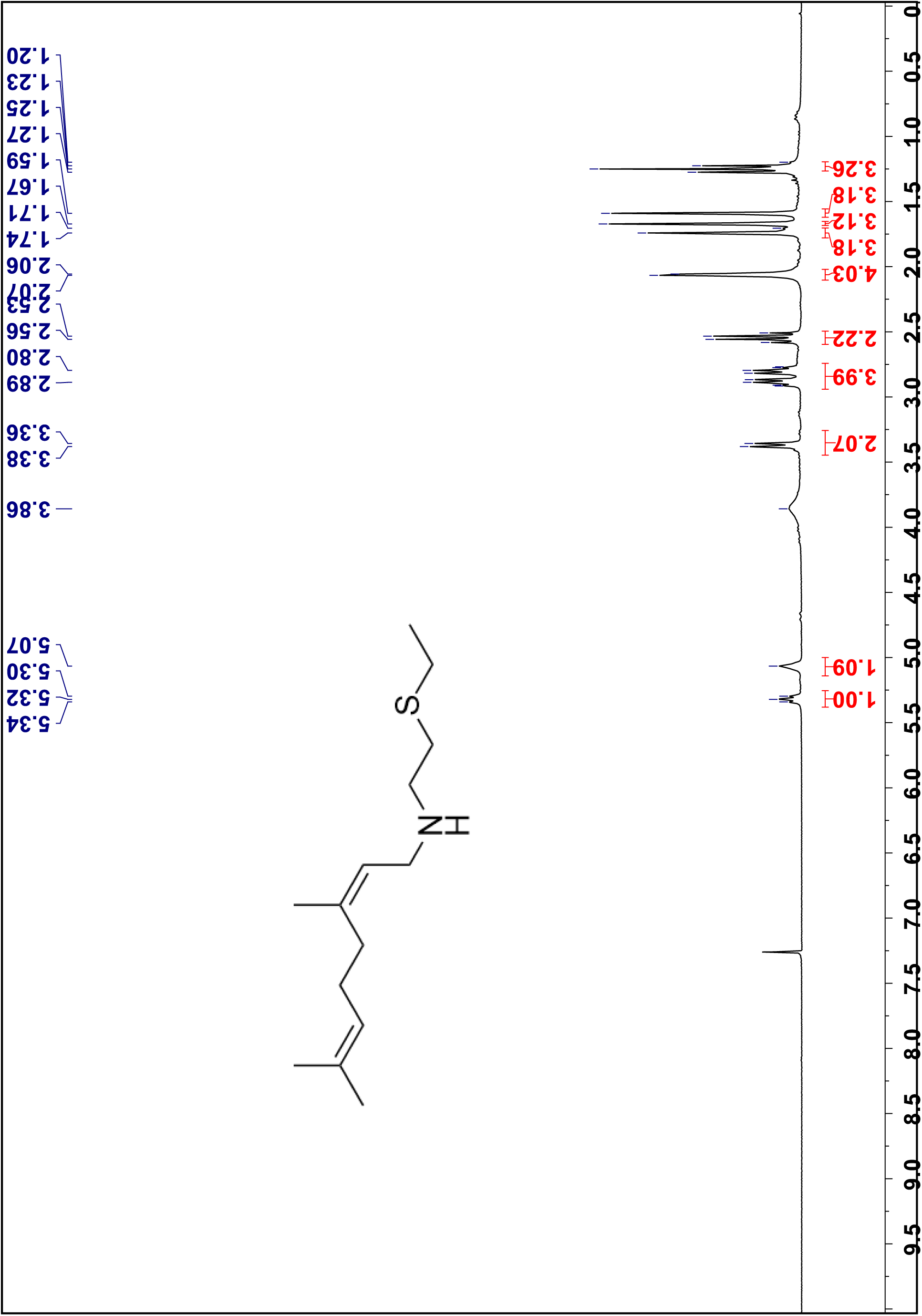
¹H NMR spectra of compound **3e** in CDCl_3_, 300 MHz.

**Figure 19.**
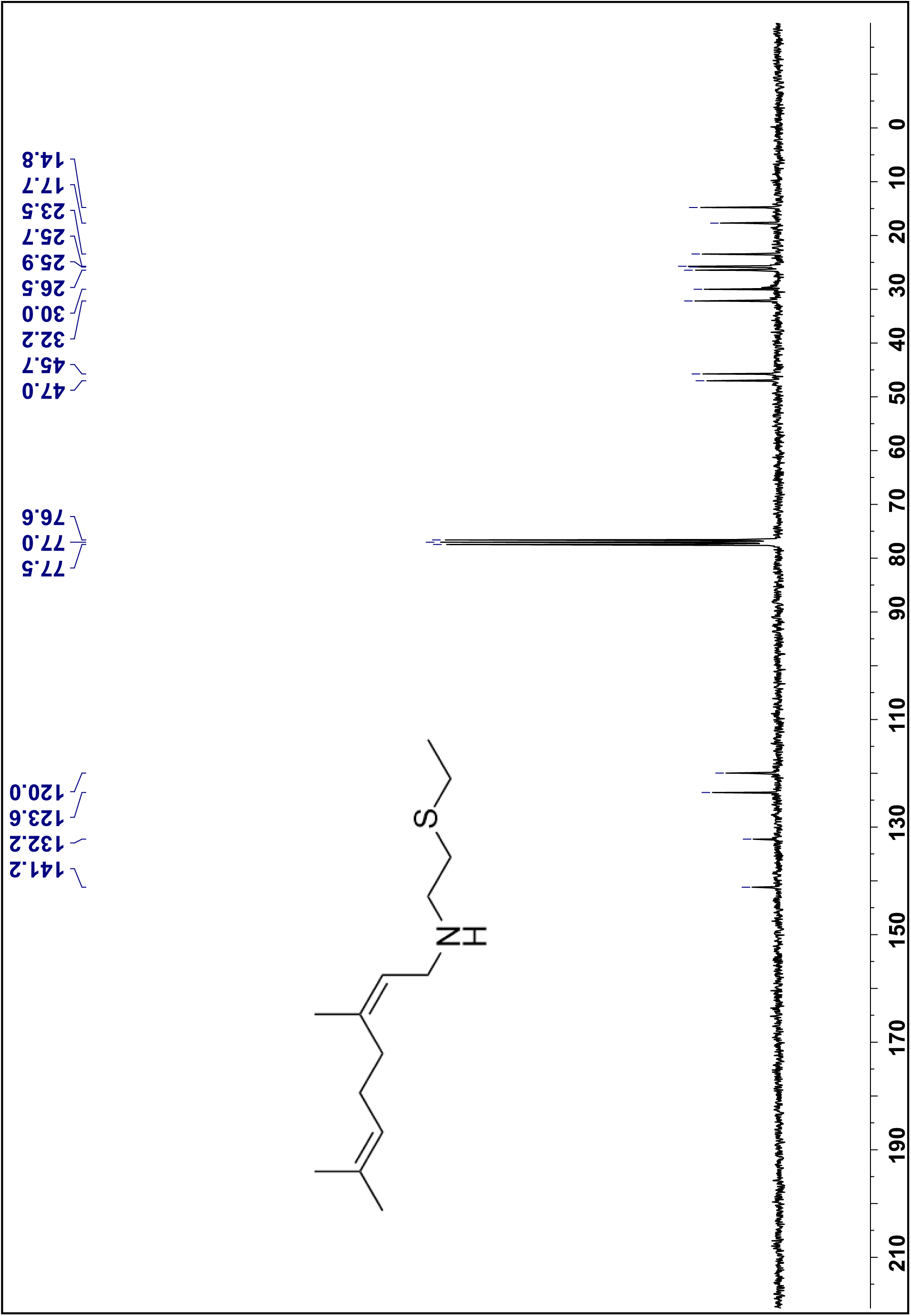
¹³C NMR spectra of compound **3e** in CDCl_3_, 75 MHz.

**Figure 20.**
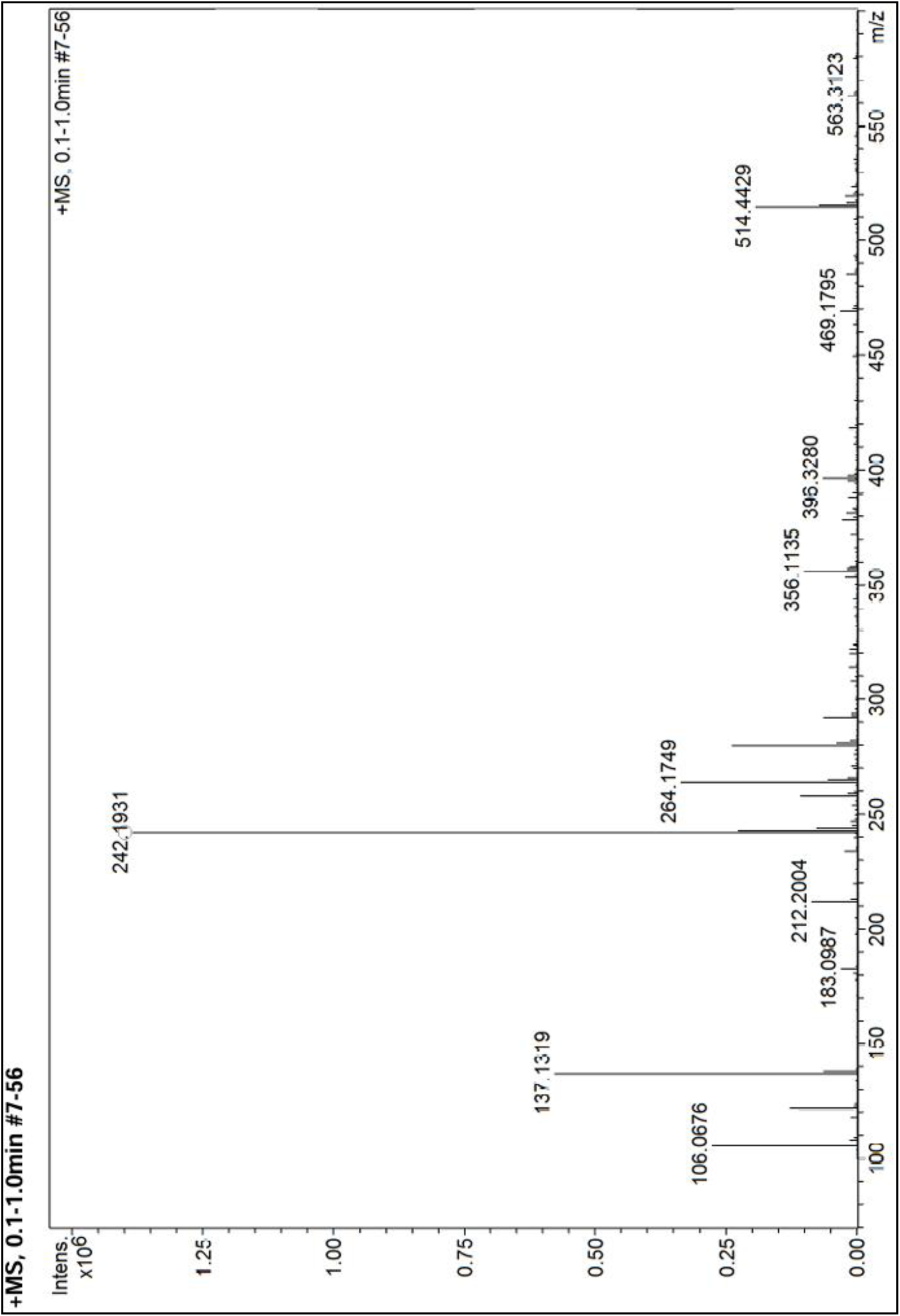
High Resolution Mass spectra of compound **3e**.

Compound 3f

**Figure 21.**
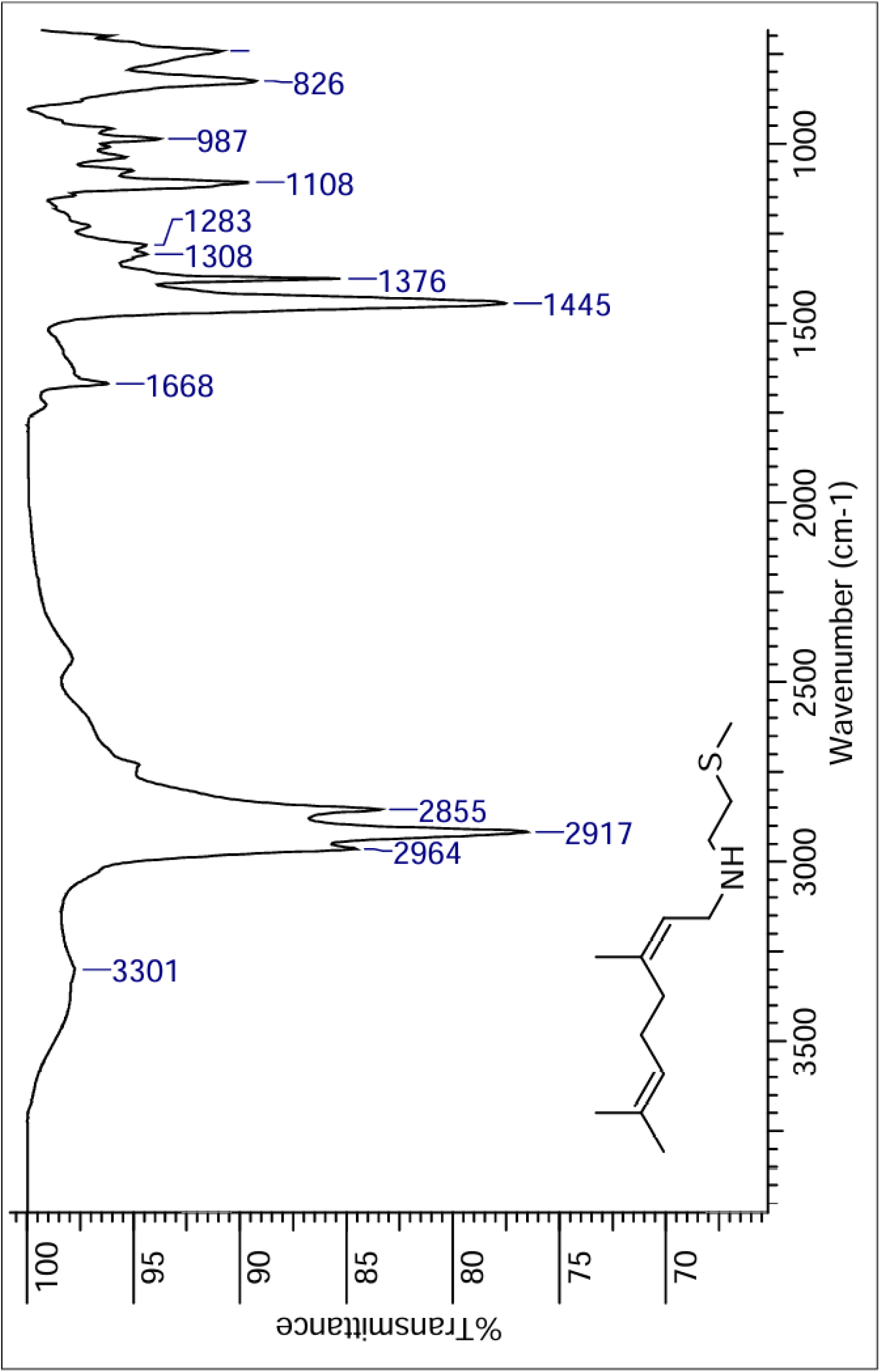
IR spectra of compound **3f**.

**Figure 22.**
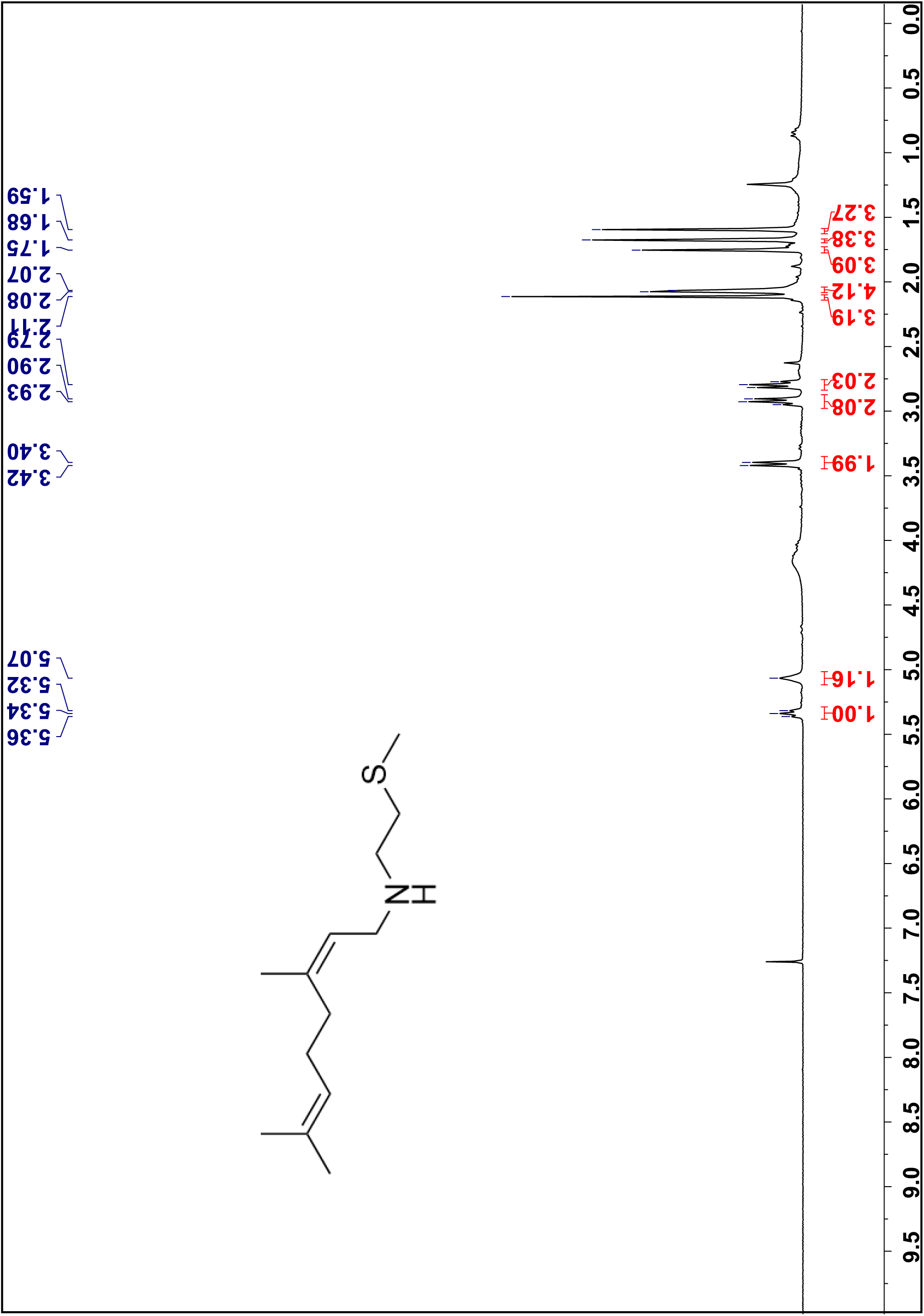
¹H NMR spectra of compound **3f** in CDCl3, 300 MHz.

**Figure 23.**
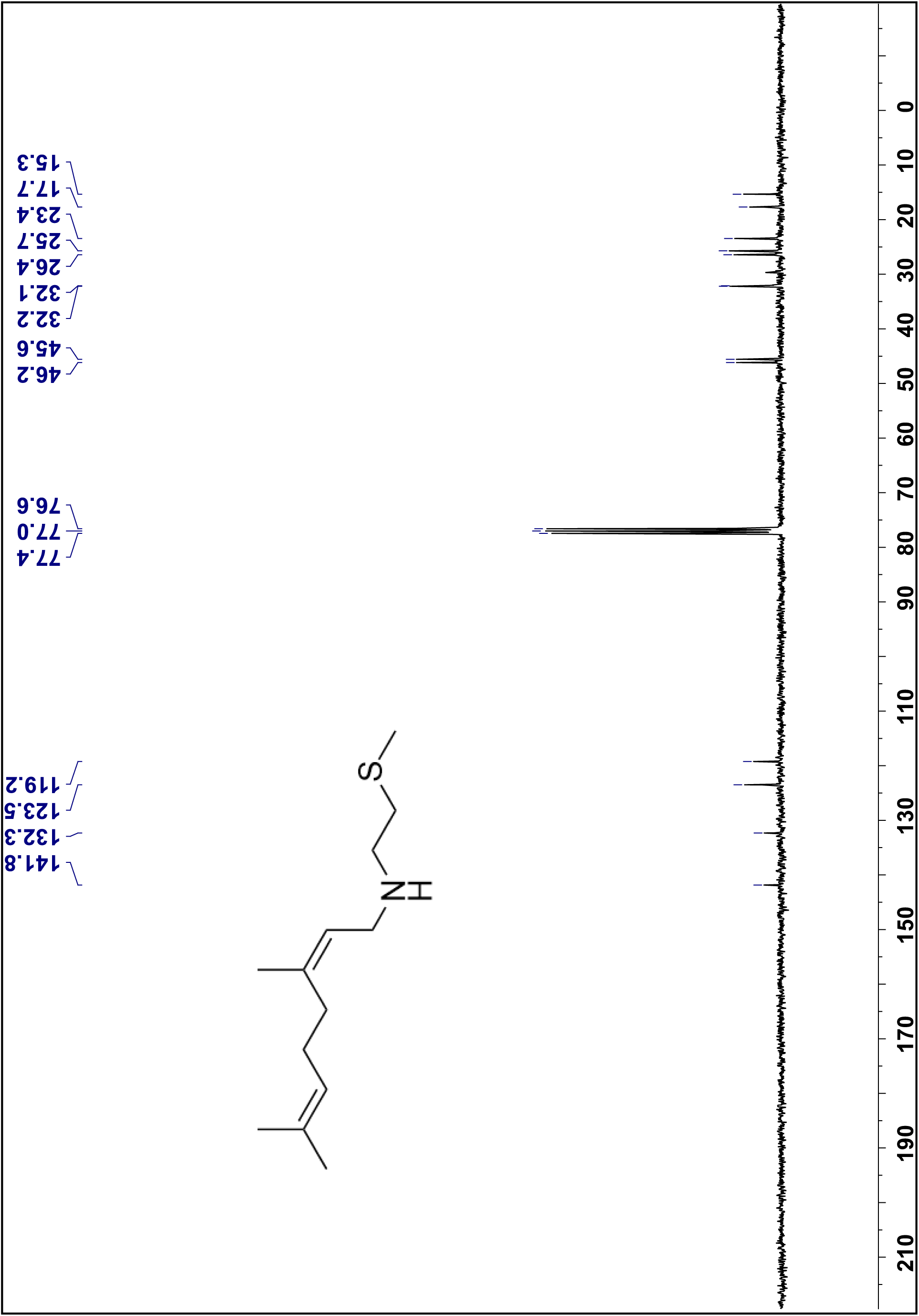
¹³C NMR spectra of compound **3f** in CDCl3, 75 MHz.

**Figure 24.**
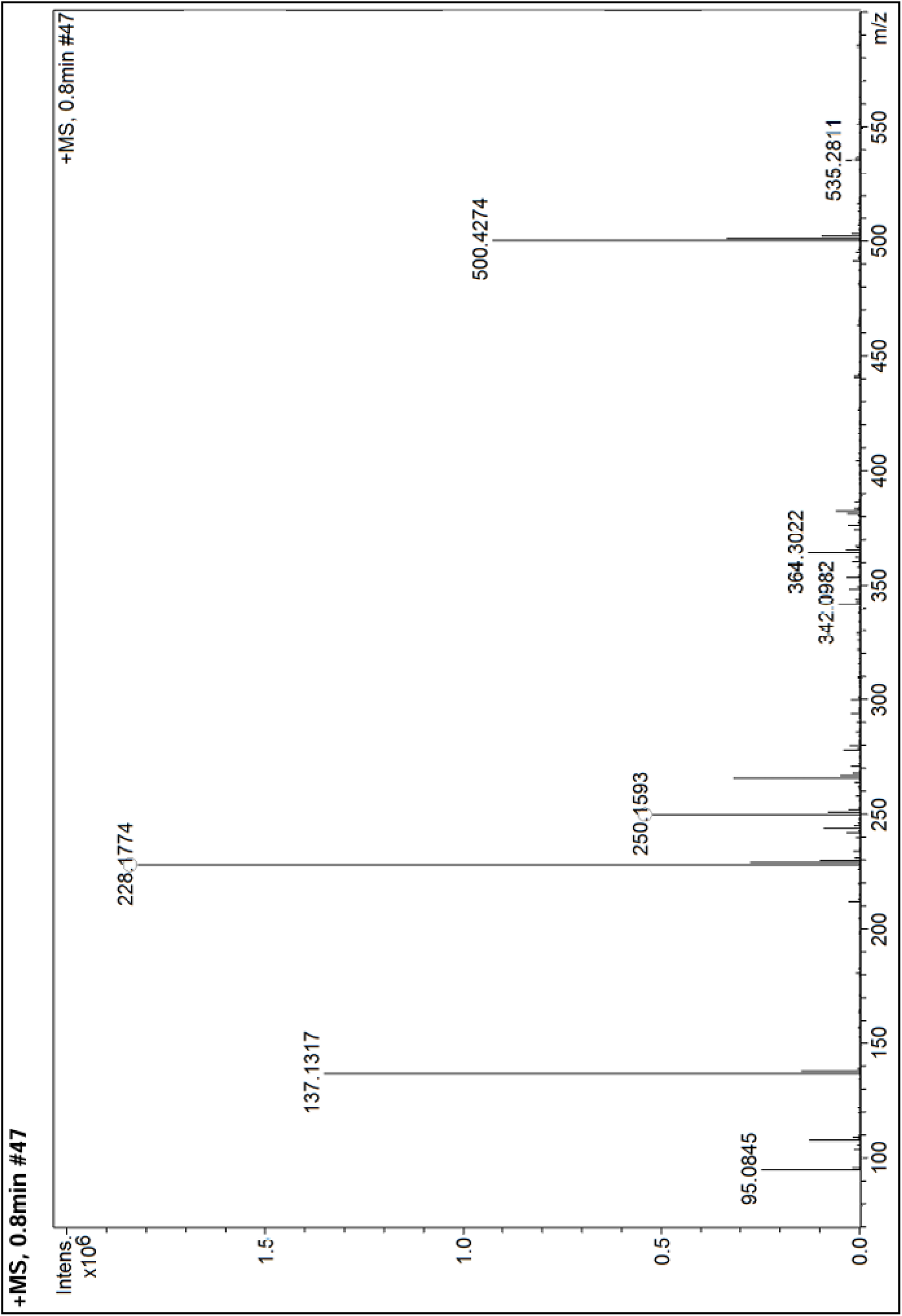
High Resolution Mass spectra of compound **3f**.

Compound 3g

**Figure 25.**
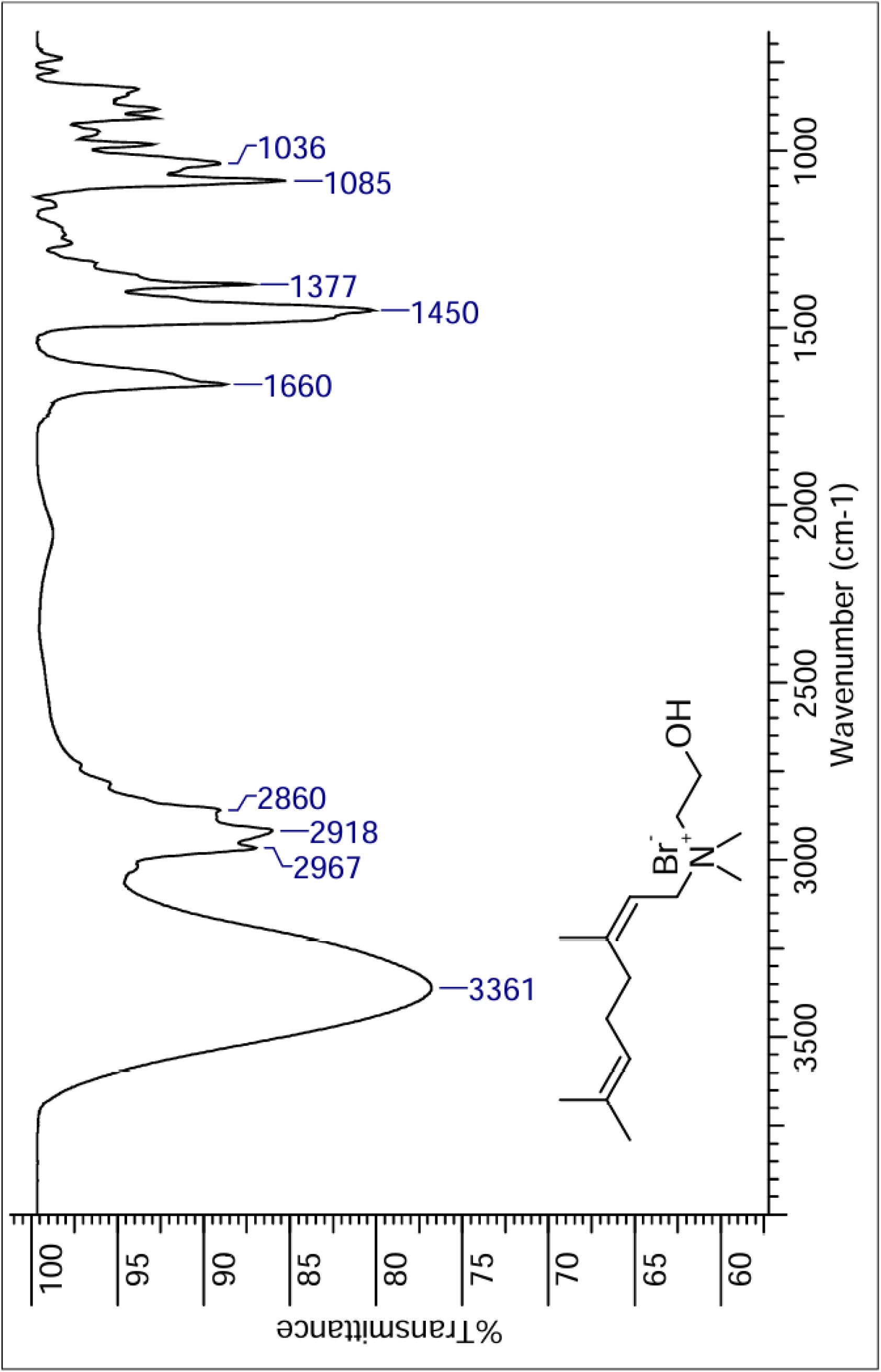
IR spectra of compound **3g**.

**Figure 26.**
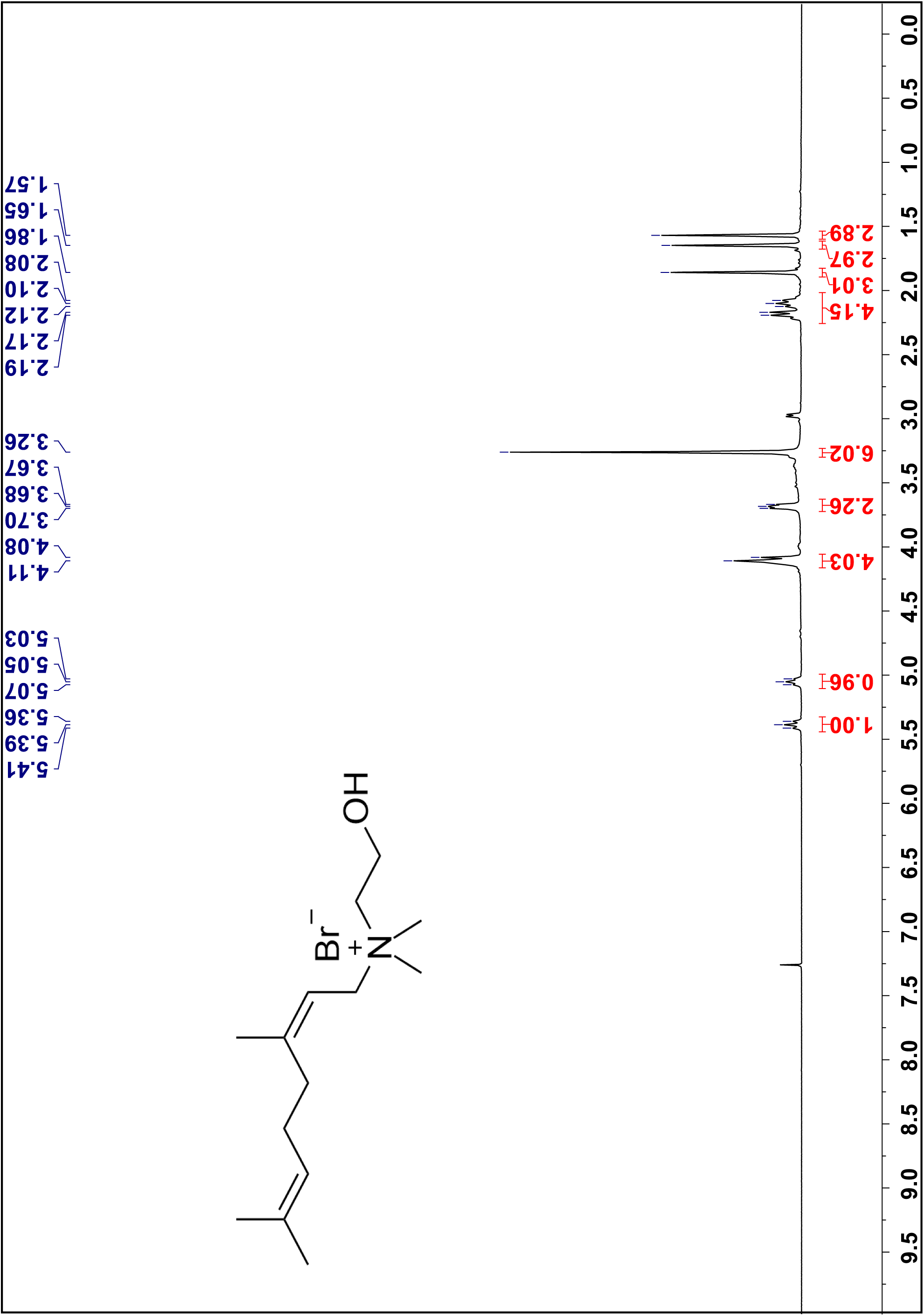
¹H NMR spectra of compound **3g** in CDCl_3_, 300 MHz.

**Figure 27.**
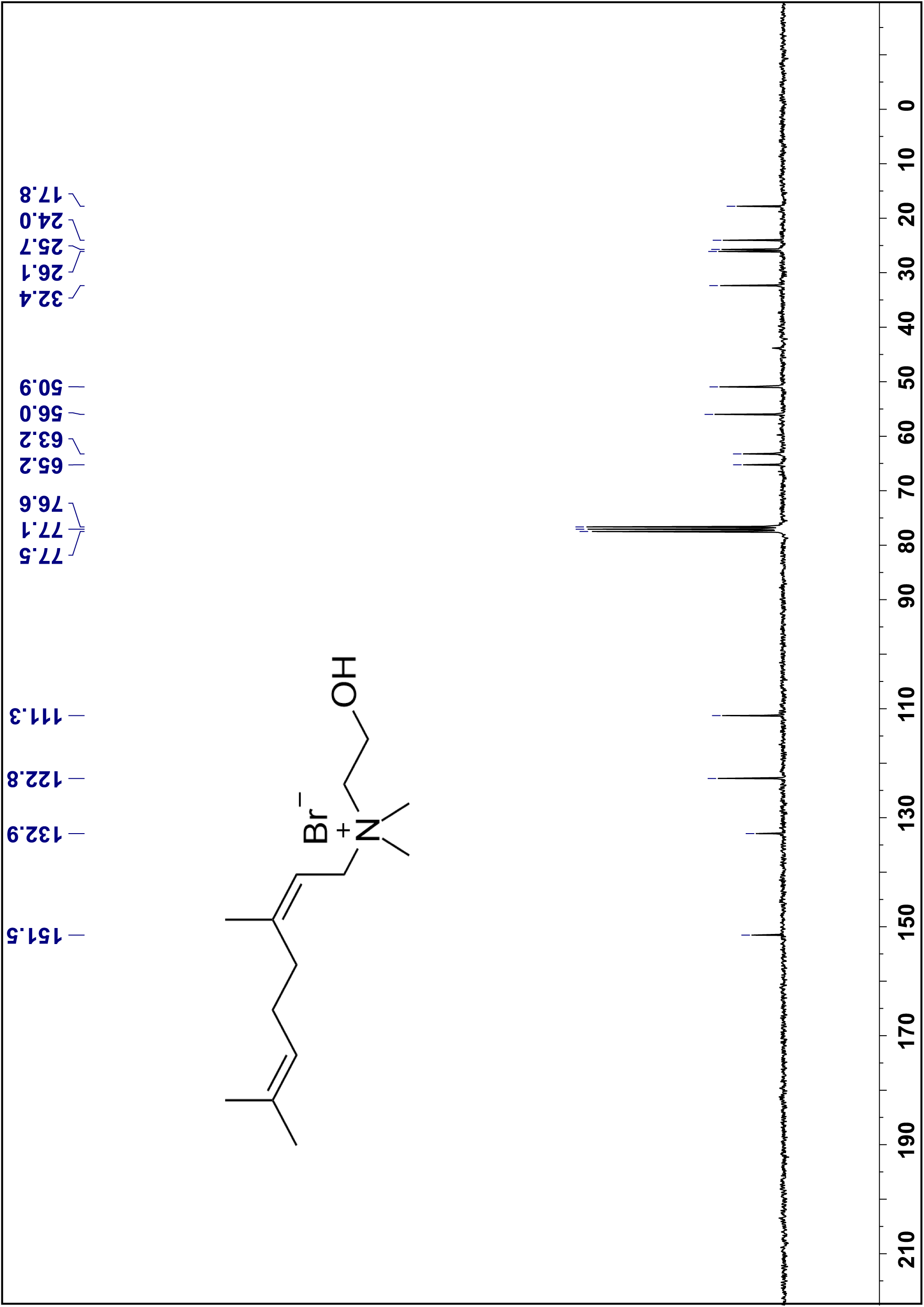
¹³C NMR spectra of compound **3g** in CDCl_3_, 300 MHz.

**Figure 28.**
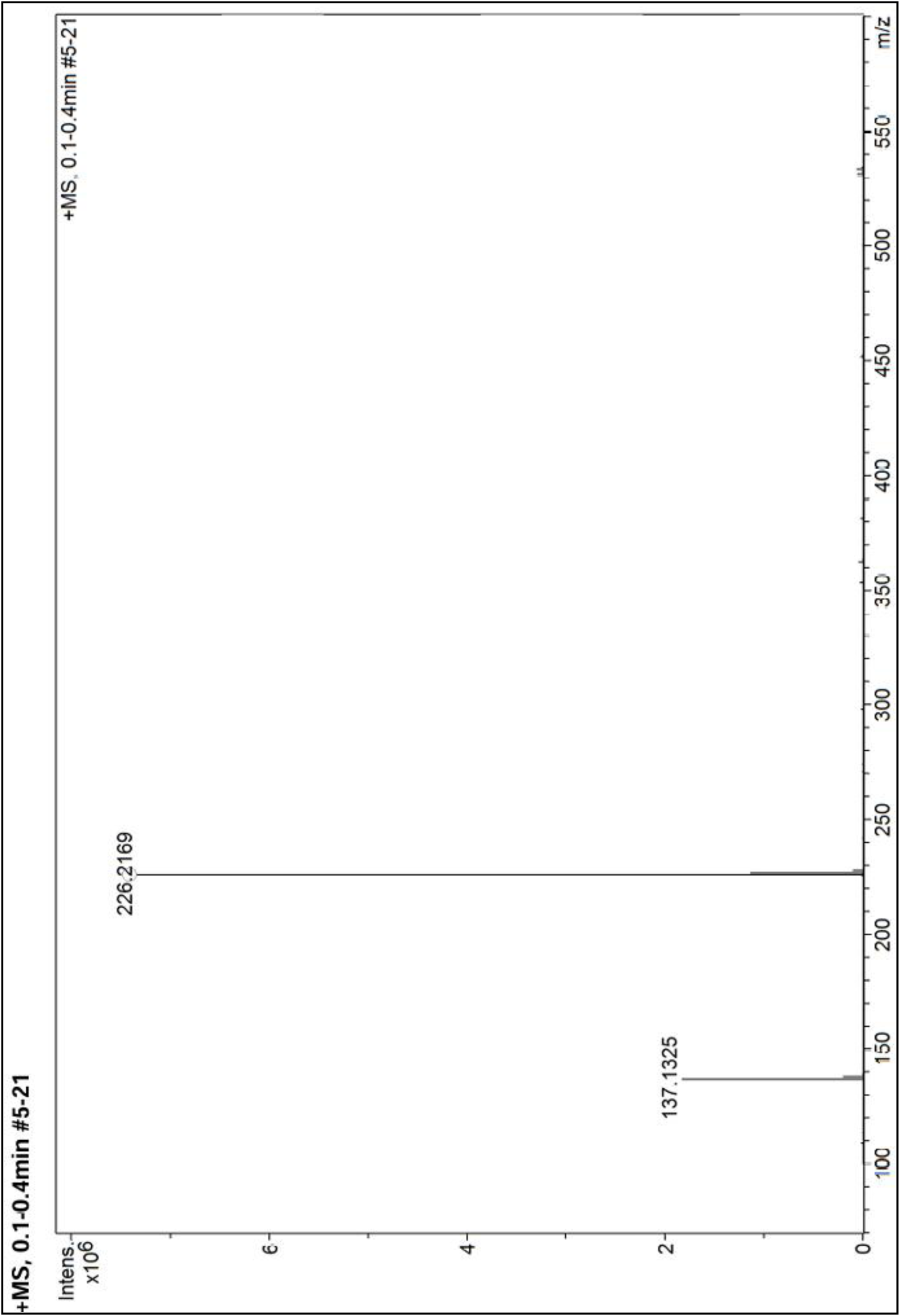
High Resolution Mass spectra of compound **3g**.

Compound 3h

**Figure 29.**
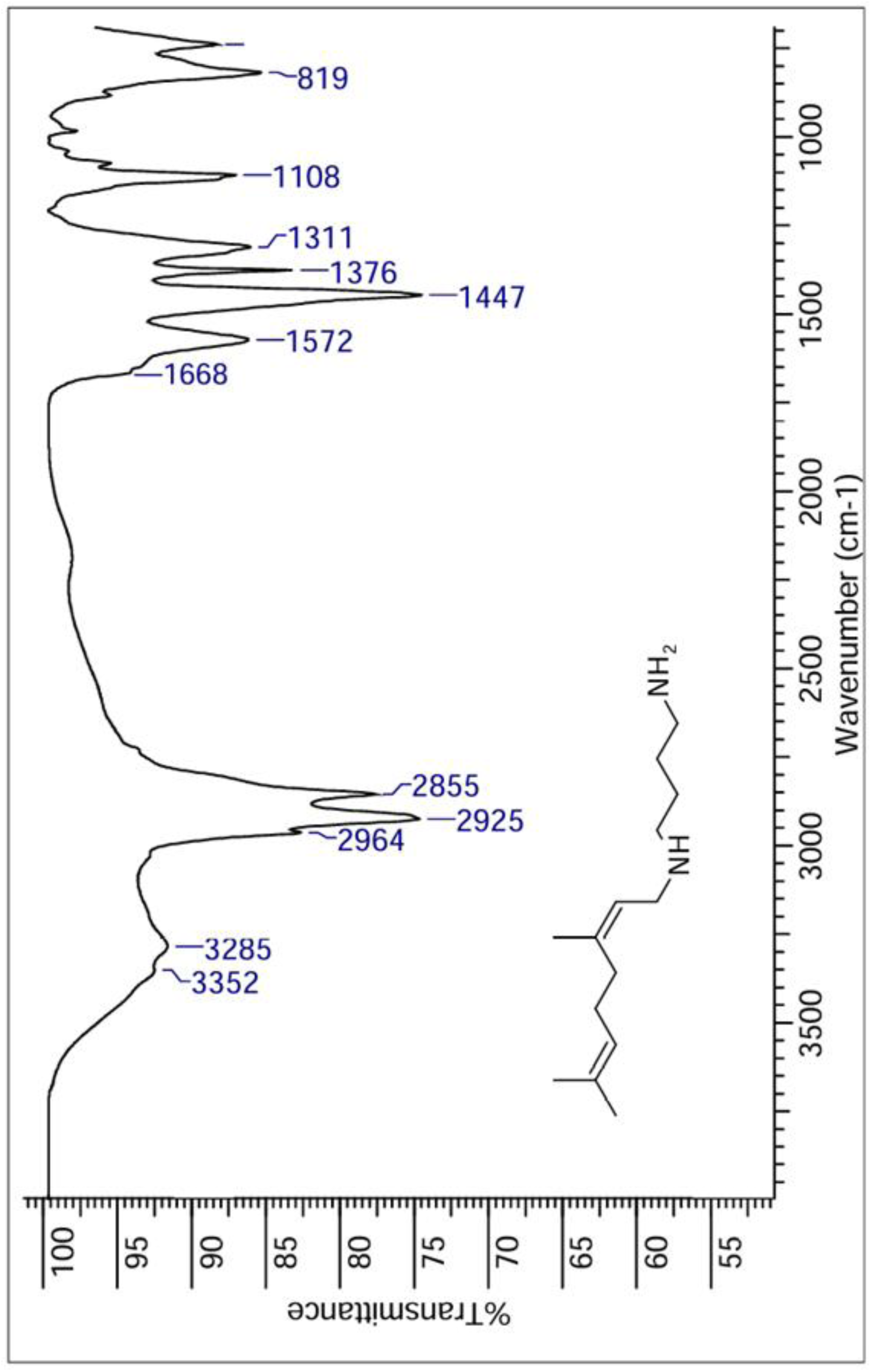
IR spectra of compound **3h**.

**Figure 30.**
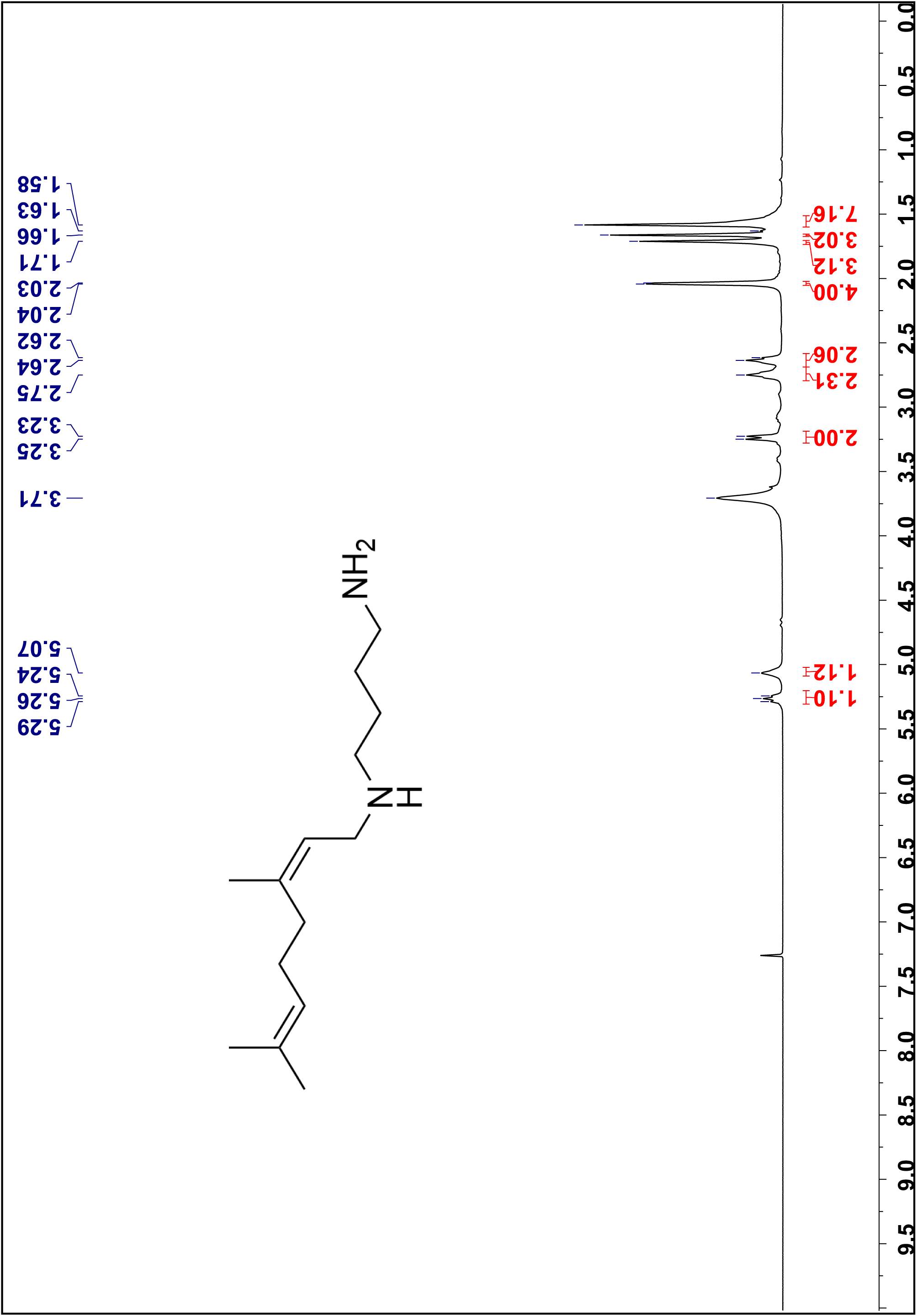
¹H NMR spectra of compound **3h** in CDCl_3_, 300 MHz.

**Figure 31.**
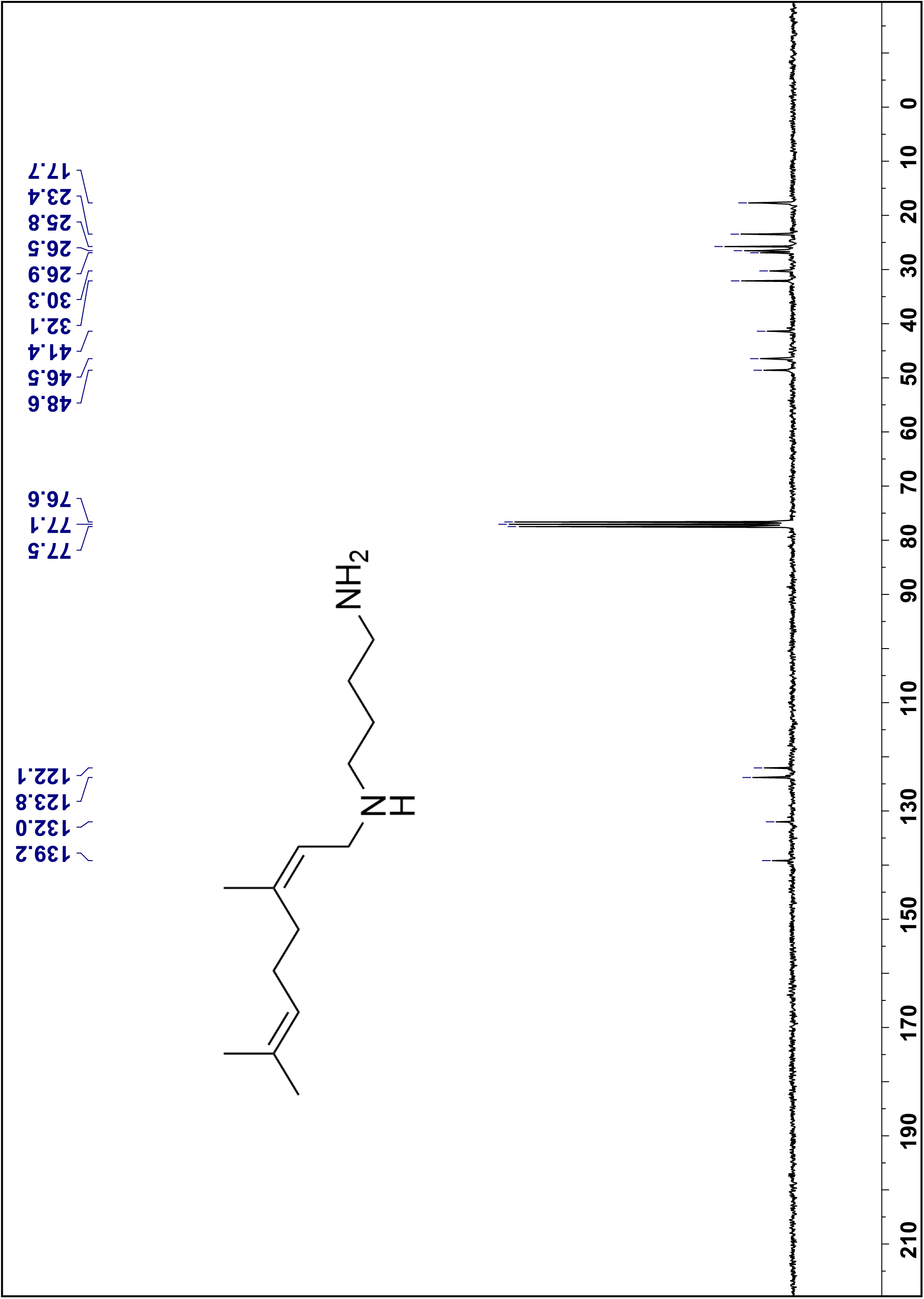
¹³C NMR spectra of compound **3h** in CDCl_3_, 75 MHz.

**Figure 32.**
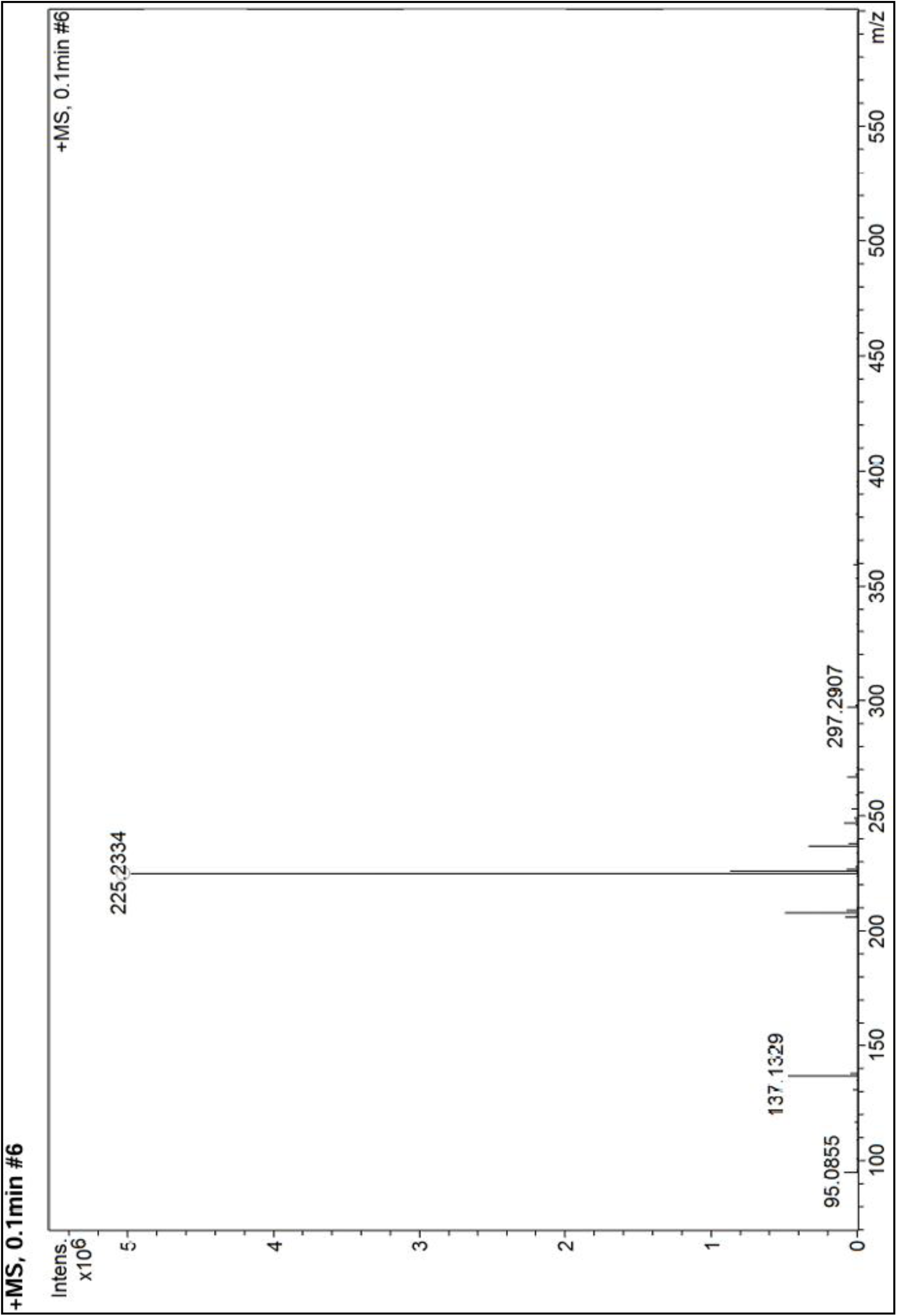
High Resolution Mass spectra of compound **3g**.

Compound 5a

**Figure 33.**
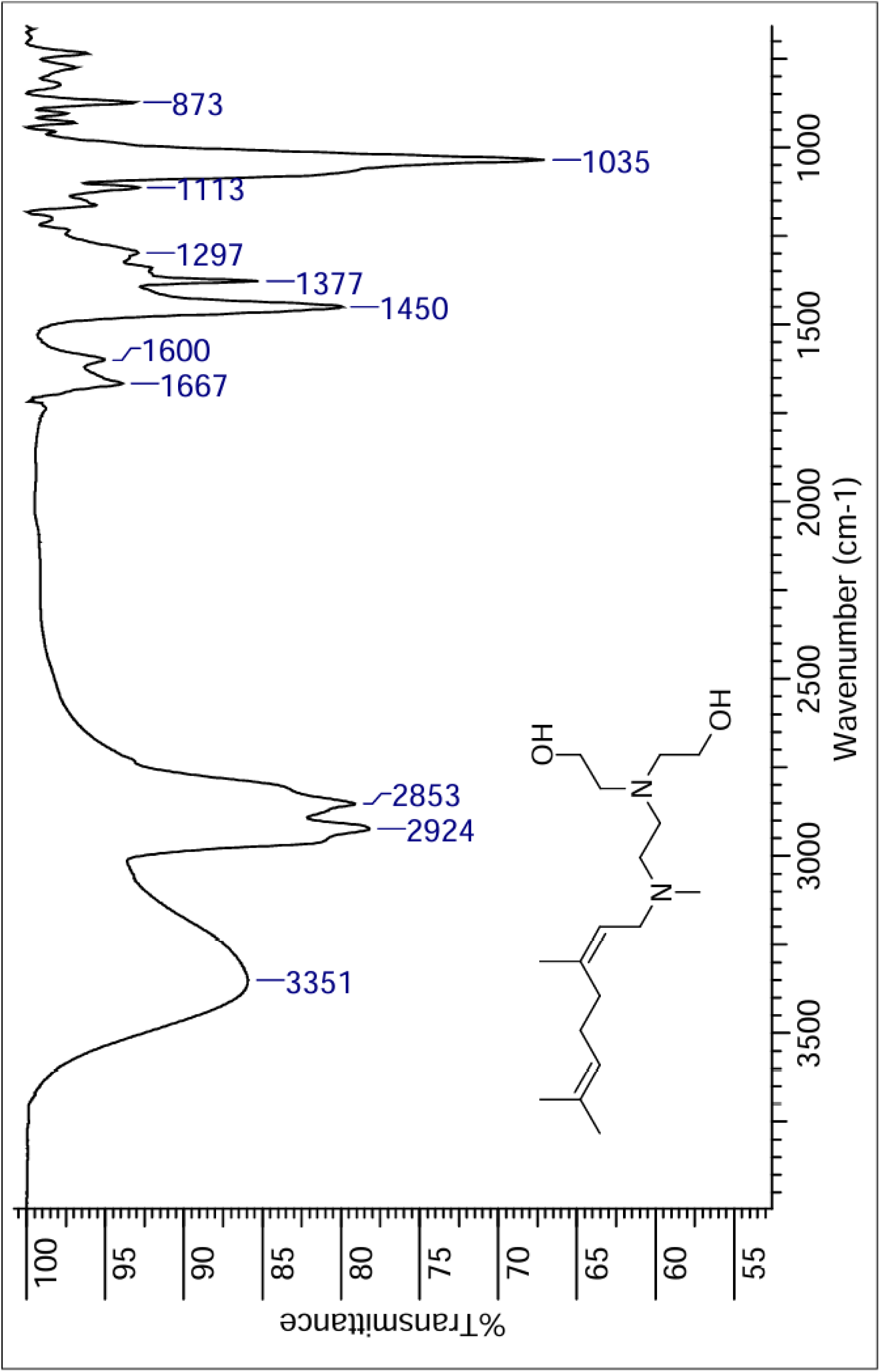
IR spectra of compound **5a**.

**Figure 34.**
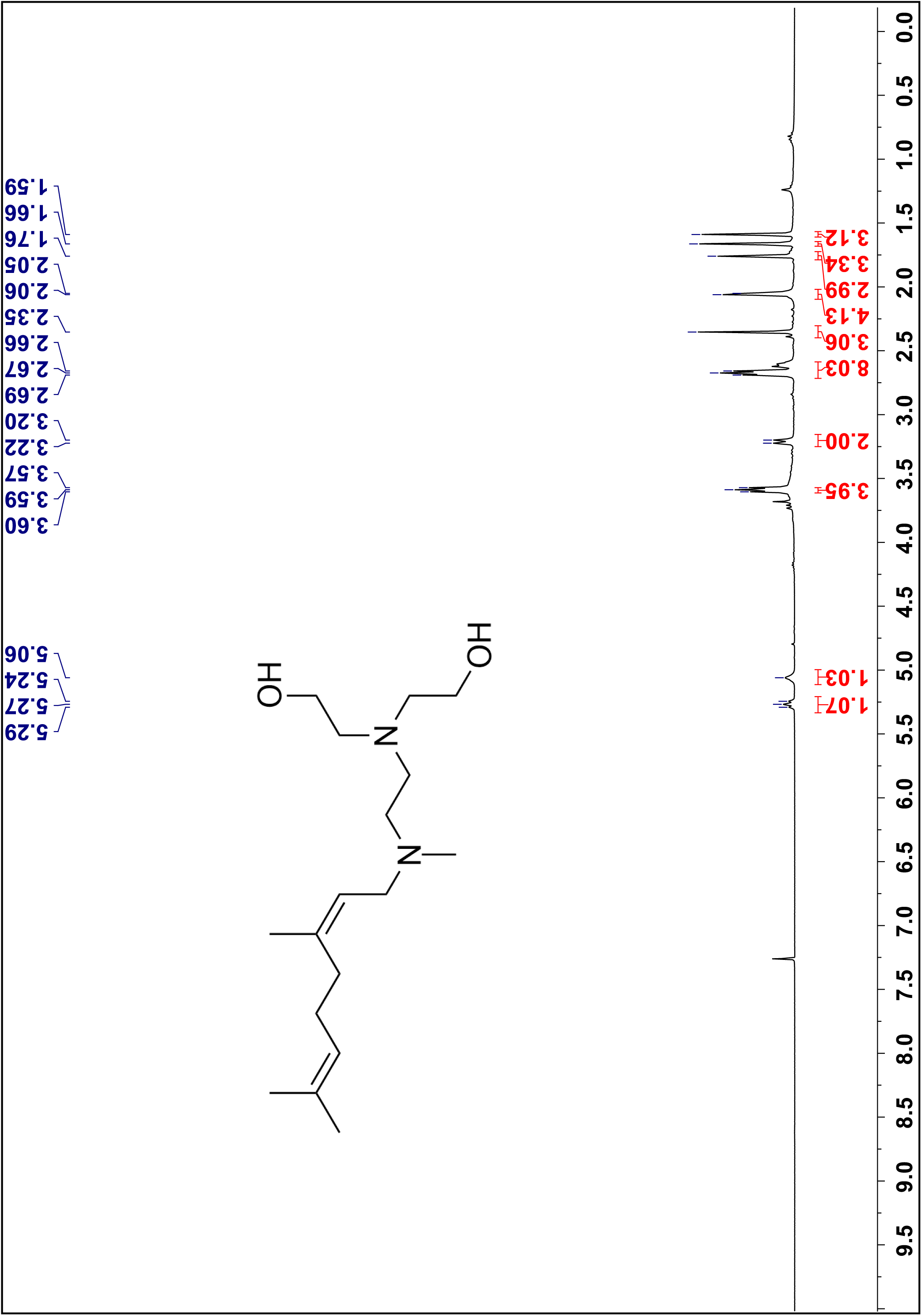
¹H NMR spectra of compound **5a** in CDCl3, 300 MHz.

**Figure 35.**
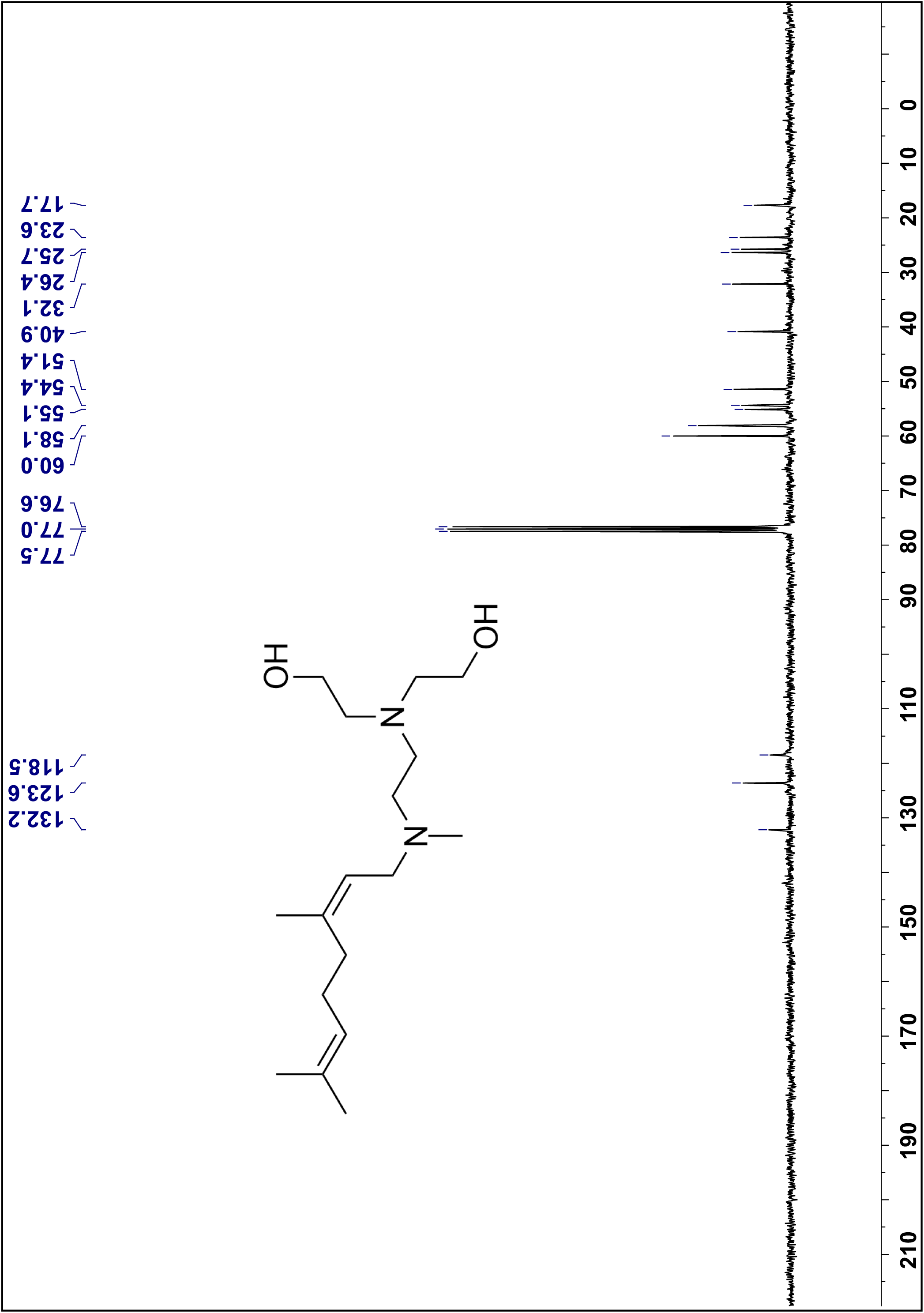
¹³C NMR spectra of compound **5a** in CDCl3, 75 MHz.

**Figure 36.**
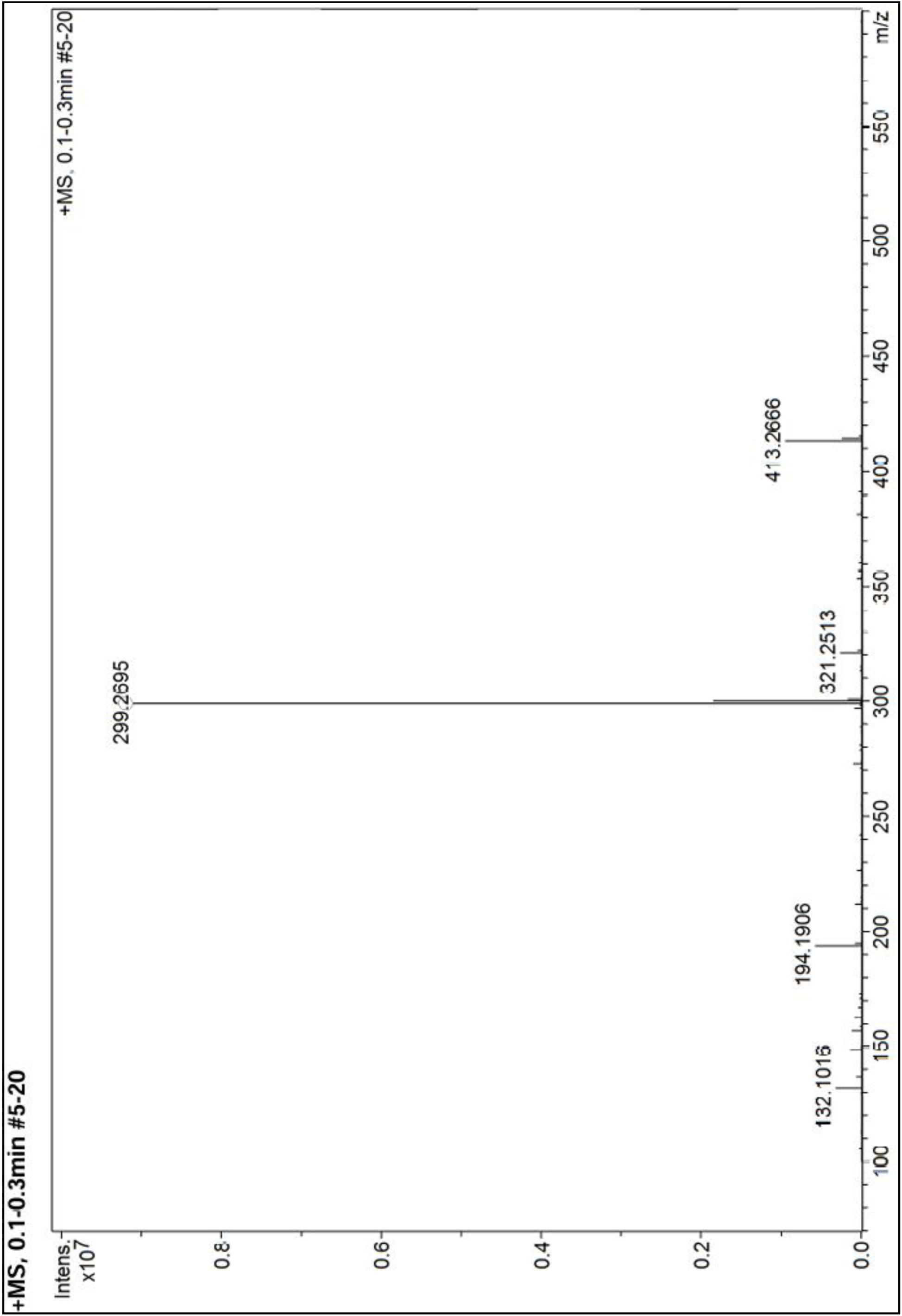
High Resolution Mass spectra of compound **5a**.

Compound 5b

**Figure 37.**
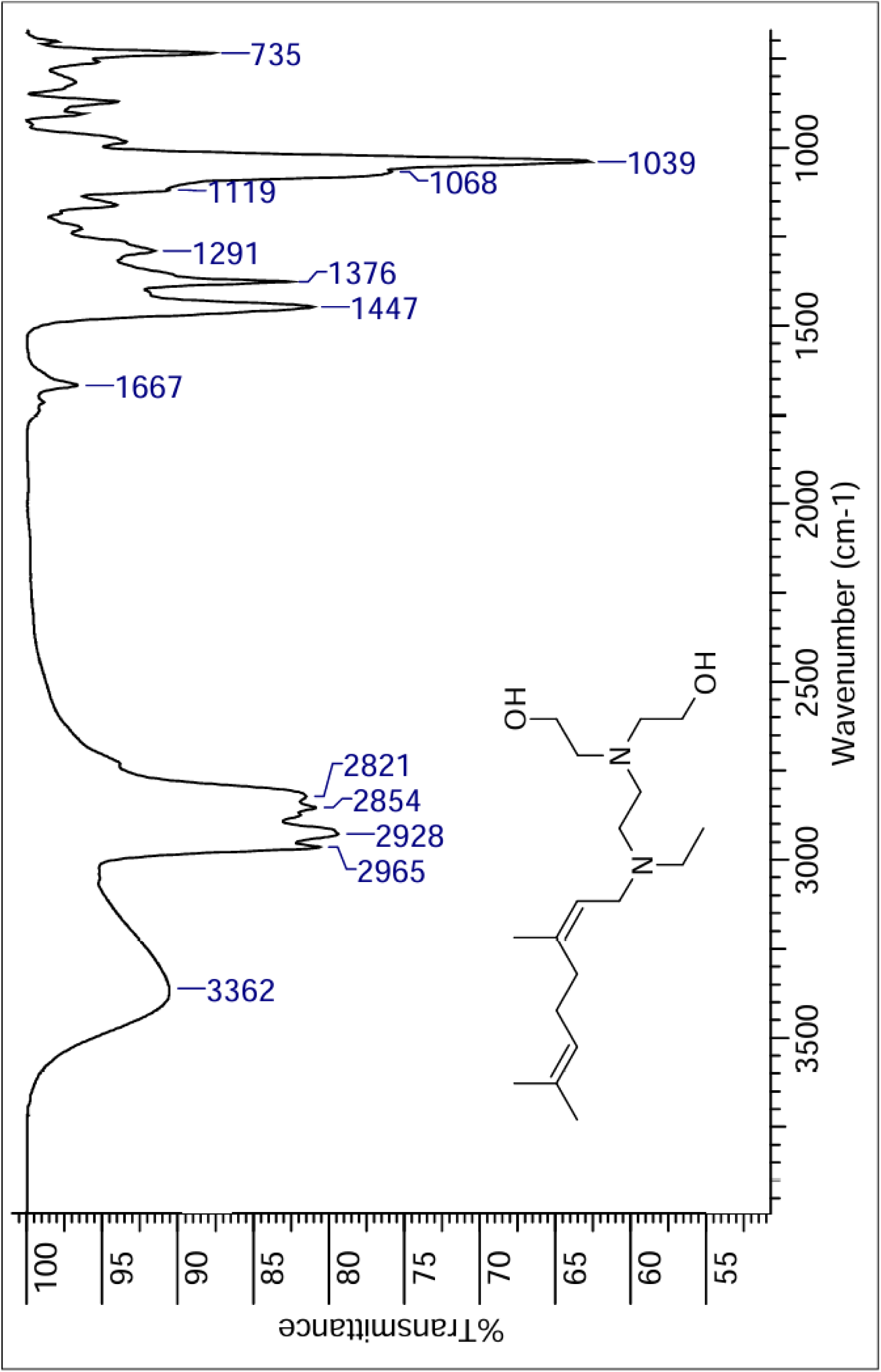
IR spectra of compound **5b**.

**Figure 38.**
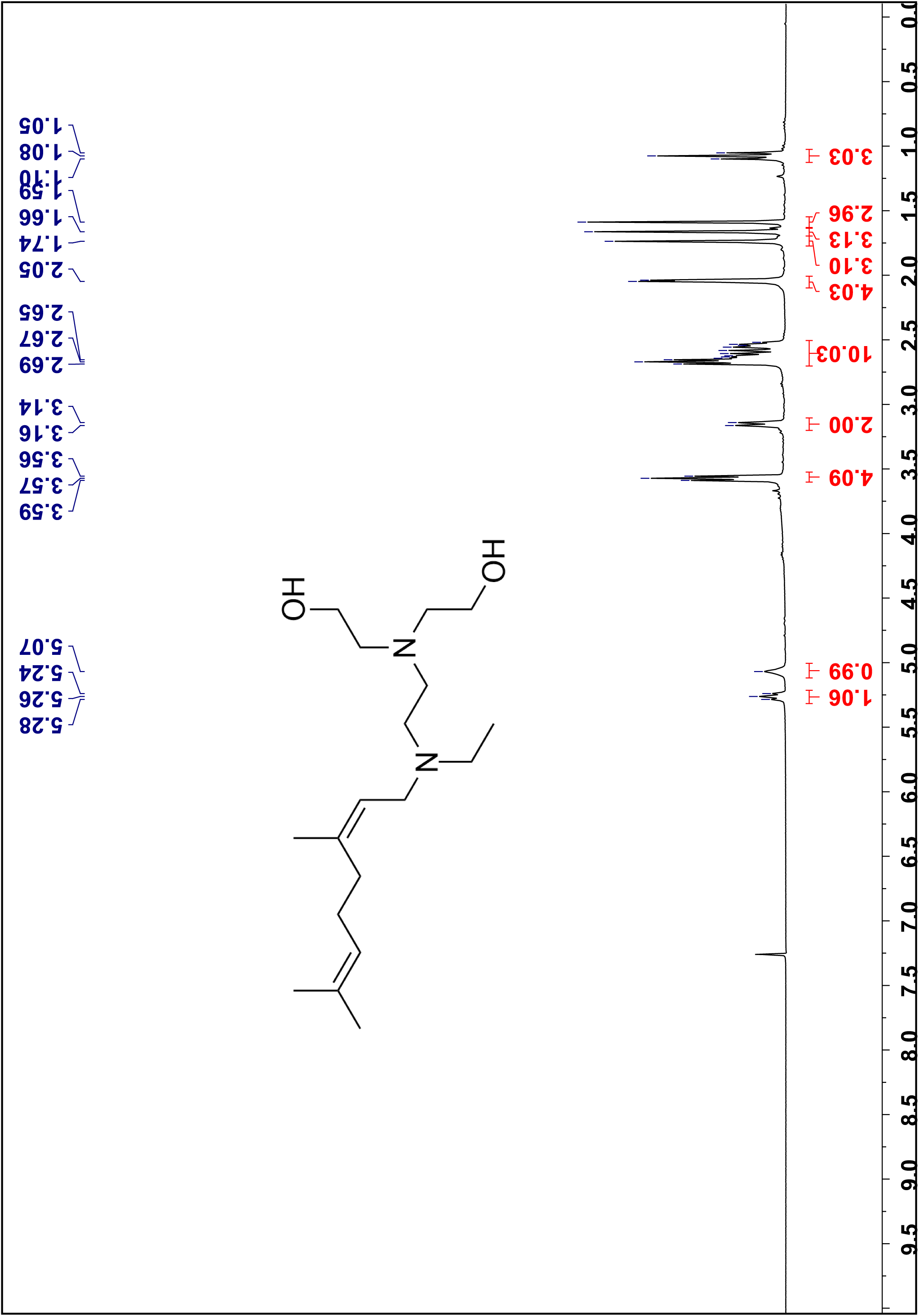
¹H NMR spectra of compound **5b** in CDCl3, 300 MHz.

**Figure 39.**
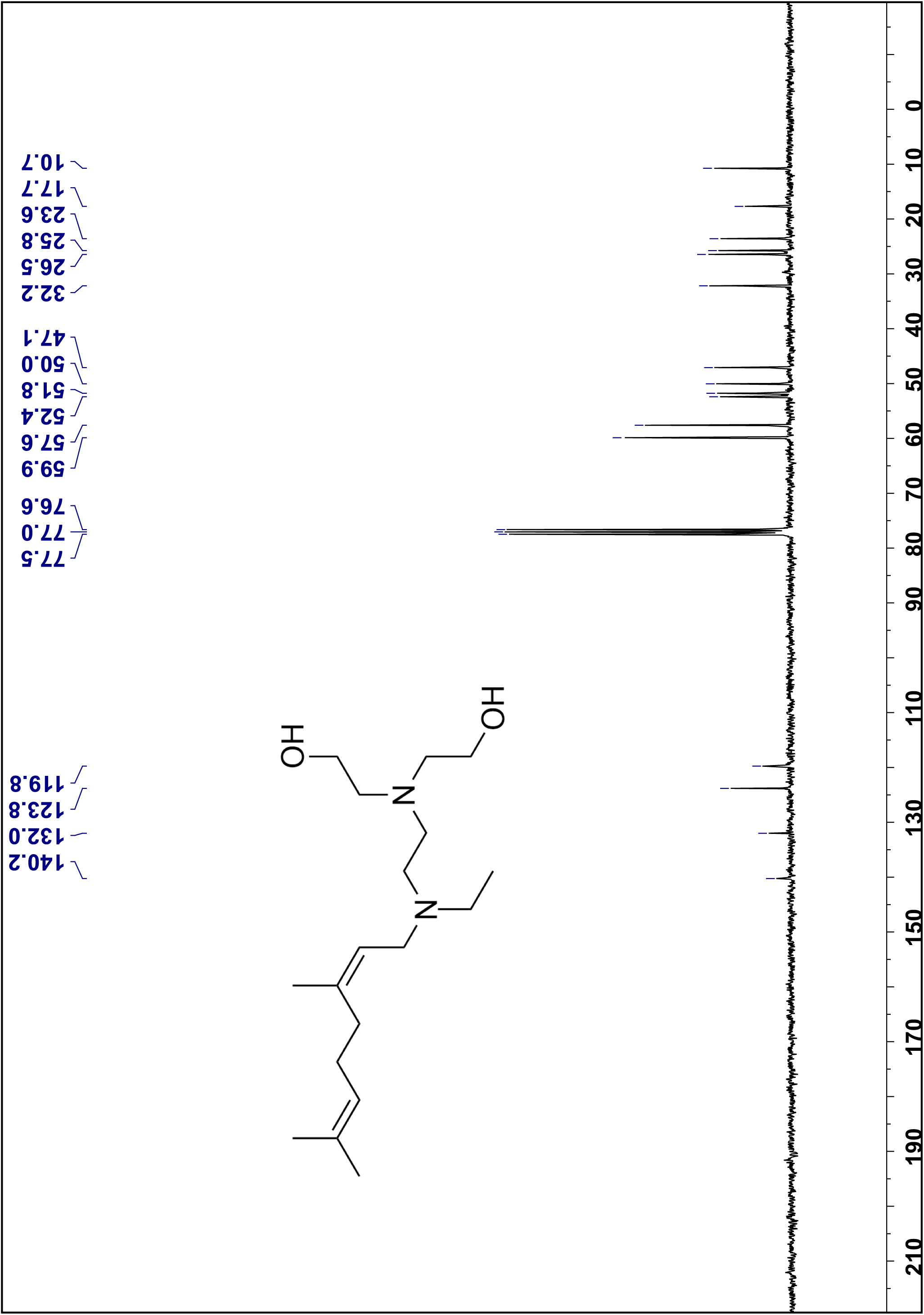
¹³C NMR spectra of compound **5b** in CDCl3, 75 MHz.

**Figure 40.**
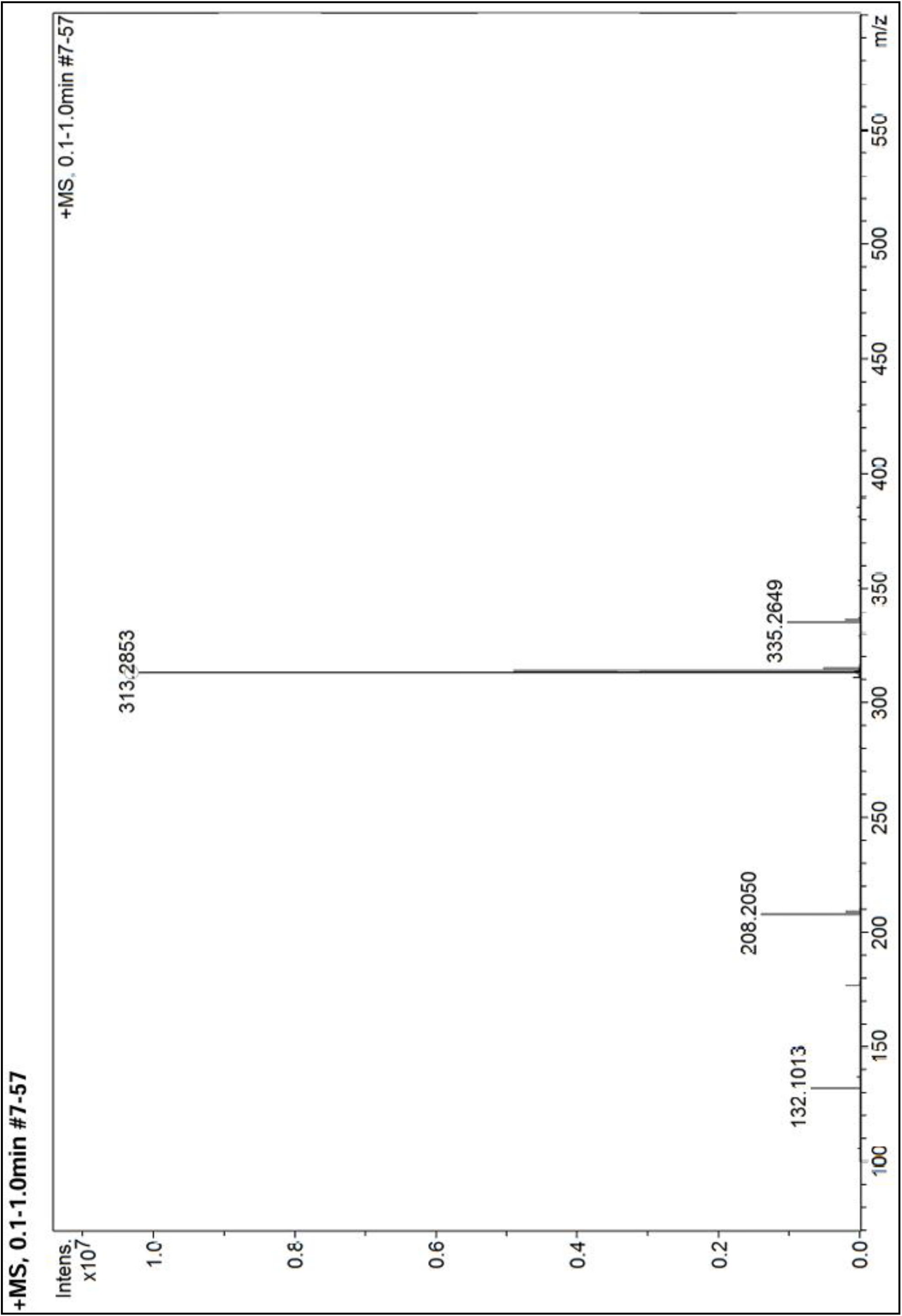
High Resolution Mass spectra of compound **5b**.

## Notes

### Competing Interest Statement

The authors have declared no competing interest.

